# Evolutionary rates provide genomic insights into convergent and lineage-specific evolution in carnivorous plants

**DOI:** 10.64898/2025.12.17.694974

**Authors:** Sukuan Liu, Evan S. Forsythe, Daniel B. Sloan

## Abstract

Carnivorous plants (CPs) are striking examples of convergent evolution across ecological, morphological, and molecular scales, making them ideal systems for investigating genomic adaptation to extreme environments. While genomic studies have shown that gene family expansion and functional recruitment underlie key carnivorous traits, the effects of these transitions on rates of sequence evolution remain incompletely understood. Here, we present a comparative framework to examine evolutionary rate shifts across 30 carnivorous and non-carnivorous eudicots in over 12,000 gene families. After normalizing for genome-wide background effects, analysis of branch-lengths and evolutionary rate covariation (ERC) detected both lineage-specific and convergent acceleration in CPs for genes associated with stress response, stimulus signaling, and light perception. Accelerated genes often belong to expanded families and show relaxation of purifying selection pressure. Despite the limited number of specific genes or gene families showing convergent acceleration across CP lineages, we find strong convergence at the functional level. Distinct sets of accelerated genes in each CP lineage are enriched for similar functional pathways, particularly light and hormone responses. ERC network analysis identifies a stress- and stimulus-responsive gene community with lineage-specific acceleration in Sarraceniaceae species and moderate acceleration in other CP lineages. Hub genes in this network are also co-expressed in a distantly related non-CP model system, suggesting repeated co-option of an ancestral regulatory module for carnivorous functions. Together, our results show that convergent genomic rate accelerations in CPs primarily involve functionally equivalent or similar but lineage-distinct genes, shaped by gene duplication, relaxed selection, and co-option of stress-response networks.

**Significance Statement:** Investigations of carnivorous plant (CP) evolution over the last decade have produced a large amount of genomic data that has yet to be fully explored. Using existing genomes and transcriptomes from five independently evolved CP lineages and their non-carnivorous relatives, we compared how rapidly different genes change to identify both shared and lineage-specific shifts associated with carnivory. We show that CP lineages frequently speed up evolution in genes involved in similar biological functions, such as responses to stress, light, and hormones. Using evolutionary rate covariation analysis, we also identify groups of genes that appear to have evolved together and been repurposed for carnivory from ancestral stress- and stimulus-response functions. These findings complement earlier work on gene family expansion in CPs and demonstrate that examining patterns of evolutionary rate can reveal new insights into how complex traits like carnivory evolve.

## Introduction

Carnivorous plants (CPs) represent one of the most striking examples of convergent evolution, in which distantly related lineages have independently repurposed leaves into traps that attract, capture, and digest small-animal prey— followed by enzymatic breakdown and nutrient absorption—to offset nitrogen and phosphorus limitation in nutrient-poor environments (i.e., the carnivorous syndrome (Juniper et al., 1989; Sinnott-Armstrong et al., 2022)). Documented at least eleven times across six flowering plant orders (Fleischmann et al., 2018), these repeated origins position CPs as ideal systems for investigating convergent adaptation to extreme environments across multiple biological scales.

Ecologically, CPs show convergence not only in their carnivorous syndrome but also in their habitat preferences (Brewer & Schlauer, 2018). Across diverse geographic regions and evolutionary lineages, CPs are typically found in humid, high-light environments with nutrient-depleted soils (Brewer & Schlauer, 2018). Morphologically, similar trap structures and associated prey-capture mechanisms have evolved independently in multiple lineages (Fleischmann et al., 2018). For instance, sticky flypaper traps have independently arisen four times in Lamiales (independent emergence in *Byblis* and *Pinguicula*), Ericales (i.e. *Roridula*), and Caryophyllales (e.g. *Drosera*), whereas pitcher traps have evolved independently in *Nepenthes* (Caryophyllales) and Sarraceniaceae (Ericales) species as well as *Cephalotus follicularis* (Oxalidales) (Płachno & Muravnik, 2018; Thorogood et al., 2018).

At the molecular level, genomic studies of several CP taxa in Lamiales (Lentibulariaceae: *Pinguicula spp.* (Fleck et al., 2025), *Utricularia spp.* (Ibarra-Laclette et al., 2011; Ibarra-Laclette et al., 2013; Silva et al., 2016; Silva et al., 2019), and *Genlisea spp.* (Wicke et al., 2014)) and Caryophyllales (Droseraceae: *Drosera spatulata*, *Dionaea muscipula*, and *Aldrovanda vesiculosa* (de Vries & de Vries, 2020; Nevill et al., 2019; Palfalvi et al., 2020; Robledillo et al., 2025); Nepenthaceae: *Nepenthes spp.* (Gruzdev et al., 2019; Robledillo et al., 2025; Saul et al., 2023)) as well as the monotypic *C. follicularis* (Cao et al., 2019; Fukushima et al., 2017) have revealed convergent genome shifts associated with the evolution of carnivory. Tandem and whole-genome duplications appear to play central roles in expanding gene families involved in trap development and nutrient processing, with defense-related genes often co-opted or sub-functionalized for carnivorous functions such as prey digestion and nutrient uptake (Palfalvi et al., 2020; Renner et al., 2018). Convergent gene losses and pseudogenization are also widespread (e.g. plastid *ndh* genes)—consistent with broader genomic shifts linked to the carnivorous lifestyle (de Vries & de Vries, 2020). These studies, which have employed metabolomic (Dávila-Lara et al., 2020; Hotti et al., 2017; Lathika et al., 2025), proteomic (Hatano & Hamada, 2012; Takahashi et al., 2005; Zakaria et al., 2018), transcriptomic (Saganová et al., 2018; Srivastava et al., 2011; Wakatake & Fukushima, 2025), and comparative genomic (Fleck et al., 2025; Fukushima et al., 2017; Palfalvi et al., 2020; Robledillo et al., 2025) approaches, have generated critical evolutionary insights and a large body of CP genomic resources.

Together, these advances provide a strong foundation, but several key gaps in our understanding of CP genome evolution persist. First, several carnivorous lineages, such as Sarraceniaceae and *Byblis*, remain underrepresented in previous genome-wide comparative analyses, limiting our ability to generalize findings across the full evolutionary diversity of CPs. Second, changes in rates of sequence evolution could be indicative of selection pressures (e.g. purifying vs. positive and intensified vs. relaxed selection) associated with the adaptation to carnivory (Wicke et al., 2014). However, few studies have adopted phylogenomically broad, rate-based frameworks to systematically evaluate both convergent and lineage-specific genomic changes. Finally, although carnivory represents the repeated evolution of complex, integrated traits, it remains unknown whether independent CP lineages might have convergently co-opted similar ancestral gene networks for carnivorous functions. Under a phylogenomically broad, rate-based framework that samples independently evolved CP lineages and their non-CP relatives, this question can be addressed using evolutionary rate covariation (ERC) analysis (Clark et al., 2012). ERC quantifies coordinated substitution-rate changes between genes across lineages, reflecting the tendency of genes in the same pathways or functional modules to experience shared selective pressures and thus correlated rate shifts. Moreover, ERC detects not only direct physical interactions between protein pairs but also shared functionality and co-expression (Clark et al., 2012; Forsythe et al., 2021; Gatts et al., 2024; Little et al., 2024), allowing existing genomic resources to be used to infer putative gene networks underlying carnivorous traits. Thus, ERC provides a genome-scale framework for identifying potentially co-evolving gene networks and testing whether CP lineages have repeatedly recruited similar molecular pathways during the evolution of carnivory.

In this study, we address these knowledge gaps by establishing a broad, rate-based comparative framework to investigate the genomic underpinnings of carnivory across independently evolved carnivorous angiosperm lineages. By incorporating available genomic data from well-studied CP species as well as species from previously under-sampled lineages (Figure 1, Supplementary Table A), we systematically compare evolutionary rate shifts between CPs and non-CPs across thousands of gene families. After normalizing each branch to its genome-wide mean (accounting for both lineage-specific rate differences and branch-specific divergence-time variation) we identified, in each CP species, gene families that were relatively accelerated or conserved relative to non-CPs. Leveraging ERC analysis via the ERCnet pipeline (Forsythe et al., 2025), we identify a convergently accelerated, co-evolving gene network community potentially repurposed for trap development, stress response, and digestive functions. Finally, we integrate our findings with gene expression data from CPs and non-CPs to assess the functional and regulatory coherence of these gene modules, illuminating how deeply conserved stress-responsive networks may have been repeatedly co-opted for carnivorous adaptations.

**Figure 1.**
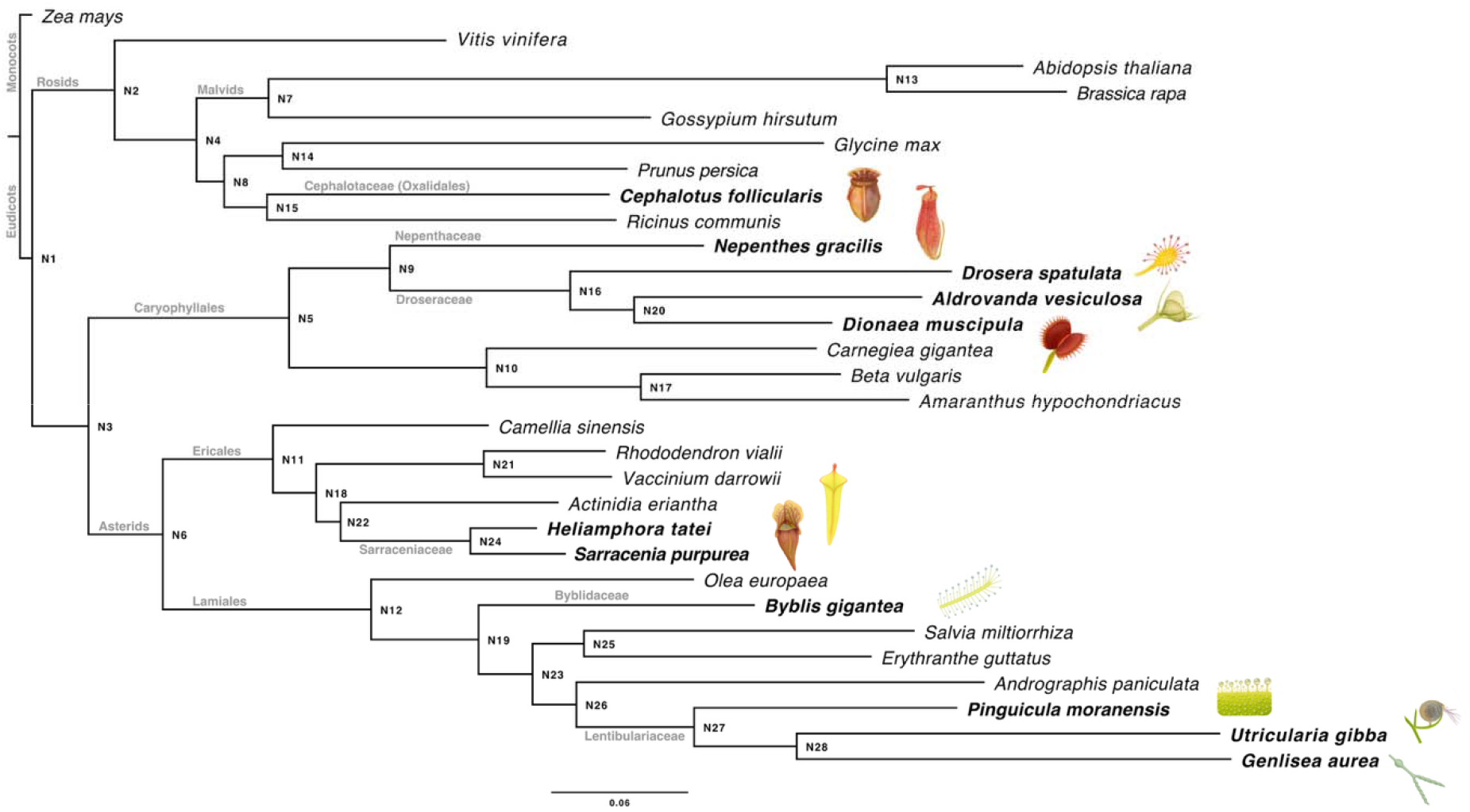
**Phylogeny of sampled taxa**, with carnivorous lineages in bold. Internal nodes are labeled, and branch lengths represent genome-wide mean branch lengths. Scale bar: mean substitutions per site. Stylized drawings illustrating the characteristic trap type of each carnivorous species are shown next to the corresponding CP species.

## Results

### Reduced BUSCO completeness in CPs reflects genome contraction and sequencing constraints

Protein-coding sequence (CDS) datasets from 30 CP and non-CP taxa revealed marked differences in completeness based on BUSCO (Benchmarking Universal Single-Copy Orthologs) assessments (Manni et al., 2021) (Supplementary Table B): genome-derived CDS sets were consistently more complete than transcriptome-derived sets. On average, genome-derived datasets contained 88.7% complete and 8.3% missing BUSCOs, whereas transcriptome-derived datasets averaged 72.2% complete and 20.6% missing. Carnivorous taxa— four of which are represented by transcriptomes—showed lower completeness than non-carnivorous taxa (complete: 76.7% vs. 92.3%; missing: 17.5% vs. 5.5%). Lower completeness in several carnivorous taxa likely reflects both technical factors (limited tissue sampling and shallow RNA-seq coverage; e.g., *B. gigantea*, *P. moranensis*) and biological factors consistent with convergent as well as lineage-specific genome reduction reported in CPs (e.g., Lentibulariaceae: *P. gigantea* (Fleck et al., 2025), *Genlisea aurea* (Leushkin et al., 2013), *Utricularia gibba and U. reniformis* (Carretero-Paulet et al., 2015; Ibarra-Laclette et al., 2013; Silva et al., 2019); Droseraceae: *A. vesiculosa*, *Di. muscipula*, *Dr. spatulata* (Nevill et al., 2019; Palfalvi et al., 2020)). A visualization of the workflow implemented in this study can be found in Supplementary Figure 25.

### Stringent HOG filtering reduces rate-estimate coverage but improves reliability for downstream rate comparison and network inference

OrthoFinder (Emms & Kelly, 2019) identified 43,632 (at eudicot level) hierarchical orthologous groups (HOGs) across the input dataset. Of these, 33,163 contained sequences from two or more taxa (including 2,272 shared across all 29 sampled eudicot species), whereas 10,469 were species-specific. Across all HOGs, the mean and median numbers of sequences per group were 19.3 and 10.0, respectively, and no HOG was strictly single-copy. Gene families were broadly shared among carnivorous lineages: 21,620 HOGs contained sequences from two or more of the 11 carnivorous taxa, and 3,260 contained sequences from all 11 (Supplementary Figure 31). We applied the ERCnet representation filters (≥15 species and ≤20 gene copies per species, with HOGs not meeting these requirements discarded in full), yielding 13,663 HOGs suitable for downstream analysis (Supplementary Table C). With this set of HOGs, we performed our phylogenomic analyses in multiple replicates, allowing us to assess the robustness of our evolutionary rate estimates (see Methods and Supplementary Figure 25). The cross-replicate consensus matrix used for all downstream rate analyses comprised 12,490 HOGs (Supplementary Figure 26).

Although these HOGs are broadly shared, comparable BL estimates obtained from our consensus matrix are far sparser across lineages after filtering (Supplementary Figure 32). Each CP lineage carries BL estimates for 2,579– 8,420 HOGs in the consensus matrix, but of the 12,059 HOGs with an estimate in at least one CP lineage, only 114 carry estimates on all 11 CP terminal branches simultaneously, and 59% are present in five or fewer CP lineages (Supplementary Figures 1 and 30). This collapse—from 3,260 fully shared HOGs (Supplementary Figure 31) to 114 with comparable estimates—reflects the cumulative effect of the analysis filters (≥ 15-species representation, ≤ 20 copies per species, ≤ 5% gap in multiple sequence alignment, and the requirement of a replicate-consistent BL estimate on each branch), together with the deep, phylogenetically dispersed placement of carnivorous lineages and the incomplete CDS/transcriptome assemblies of several CP taxa (lower BUSCO completeness; see above).

On average, fewer than half of the HOGs present in each lineage retained a usable BL estimate across CPs (mean 40%; range 21–58%; Supplementary Figure 32), and this recovery rate varied systematically among lineages. The pitcher plants (*Nepenthes*, *Cephalotus*, *Sarracenia*, and *Heliamphora*) and *Drosera* retained the most estimates (47– 58%), while the snap-trap species of Droseraceae (*Dionaea*, *Aldrovanda*) and Lentibulariaceae (*Pinguicula*, *Utricularia*, *Genlisea*) retained the fewest (21–35%). This gradient likely reflects genome reduction acting not only at the gene level, through lineage-specific gene loss, but also at the sequence level: in the most reduced genomes, many protein-coding sequences are truncated and fall below the minimum alignment-length and coverage thresholds required to enter gene-tree inference, and are therefore excluded before BL estimation. Although this filtering reduced the number of estimates available for cross-lineage rate comparison among the independently evolved CP lineages, the trade-off is necessary: it removes noise arising from unreliable branch-length estimates in poorly resolved, weakly informative, or highly paralogous gene families, where gene-tree inference and branch-length reconciliation—which averages across paralogous copies—are most error-prone (Salichos & Rokas, 2013; Sarton-Lohéac et al., 2025). Functions over-represented among estimable HOGs (relative to all sequence-bearing HOGs from OrthoFinder outputs) were restricted to conserved core-cellular and housekeeping categories and showed no overlap with the accelerated-gene enrichments (0 of 21 terms), with only two marginal overlaps with conserved-gene enrichments (2 of 21; Supplementary Table H), indicating that estimate availability does not bias the dataset toward the carnivory-associated accelerated functions central to our conclusions. Notably, these enrichments were largely confined to the genome-reduced Lentibulariaceae (*Pinguicula*, *Utricularia*, and *Genlisea*), which accounted for most over-represented terms and both conserved-gene overlaps—consistent with the lineage-specific genome reduction discussed above.

Because requiring HOGs shared across all lineages would discard the vast majority of the data, we did not restrict comparisons to shared HOGs; instead, evolutionary rate was assessed per HOG × branch as a Z-score relative to the non-CP distribution (see Methods), allowing each CP lineage to contribute its full set of estimable HOGs. Importantly, because each Z-score is derived from per-branch normalized BLs—which already correct for lineage-specific genome-wide rate variation and for the divergence time of each branch—and is then standardized against the distribution of the same HOG across non-carnivorous taxa, it additionally accounts for HOG-specific rate variation. Each Z-score thus represents the relative rate shift of a given HOG on a CP terminal branch, measured against all non-CP taxa sampled in this study.

### Modest rate shifts in CPs, with lineage-specific asymmetry

We used the ERCnet pipeline to generate BL estimates (BL consensus matrix from 5 replicates, see Methods) for each HOG (Supplementary Table C, Supplementary Figure 26). To make these BL estimates comparable across lineages, we then normalized them to account for lineage-specific rate differences and variation in branch durations (Supplementary Figure 3, see Methods). To quantify relative rate shifts between CPs and non-CPs and to account for mutational-rate variation among HOGs, we further transformed the normalized BLs into Z-scores corresponding to the extent of rate variation between CP and non-CP lineages (DeVore, 2017). We also separately applied a log transformation to the BL estimates to mitigate heteroskedasticity in the dataset (Little et al., 2025) (see Methods for details).

Our BL Z-score analysis indicates that variable numbers of genes across CP lineages (302 to 2,155 per lineage, out of 2,579–8,418 HOGs tested; Supplementary Table D) show rate shifts relative to non-CPs (|Z| ≥ 1 in both the normalized and log-normalized datasets). Comparison of normalized BLs and Z-scores revealed a generally positive but highly dispersed relationship between these two measures (Supplementary Figure 4 & 5). This pattern indicates that, within the genome of a given CP lineage, genes that evolve faster than other genes in the same genome (higher normalized BLs; within-genome comparison) also tend to evolve faster than their homologs in non-CPs (higher Z-scores; across-phylogeny comparison), although the wide scatter in this relationship shows that normalized BL alone does not fully determine whether a given gene in CP species will exhibit a detectable rate shift relative to non-CPs. Importantly, the classifications of genes as accelerated or conserved in CPs are based on BL estimates that have been normalized to account for genome-wide rate differences across species.

We also examined the non-CP reference set used to compute Z-scores, which comprised a median of 25 non-CP branches per HOG (range 3–39; Supplementary Table D). We found that accelerated and conserved calls were not concentrated among HOGs with few reference branches (HOGs with fewer than 10 non-CP branches accounted for 2.7% of tests and 2.3% of calls), suggesting that applying a single threshold (|Z| ≥ 1) across HOGs with different reference sizes did not bias our classifications. Because the non-CP reference branches share evolutionary history (e.g., a gene with relaxed constraint in a common ancestor may evolve rapidly in all of its descendant lineages), we also estimated phylogenetic signal in normalized BLs using Pagel’s λ, fitted separately for each HOG across all branches of the species tree (see Methods). Phylogenetic signal was weak (median λ = 0.04 and 0.00 for normalized and log-normalized BLs, respectively) and significant for fewer than 1% of HOGs after FDR correction. We therefore retained this Z-score framework rather than phylogenetically explicit approaches such as RERconverge (Kowalczyk et al., 2019), which require single-copy orthologs whose gene trees follow the species-tree topology and would exclude the expanded, multi-copy gene families central to our analyses.

We found that rate-shift patterns were lineage-specific. In Lentibulariaceae, accelerated HOGs exceeded conserved HOGs by approximately two- to threefold—*P. moranensis* (synonym = *P. caudata*, 7.9% vs. 2.6%), *U. gibba* (8.1% vs. 3.6%), and *G. aurea* (8.5% vs. 4.5%). Several species showed few rate-shifted genes overall (e.g., *C. follicularis* 2.0%/2.6%; *N. gracilis* 4.1%/3.2%; *B. gigantea* 4.7%/3.5%), whereas two Sarraceniaceae species exhibited the highest fractions, with similar proportions of accelerated (*H. tatei* 14.0%, *S. purpurea* 11.8%) as well as conserved (*H. tatei* 14.5%, *S. purpurea* 11.3%) HOGs.

Because four CP lineages (*H. tatei*, *S. purpurea*, *B. gigantea*, and *P. moranensis*) are represented by transcriptome-derived datasets, we examined whether data quality could account for this variation. In a regression including both variables, the fraction of HOGs with |Z| ≥ 1 per CP lineage was not associated with BUSCO completeness (P = 0.80) or mean alignment length (P = 0.57), nor did it differ by data source (transcriptome- vs. genome-derived; P = 0.53); *B. gigantea* and *P. moranensis* showed among the lowest fractions (0.08 and 0.11). Instead, the two Sarraceniaceae species have the shortest terminal branches among the sampled CPs (genome-wide mean raw BL of 0.028 and 0.042 substitutions per site, compared with 0.087–0.191 in the other nine CP lineages), and terminal branch length explained 70% of the variation among lineages when included in the regression (P = 0.006). On such short branches, each gene accumulates relatively few substitutions, so normalized BLs are expected to be less precise, inflating both accelerated and conserved calls. The elevated and similar fractions of accelerated and conserved HOGs in Sarraceniaceae therefore likely reflect, at least in part, noise captured by the normalization of short branches rather than stronger lineage-specific rate shifts.

### Accelerated gene families are generally expanded in CP genomes

Although our ERC-based framework was not specifically designed to detect gene family expansion/contraction and may be affected by stringent sequence-filtering steps and the incomplete nature of available CP transcriptome assemblies, it nevertheless provides a useful approximation by counting homolog numbers per species in reconciled gene trees. We compared paralog counts among accelerated, conserved, and all other HOGs across CP species (Supplementary Figure 6, Supplementary Table E).

Across most CP lineages, accelerated genes consistently exhibited higher copy numbers than both conserved genes and the genomic background (i.e., genes not annotated as accelerated or conserved). In *A. vesiculosa*, *C. follicularis*, *Di. muscipula*, *H. tatei*, *N. gracilis*, *S. purpurea*, and *U. gibba*, accelerated genes contained significantly more paralogs than conserved genes, as supported by all three statistical tests (Welch’s t-test, Mann–Whitney U, and Mood’s median). By contrast, *B. gigantea*, *Dr. spatulata*, *G. aurea*, and *P. moranensis* did not exhibit significant differences in gene copy number between accelerated and conserved sets, although accelerated genes still displayed slightly higher mean copy numbers in all cases. Even in these lineages, accelerated genes often exceeded background levels (i.e., significantly more gene copies in several accelerated gene sets than in “all other” HOGs), indicating a general trend for faster-evolving genes to occur in expanded families.

Conserved genes, in contrast, rarely differed significantly from the genome-wide background. When differences were detected (e.g., in *S. purpurea* and *N. gracilis*), they reflected reduced copy numbers, suggesting that conserved genes are typically associated with copy number stability or mild contraction rather than expansion (Supplementary Figure 6, Supplementary Table E).

Overall, these results strengthen the link between evolutionary rate acceleration and gene family expansion across multiple independent origins of carnivory. Such expansions may have facilitated the subfunctionalization or neofunctionalization of duplicated genes into specialized roles underpinning carnivorous adaptations.

### Accelerated and conserved genes associated with distinct selective pressures and intensities

To test whether faster- and slower-evolving genes experience different selective pressures, we estimated the mean nonsynonymous-to-synonymous substitution rate ratio (ω) for every HOG across terminal branches of CP lineages (Supplementary Data). Consistent with expectations that higher Z-scores represent more non-synonymous substitution in CP species than non-CPs for a given HOG, our nonparametric Mann–Whitney U tests revealed that all contrasts between accelerated and conserved gene sets were significant after false discovery rate (FDR) correction, with median ω consistently higher in accelerated than conserved gene sets. Parametric Welch’s t-tests yielded the same direction of effect, confirming that accelerated HOGs exhibit higher mean ω values than conserved HOGs.

Across all CP species and rate categories, mean ω values (0.32–2.01) exceeded the corresponding medians (0.11– 0.34), indicating strongly right-skewed distributions (Figure 2A, Supplementary Figure 7). This pattern suggests that most genes evolve under purifying selection, whereas a smaller subset experiences elevated ω consistent with positive selection and faster evolution. Mean ω across all rate categories was significantly higher in Droseraceae than in other CP lineages, implying lineage-specific differences in selective pressure.

**Figure 2.**
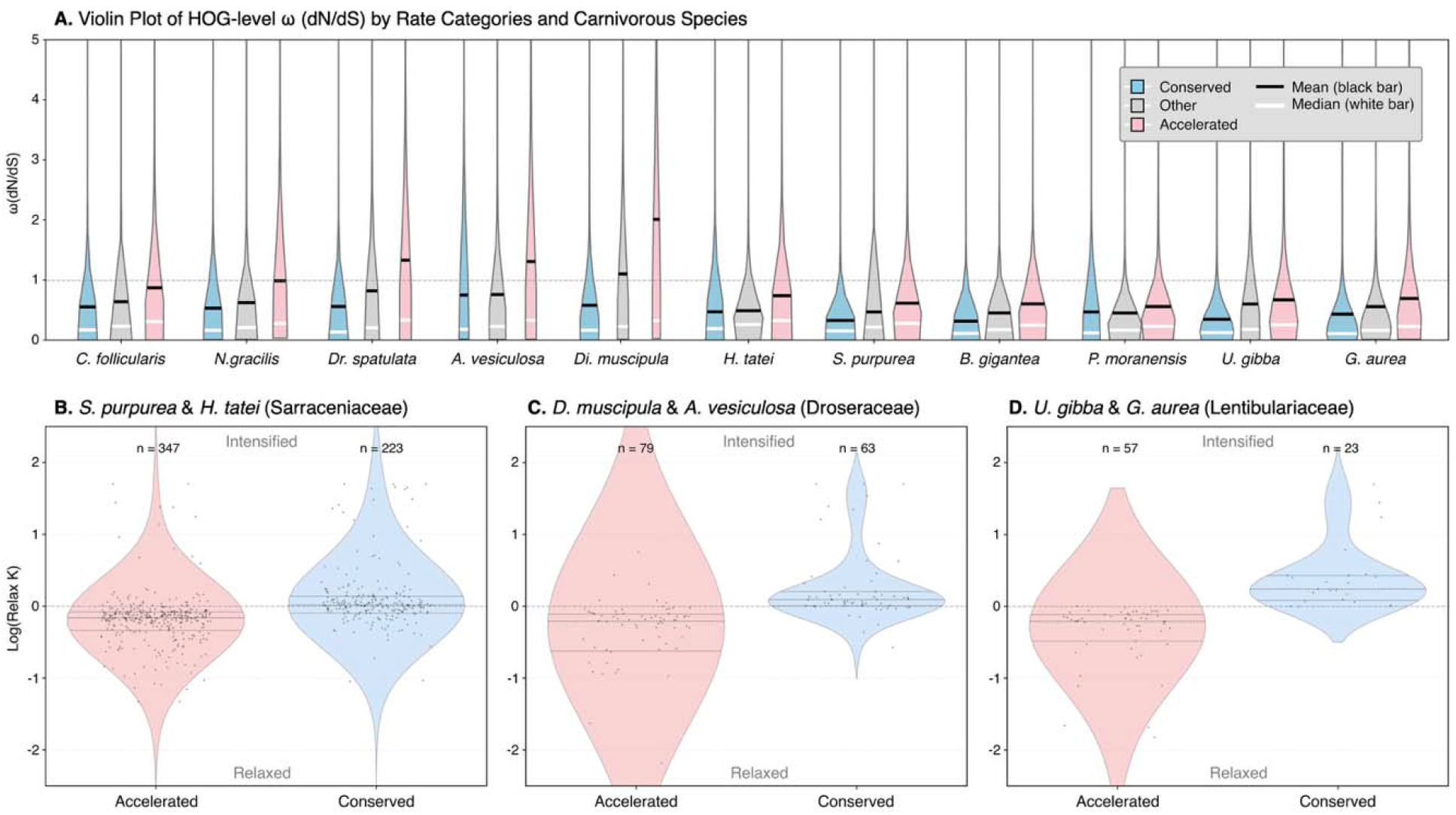
Evolutionary rate variation and selection-intensity across CP lineages. A. Distribution of HOG-level mean ω (dN/dS, estimated with aBSREL) across 11 CP species and three rate categories. For each species, violins show the distribution of mean ω across hierarchical orthogroups (HOGs). Categories are Accelerated (pink), Conserved (blue), and Other (gray, HOGs without significant rate shifts). Horizontal bars indicate the mean (black) and median (white) ω per distribution. The y-axis is truncated at 5 to emphasize the main range of values. B–D. Selection-intensity (log-transformed RELAX K) by rate category for homologous HOGs shared between representative CP lineages. Violin plots show the distribution of K for Accelerated and Conserved genes in (B) Sarraceniaceae (*S. purpurea, H. tatei*), (C) Droseraceae (*Di. muscipula, A. vesiculosa*), and (D) Lentibulariaceae (*U. gibba, G. aurea*). Points represent individual HOGs; dashed lines mark K = 1 (no net change in selection intensity). Horizontal dotted lines indicate medians and interquartile ranges, and n denotes the number of HOGs per category.

To further compare selection regimes, we estimated the selection-intensity parameter K with the RELAX method (Wertheim et al., 2015) for HOGs shared between closely-related sister taxa within Sarraceniaceae, Droseraceae, and Lentibulariaceae (Figure 2B, Supplementary Figure 8). Only HOGs showing consistent acceleration or conservation in both sister taxa were retained for this analysis (i.e., a HOG was considered “accelerated” or “conserved” only if classified identically in both taxa). Strikingly, in all three lineages, accelerated genes tended to exhibit relaxed selection (K < 1), whereas conserved genes tended to experience intensified selection (K > 1).

Further examination of the RELAX outputs suggested that the intensified selection observed in conserved gene sets was primarily driven by the intensification of purifying selection, rather than that of positive selection. In these cases, the inferred ω category corresponding to positive selection (ω) was typically associated with a very small proportion of sites (e.g. < 0.1%). By contrast, the ω categories representing purifying selection (ω) were generally inferred to be close to 0, encompassing more than 70% of codons in the aligned sequences. Together, these results indicate that accelerated and conserved genes have experienced systematically distinct selection intensities during the diversification of CPs and that the strength of purifying selection may be the primary determinant of variation in rates.

### Convergent acceleration of genes involved in stimulus and stress responses

We performed Gene Ontology (GO) enrichment analyses (Supplementary Data) for accelerated and conserved HOGs in each CP species (terminal branches). Across all lineages, we identified 370 unique Biological Process (BP) terms, 64 Cellular Component (CC) terms, and 142 Molecular Function (MF) terms that were significantly enriched in the accelerated and/or conserved HOGs from at least one carnivorous species. Of these, 158 BP, 27 CC, and 77 MF terms were enriched exclusively in accelerated HOGs, whereas 99 BP, 28 CC, and 37 MF terms were enriched exclusively in conserved HOGs. A substantial number of terms—113 BP, 9 CC, and 28 MF—were shared between the two categories.

The most frequently enriched terms across all three GO categories— BP (e.g., biosynthetic process, primary metabolic process, cellular component organization or biogenesis, small-molecule metabolic process, localization), MF (e.g., catalytic activity, hydrolase activity), and CC (e.g., protein-containing complex, catalytic complex)—were either shared between accelerated and conserved sets across many lineages or exhibited lineage-specific acceleration alongside relative evolutionary constraint in all other lineages. This pattern suggests that core metabolic, regulatory, and structural pathways harbor both rapidly evolving and deeply conserved components.

Comparing enriched GO terms in the accelerated gene sets across carnivorous lineages revealed several signals of convergence in both BP and MF. Accelerated BP terms were consistently enriched for stimulus-response pathways, including response to light stimulus in most carnivorous lineages (*A. vesiculosa*, *C. follicularis*, *Di. muscipula*, *N. gracilis*, *P. moranensis*, *S. purpurea*, *U. gibba*) and response to chemical stimulus in the pitcher plants (*H. tatei*, *N. gracilis*, *S. purpurea*). Response to stress was also repeatedly enriched among accelerated genes in *C. follicularis*, *N. gracilis*, *Di. muscipula*, *S. purpurea*, *H. tatei*, and *U. gibba*. At the molecular-function level, catalytic activity acting on a protein was significantly enriched among accelerated genes in *N. gracilis*, *Di. muscipula*, *A. vesiculosa*, *B. gigantea*, and *P. moranensis*, suggesting convergent selection on protein-modifying enzymes, potentially including digestive hydrolases central to prey breakdown.

Despite this convergence in broad functional categories, relatively few specific genes exhibited convergent acceleration across independent CP lineages. Within the response to light stimulus process (Figure 3), 149 HOGs were associated with acceleration in at least one CP lineage, but only seven—PHYA (HOG0011752), ADO2 (HOG0017674), WAV2 (HOG0028790), MPK3 (HOG0010010), POB1 (HOG0017598), MET1 (HOG0029756), and BCDHβ1 (HOG0007744)—showed convergent acceleration in three or more CP lineages, with the rest showing more isolated lineage-specific acceleration patterns. These 149 accelerated HOGs associated with response to light stimulus exhibited diverse yet functionally interconnected roles spanning photoreception, signaling, transcriptional regulation, and photosynthetic adaptation. Based on their functional annotation, accelerated HOGs could be grouped into 5 major categories: (1) Photoreception and light-responsive transcriptional regulation (e.g., PHYA (Franklin et al., 2007), CRY1 (Lin et al., 1996), BBX21 (Xu et al., 2017), COL8 (Griffiths et al., 2003)), encompassing photoreceptors, circadian components, and downstream light-regulated transcription factors; (2) Polarity and morphogenesis genes (e.g., WAV2 (Mochizuki et al., 2005), HAT3/5 (Bou-Torrent et al., 2012; Liu et al., 2022)), which link light and auxin signaling to trap architecture; (3) Ubiquitination and proteasomal degradation (e.g., COP1 (Kim et al., 2017), POB1 (Orosa et al., 2017)), which mediate post-translational control of light responses; (4) Hormone and stress-response integrators (e.g., MPK3 (Pecher et al., 2014), connecting light perception with defense and hormonal crosstalk; and (5) Transport, metabolic, regulation and repair enzymes (e.g., BCDHβ1 (Fujiki et al., 2000; Peng et al., 2015), MET1 (Srikant et al., 2022), CML23 (Wang et al., 2025), HPR (Pracharoenwattana et al., 2010; Timm et al., 2010)), reflecting adjustments in redox balance, ion flux, and photorespiration. Together, these categories highlight a modular pattern of photoregulatory evolution in CPs, wherein distinct gene sets have repeatedly adapted to fulfill similar functional roles in light perception, signaling, and developmental integration.

**Figure 3.**
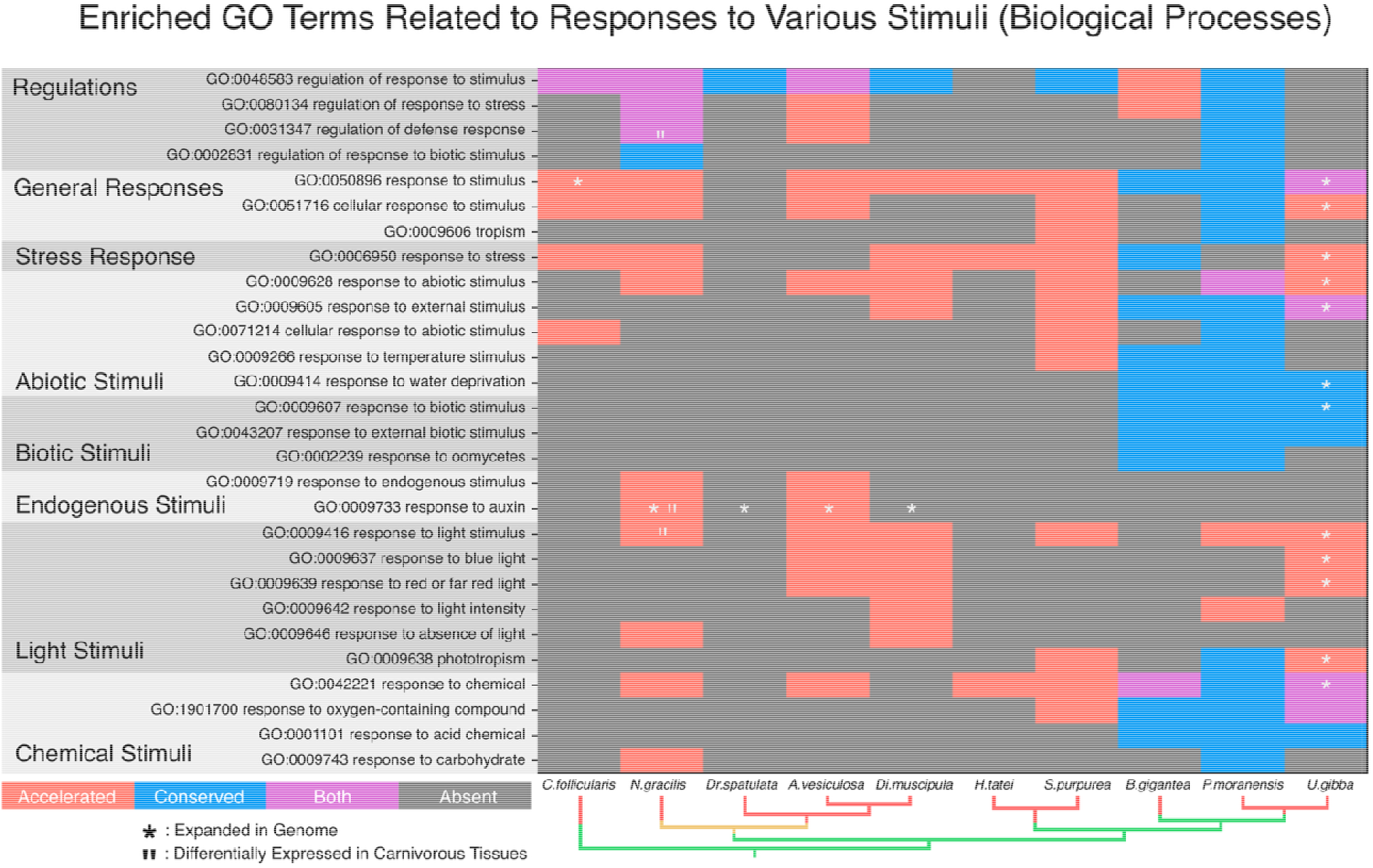
**Enriched GO terms (BP) related to responses to stress and various stimuli across six independent CP lineages**, shown as branches highlighted in red (alternative hypothesis of single emergence of carnivory in Caryophyllales is highlighted in yellow). Each cell represents the enrichment status of a GO term within a lineage. GO terms uniquely enriched among accelerated gene sets are shown in red, those uniquely enriched among conserved gene sets in blue, and those significantly enriched in both categories in purple. Terms not significantly enriched in either category are shown in grey. G. aurea is not shown because none of the GO terms displayed were significantly enriched in its accelerated or conserved gene sets. GO terms previously reported as enriched in expanded gene families or as differentially expressed in carnivorous tissues are

### Uniquely enriched ontology terms reflect lineage-specific rate shifts in CPs

In contrast to the convergent acceleration of genes involved in stimulus and stress responses, enriched GO terms in the conserved gene sets showed no consistent pattern of convergence across lineages. Instead, they appeared sporadically among species and were largely associated with essential cellular processes such as membrane transport, ion homeostasis, and ribosomal structure. These functions are critical for maintaining cellular integrity and are likely under strong purifying selection, resulting in evolutionary stability rather than repeated lineage-specific change.

We also detected lineage-specific GO term enrichments in closely related carnivorous taxa with similar or divergent trap types. In the sister pitcher-plant lineage Sarraceniaceae, lipid metabolic process was enriched among accelerated genes, whereas regulatory ncRNA-mediated gene silencing was enriched among conserved genes. In Droseraceae, functions associated with chloroplast organization were enriched in the accelerated genes of *Dr. spatulata* and *Di. muscipula*, while functions involved in chlorophyll catabolic processes were enriched in their conserved genes. Additionally, proteins localized to the apical plasma membrane were consistently found to evolve more slowly across all Droseraceae species.

### Concordance between evolutionary rate acceleration and functional recruitment in CPs

When comparing our lineage-specific enrichment results for accelerated and conserved genes with previous studies of gene family expansion and differential gene expression linked to carnivorous adaptations, we observed extensive overlap in functional categories (Figure 3). Many of the GO terms enriched in our accelerated gene sets were also reported as expanded or differentially expressed in carnivorous tissues in earlier single-species and lineage-level analyses of *C. follicularis* (Fukushima et al., 2017), *N. gracilis* (Saul et al., 2023), *S. purpurea* (Alabady et al., 2026; Albert et al., 2025), *Dr. spatulata*, *Di. muscipula*, *A. vesiculosa* (Palfalvi et al., 2020), and *U. gibba* (Carretero-Paulet et al., 2015). Therefore, to further understand the dynamics of accelerated evolution in CPs, we compared our enrichment results to prior analyses focused on CP genome expansion and tissue-specific gene expression.

In *C. follicularis*, *S. purpurea* and *U. gibba*, we found the broad stress-related term GO:0050896 (response to stimulus) was enriched among accelerated genes, and previously shown to have expanded in their genomes (Alabady et al., 2026; Carretero-Paulet et al., 2015; Fukushima et al., 2017). In *U. gibba*, we found additional terms such as GO:0009416 (response to light stimulus), GO:0042221 (response to chemical), and GO:0006950 (response to stress) showed parallel enrichment in both accelerated and expanded gene sets, as shown in (Carretero-Paulet et al., 2015). GO:0044042 (glucan metabolic process), is enriched both in the accelerated and expanded genes of *S. purpurea* (Alabady et al., 2026). In the snap-trap *Di. muscipula*, GO:0042545 (cell wall modification) was not only enriched among accelerated genes in present study, but also previously found to be expanded in its genome and upregulated in trap tissues—consistent with its proposed role in the rapid closure mechanism of the motile flytrap (Palfalvi et al., 2020). Similarly, GO:0071705 (nitrogen compound transport), GO:0016787 (hydrolase activity), and GO:0007165 (signal transduction) were enriched among accelerated genes of multiple Droseraceae species and were previously shown to have expanded at the base of the family prior to its subsequent diversification (Palfalvi et al., 2020). Notably, GO:0009733 (response to auxin)—enriched among accelerated genes in *N. gracilis* and *A. vesiculosa*—was also previously identified as an expanded function predating the independent origins of carnivory in Droseraceae and Nepenthaceae, as well as their non-sampled sister lineages Drosophyllaceae, *Triphyophyllum*, and Ancistrocladaceae (Palfalvi et al., 2020).

In *C. follicularis* and *N. gracilis*, where comprehensive expression-based enrichment datasets were available (Fukushima et al., 2017; Saul et al., 2023; Wakatake & Fukushima, 2025), we compared our rate-shift–based enrichment results with expression profiles and found partial but functionally meaningful overlap between our dataset and genes that are differentially regulated after prey capture. In *C. follicularis*, GO:0008652 (cellular amino acid biosynthetic process) and GO:1901605 (α-amino acid metabolic process) were found enriched among accelerated genes and known to be upregulated in pitcher digestive zones following prey capture (Fukushima et al., 2017; Wakatake & Fukushima, 2025). The conserved category included GO:0019843 (rRNA binding), which was similarly upregulated in multiple pitcher tissues in response to prey (Fukushima et al., 2017; Wakatake & Fukushima, 2025). Conversely, several GO terms enriched among accelerated genes were found to be downregulated in pitcher tissues after feeding, including GO:0009086 (methionine biosynthetic process) and GO:0009066 (aspartate family amino acid metabolic process) (Fukushima et al., 2017; Wakatake & Fukushima, 2025).

In *N. gracilis*, GO:0046873 (metal ion transmembrane transporter activity) and GO:0005262 (calcium channel activity)—terms enriched among conserved genes—were enriched among genes upregulated in the pitcher trap’s digestive zone after prey stimulation (Saul et al., 2023; Wakatake & Fukushima, 2025). In contrast, GO:0009416 (response to light stimulus), GO:0009733 (response to auxin), and GO:0031347 (regulation of defense response) were enriched among accelerated genes but corresponded to downregulated expression in the digestive and tendril tissues following prey induction (Saul et al., 2023; Wakatake & Fukushima, 2025).

Together, these findings indicate that the expression dynamics of accelerated and conserved genes associated with carnivory are complex and context-dependent. Some genes appear to be constitutively expressed to sustain core carnivorous functions, whereas others are differentially regulated in response to developmental transitions or prey stimuli (Saganová et al., 2018; Saul et al., 2023; Wakatake & Fukushima, 2025; Yilamujiang et al., 2016). Thus, evolutionary acceleration in carnivory-associated genes does not necessarily correspond to uniform upregulation, but instead reflects functional diversification and regulatory specialization across trap types, tissues, and ecological contexts.

### Network filtering improves ERC robustness

We inferred ERC networks from both normalized and log-normalized BLs to evaluate and mitigate heteroskedasticity in ERC inference and to assess consistency across variance treatments. The normalized network comprised 9,698 nodes (HOGs) and 30,993 edges (significant ERC between connected nodes), whereas the log-normalized network comprised 10,752 nodes and 90,835 edges. The two networks shared 8,848 nodes, corresponding to 91.2% of the normalized set and 82.3% of the log-normalized set (node Jaccard ≈ 0.76). Edge overlap was smaller— 5,508 shared edges, or 17.8% of normalized edges and 6.1% of log-normalized edges (edge Jaccard ≈ 0.05; Supplementary Figure 9). Because edges supported under both variance treatments are more likely to be robust to heteroskedasticity and upstream noise, we retained the intersection network for all downstream analyses. This conservative filter yields a core set of ERC relationships consistently recovered after variance stabilization, reducing sensitivity to treatment-specific false positives while preserving shared signal.

As an additional validation of the inferred ERC network, we computed Newman’s attribute assortativity coefficient (r), quantifying clustering by subcellular compartment (plastid, nucleus, mitochondrion). We observed significant positive assortativity—indicating that colocalized proteins tend to connect—with r = 0.089, P = 1 × 10 using the three compartments, and r = 0.031, P = 3 × 10 when including “Other” as a fourth category (Supplementary Figure 10). These results support that the ERC network preferentially links proteins localized in the same organelles. Additionally, we used these protein localization data to test whether proteins in each CP lineage experienced acceleration or conservation relative to non-CPs, with results shown in Supplementary Figures 11–13.

### Accelerated ERC community associated with lineage-specific acceleration in Sarraceniaceae

The inferred ERC network was divided into multiple components, with no edges connecting them (Supplementary Figure 14). The largest component contained 3,534 nodes and 4,305 edges, representing 65% of all nodes and 78% of all edges. The remainder comprised 705 smaller components: 234 with more than two nodes (965 nodes, 732 edges) and 471 two-node components (“single ERC hits”; 942 nodes, 471 edges). To characterize the structure of the largest interconnected component, we applied GLay clustering, which partitioned this giant component into 50 smaller network communities (median size = 62 nodes; range = 5–356).

We focused on communities containing at least 20 HOGs and tested whether any exhibited patterns of convergent or lineage-specific rate shifts across sampled CP lineages (Figure 4, Supplementary Table F). Our analysis revealed that most inferred ERC community clusters were not associated with significant rate shifts in CPs. This suggests that the bulk of the ERC signature we detect across the genome is driven by factors other than carnivory. Furthermore, in the few communities where significant shifts were detected, these were typically restricted to a single lineage (e.g., Communities 5, 7, 9) or showed mixed rate-shift patterns across multiple CP species (e.g., Communities 2, 11, 30).

**Figure 4.**
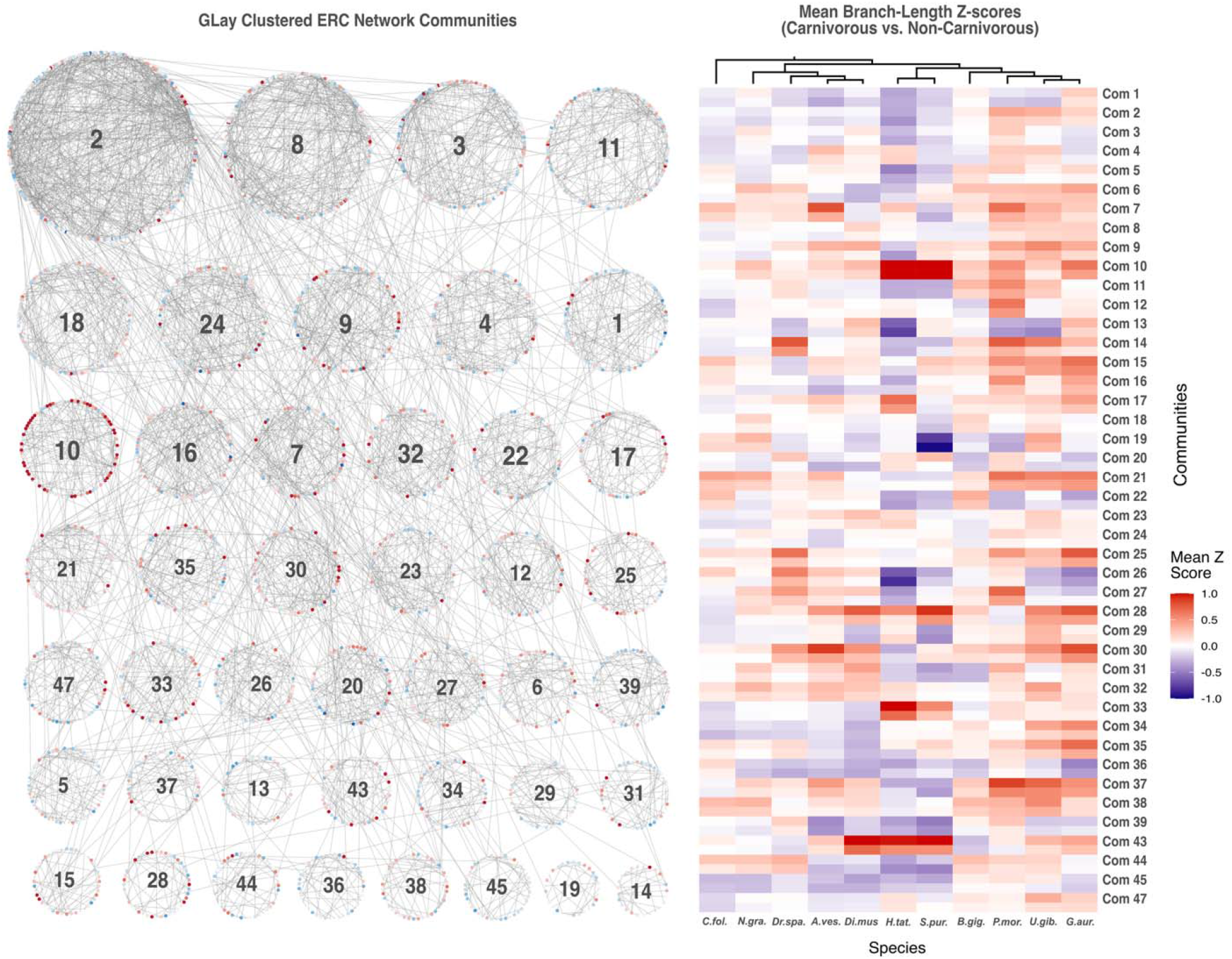
ERC network communities and lineage-specific branch-length shifts. **Left:** GLay-clustered communities of the shared ERC network (intersection of the normalized and log-normalized edge sets); only communities with ≥20 HOGs are shown. Nodes are HOGs and edges are significant ERC pairs. Node color shows the mean Z-score of the normalized BL difference (carnivorous − non-carnivorous) for each HOG: red = relative acceleration in carnivores; blue = relative conservation. Numbers indicate community IDs. **Right:** Heatmap of community-level mean Z-scores per carnivorous terminal lineage (columns). For each community (row), the top stripe uses Z-scores from normalized BLs, and the bottom stripe uses Z-scores from log-normalized BLs; each cell is the average across all HOGs in that community. The same red–white–blue scale (−1 to +1) is used in both panels. Together, these views highlight communities enriched for lineage-specific acceleration or conservation.

Among these, Community 10 stood out as the only cluster exhibiting consistent acceleration across three independent CP lineages—Sarraceniaceae, Nepenthaceae, and Lentibulariaceae—with the strongest shifts observed in the Sarraceniaceae species *H. tatei* and *S. purpurea*. Closer examination of the HOGs in Community 10 (Figure 5) and their lineage-specific relative rate shifts revealed that most inferred ERC hits were primarily driven by accelerated rates in Sarraceniaceae, with additional contributions from other CP lineages (Supplementary Figure 15-17). Among reconciled gene trees, we observed that several HOGs show accelerated BLs in the terminal branches of Sarraceniaceae species, often displayed shorter BLs in the internal branch leading to the divergence of *Heliamphora* and *Sarracenia*. This phylogenetic incongruence likely reflects incomplete lineage sorting, where ancestral genetic variation persists through speciation events and causes discordance between gene trees and the species tree (Baum & Smith, 2012), or lineage-specific retention of distinct paralogs following an ancestral whole-genome duplication (Alabady et al., 2026; Albert et al., 2025) that preceded the divergence of *Heliamphora* and *Sarracenia* (Supplementary Figure 18).

**Figure 5.**
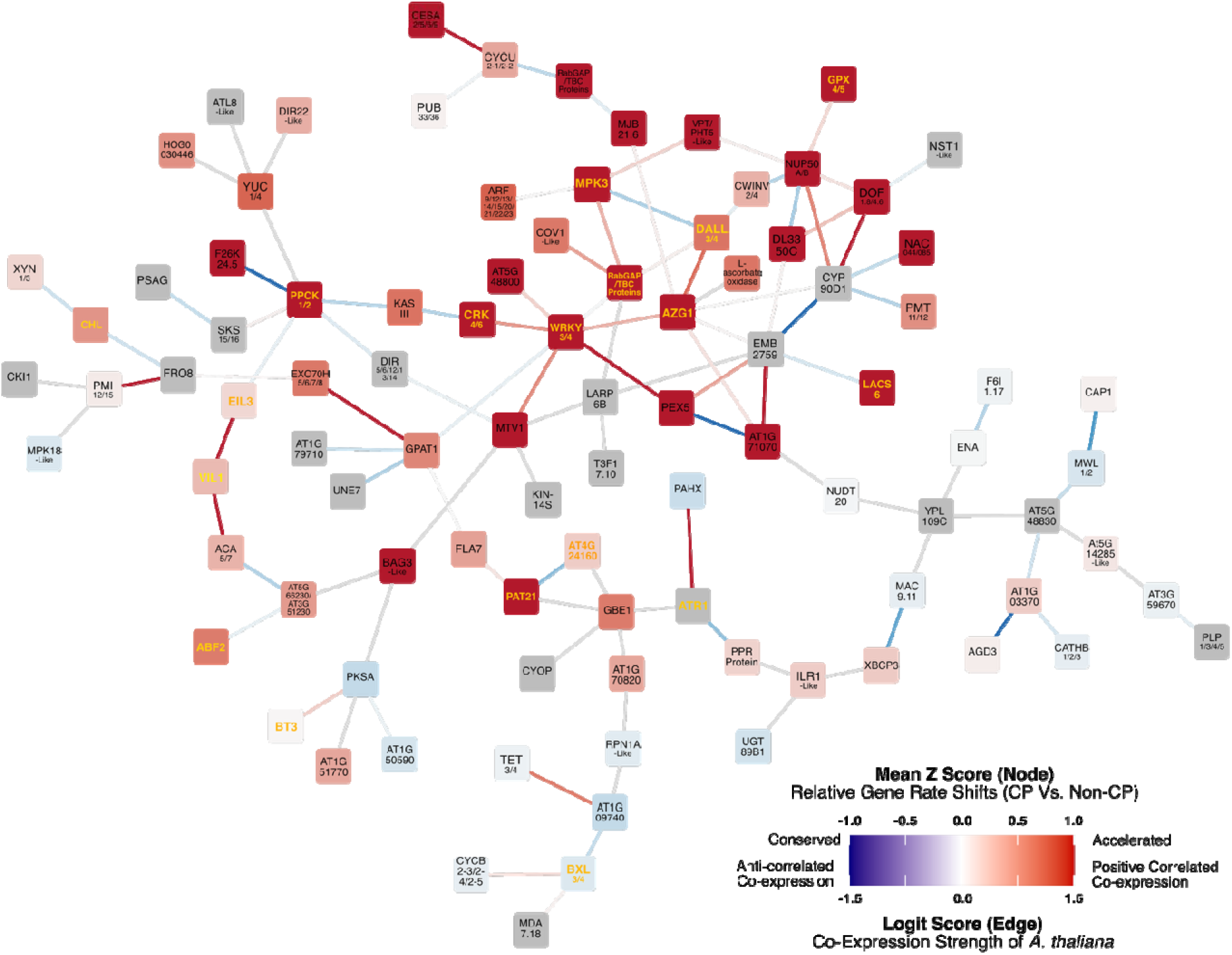
**Community 10 network** visualized with average CP rate shifts (node color) and Arabidopsis co-expression links (edge color). Nodes represent hierarchical orthogroups (HOGs), colored by their mean Z-score across CP terminal branches (from normalized branch-length ERC analysis). Warmer colors indicate accelerated evolution in CPs; cooler colors indicate conservation. Edges represent co-expression relationships between Arabidopsis orthologs derived from the ATTED-II union dataset (v23-11), which integrates thousands of Arabidopsis transcriptome profiles (microarray and RNA-seq) corrected with ComBat-PCA and normalized by sub-agging. For each HOG–HOG pair, the mean Logit Score (LS) of all ortholog-pair edges was used. Red edges (LS > 0) indicate positive co-expression, blue edges (LS < 0) indicate anti-correlated expression, and gray edges lack ATTED-II support. LS is ATTED-II’s logit-transformed probability that two genes are co-expressed (|LS| ∝ co-expression strength). Genes associated with stress response are in bold and highlighted in orange-yellow text.

To account for potential errors stemming from uncertainties in BL reconciliation for HOGs with complex duplication and retention histories, we re-estimated ERCs for gene pairs in Community 10 using recalculated BLs for *H. tate*i and *S. purpurea*, obtained by averaging their respective BLs with that of the internal Sarraceniaceae branch. Recalculation of Community 10 edges recovered 75.2% of the significant ERC hits based on the original significance threshold, suggesting that the inferred evolutionary relationships are relatively robust to noise and reconciliation uncertainty.

### Accelerated ERC community associated with stimulus and stress responses

To test whether the convergent functional signals identified in lineage-specific analyses of accelerated genes are also captured by network-based approaches, we performed GO enrichment analyses on ERC communities containing ≥ 20 HOGs (full results in Supplementary Data). Here, we focus on Community 10, which showed significant enrichment for developmental, metabolic, and regulatory processes (Figure 6, Supplementary Figure 19-21). Notable GO BP terms include developmental process and its subcategories (anatomical structure development, reproductive process, post-embryonic development). The community is also enriched for primary and macromolecule metabolic processes as well as localization and transport terms. Consistent with the lineage-specific accelerated HOG analyses, stimulus- and stress-responsive categories are overrepresented (e.g., response to stimulus, response to oxidative stress, hormone-mediated signaling pathway).

**Figure 6.**
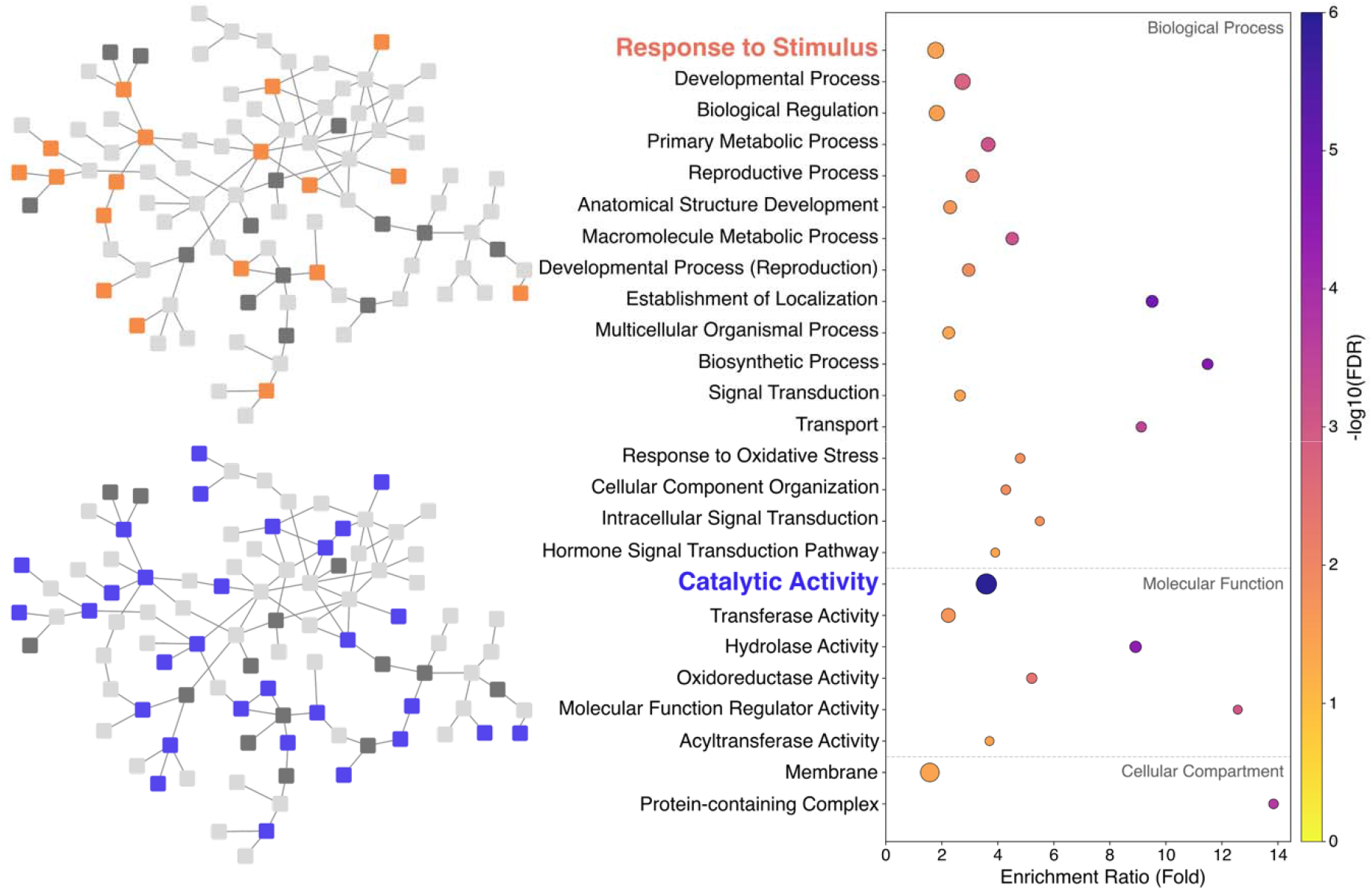
Gene Ontology (GO) enrichment and functional visualization of Community 10. **Right:** Bubble plot summarizing significantly enriched GO terms across BP, MF, and CC. The x-axis shows fold enrichment (observed/expected); bubble size indicates the number of HOGs annotated to each term (≥5 shown), and color denotes significance (−log FDR). Terms are grouped by GO namespace (BP → MF → CC) and ranked within each group by annotated HOG count. Enrichment highlights developmental and metabolic processes (BP), catalytic and transferase activities (MF), and membrane-associated protein complexes (CC). **Left:** Subnetworks of Community 10 highlighting HOGs annotated to the top enriched BP term Response to Stimulus (top) and the top MF term Catalytic Activity (bottom). Colored nodes indicate annotated HOGs; gray nodes show unannotated members.

Within Community 10, HOG0024011, annotated as a nuclear pore complex protein 50 a/b-like gene family (NUP50A/B), represents a high-connectivity hub and exhibits the strongest signal of convergent acceleration. In *A. thaliana*, NUP50A and NUP50B localize to the nucleoplasm and facilitate importin-α/RanGTP–dependent disassembly of nuclear import complexes—a critical step for the nuclear entry of stress- and hormone-responsive transcription factors (Tamura et al., 2010; Zhou et al., 2025). Consistent with this function, NUP50 regulates cell wall-related genes and is associated with seed longevity and salinity stress tolerance (Oishi et al., 2024; Zhou et al., 2025), while other nucleoporins (e.g., NUP85, NUP160) broadly modulate abscisic acid (ABA)-, salt-, and cold-response pathways (Tamura et al., 2010).

NUP50’s ERC partners connect this nuclear-transport hub to redox, nutrient, and transcriptional modules, suggesting a tightly integrated functional network. These partners include CYP90D1 (HOG0019912), a cytochrome P450 enzyme involved in brassinosteroid biosynthesis (Enoki et al., 2023); GPX4/5 (HOG0010780), glutathione peroxidases linked to oxidative stress, ABA, and drought responses (Ramos et al., 2009; Zhang et al., 2018); and VPT/PHT5-like (HOG0019750), vacuolar phosphate transporter family proteins central to Pi homeostasis and active during flowering (Luan et al., 2018). Additional partners include CWINV2/4 (HOG0010587), cell-wall invertases such as CWINV4 that are required for nectar secretion (Ruhlmann et al., 2009); DOF1.8/DOF4.6(HOG0002661), nuclear DOF transcription factors (Zou & Sun, 2023); and DL3350C (HOG0016456), DUF506 family proteins with reported roles in abiotic-stress responses (Dong et al., 2023). We hypothesize that selection on nucleocytoplasmic trafficking—potentially mediated by NUP50—has favored coordinated evolutionary change among these developmental, metabolic, and signaling components, consistent with Community 10’s enrichment for localization/transport, stimulus response, and primary/macromolecule metabolism terms.

A second hub, HOG0013840 (WRKY3/4), also shows acceleration across several carnivorous lineages, implicating stress-responsive transcriptional regulation as another core module potentially adapted for carnivory. In *A. thaliana*, WRKY3 and WRKY4 are nuclear DNA-binding proteins that positively regulate defense against necrotrophic pathogens and mediate salicylic acid (SA)–jasmonic acid (JA) crosstalk (Lai et al., 2008). Within Community 10, WRKY3/4 connects to HOGs that link signaling and intracellular trafficking with gene regulation. These include CRK4/6 (HOG0018949), calcium-dependent protein kinases (HOG0018949), AT5G48800 (HOG0005380), a BTB/POZ E3-adapter–like protein, and RabGAP/TBC domain containing proteins (HOG0016261). Additional partners include AZG1 (HOG0018011), a cytokinin importer that binds and stabilizes the auxin efflux carrier PIN1 and tunes root salt responses (Tessi & Maurino, 2024); PEX4/5 (HOG0019805) peroxisomal import receptor proteins required for the import of peroxisome-bound proteins containing one of two peroxisomal targeting signals (PTS1 or PTS2) (Khan & Zolman, 2010) (Supplementary Figure 22); MTV1 (HOG0023320), an ENTH-domain protein mediating trans-Golgi-network-to-vacuole trafficking (Heinze et al., 2020); and GPAT1 (HOG0007589), a sn-glycerol-3-phosphate 2-O-acyltransferase involved in lipid metabolism (Wendel et al., 2013).

Taken together, while the exact mechanisms remain to be tested, our ERC results are consistent with a co-evolving module linking signaling, intracellular trafficking, peroxisomal import, small-molecule transport, lipid metabolism, and WRKY-mediated transcription—a functional assemblage potentially central to the adaptive evolution of carnivory.

### Co-expression of hub genes in non-CP model system suggests ancestral stress-response origins

Using *A. thaliana* co-expression data (Obayashi et al., 2022), we visualized the co-expression strength (logit score, LS) among the *A. thaliana* homologs represented in the inferred ERC network (Supplementary Figure 23) and within Community 10 (Figure 5). Among the 5,508 significant ERC hits, 3,936 were associated with *A. thaliana* LSs (ERC hits without any associated LSs generally contain HOGs with no *Arabidopsis* homologs), with 598 edges (15.2%) showing significantly positive co-expression (LS > 1.5). Although only a modest fraction of inferred coevolving gene pairs is co-expressed in *A. thaliana*, the ERC-inferred network nevertheless exhibits significantly greater positively correlated co-expression than expected by chance, based on a one-sample, one-sided t-test comparing edge LS values to zero (network mean edge LS = 0.37, median LS = 0.10; p [mean LS > 0] = 3.8 ×10□□¹).

Within Community 10, the strongest co-expression signals clustered around the same hub genes identified in our ERC network—NUP50 and WRKY3/4, suggesting that these hub-centered, stress-responsive gene clusters represent a core stress-responsive regulatory module that has been maintained in both carnivorous and non-carnivorous angiosperms. Additionally, in a separate region of community 10, we observe a connected group of nodes that exhibit co-expression composed of EIL3 (Ethylene Insensitive 3–like), VIN3 (Vernalization Insensitive 3), and ACA5/7 (α-Carbonic Anhydrase 5/7). These genes are involved in ethylene signaling (Li et al., 2013), chromatin-mediated stress responses (Bond et al., 2009), and redox regulation (Fedorchuk et al., 2021), respectively— processes broadly implicated in plant defense and environmental adaptation.

The observation that these Community 10 orthologs are co-expressed in the non-CP *A. thaliana* suggests that this module represents an ancestral, stress- and defense-responsive network that was later recruited and co-opted during the evolution of carnivory (de Vries & de Vries, 2020; Renner et al., 2018). Such conservation of co-expression topology supports the hypothesis that carnivorous lineages, particularly Sarraceniaceae, may have repurposed pre-existing regulatory circuits governing environmental and defense responses for trap function and nutrient acquisition.

## Discussion

### Accelerated rates and gene family expansions underpin the genomic convergence of carnivory

In comparison to previous studies of CP genome evolution, our evolutionary rate–based comparative analysis not only recapitulates previously reported functional enrichments (e.g., GO terms associated with stress response, metabolism, and signal transduction) but also provides insight into the evolutionary processes that facilitated their recruitment for carnivorous functions. Across independent carnivorous lineages, genes exhibiting elevated evolutionary rates also tend to occur in expanded gene families, suggesting a close association between gene duplication and accelerated sequence evolution during the emergence of carnivory (Carretero-Paulet et al., 2015). These duplications arise both from ancient whole-genome duplications that predate the emergence of carnivory and from tandem duplications accumulated throughout the diversification of carnivorous plants (Alabady et al., 2026; Albert et al., 2026; Albert et al., 2025). Compared with conserved genes, accelerated genes are more frequently duplicated and experience relaxed or weakened purifying selection pressures—with a smaller subset showing evidence of positive selection—consistent with their potential for functional diversification and neofunctionalization in support of carnivorous adaptations (Carretero-Paulet et al., 2015; Palfalvi et al., 2020; Renner et al., 2018).

We note, however, that the association between gene family expansion and acceleration may be partly influenced by the greater topological complexity of highly paralogous families (Sarton-Lohéac et al., 2025; Smith & Hahn, 2021; Smith et al., 2022), because our HOG-based reconciliation pipeline averages BLs across multiple paralogs (Forsythe et al., 2025), potentially inflating HOG-level rate estimates in some cases. Nonetheless, the concordance between our findings and lineage-specific analyses in *U. gibba* (Carretero-Paulet et al., 2015), together with additional evidence for relaxed and weakened purifying selection in the accelerated gene sets (and vice versa), supports the inference that the observed relationship is not purely an artifact and likely reflects genuine evolutionary dynamics (Palfalvi et al., 2020; Renner et al., 2018). Moreover, similar associations between gene duplication, relaxed purifying selection, and accelerated sequence evolution have been reported in other plant and animal systems (Fischer et al., 2014; Lynch & Conery, 2000; Pegueroles et al., 2013), suggesting that the patterns we observe in CPs are part of a broader evolutionary trend in which novel gene functions frequently arise through the recruitment and divergence of ancestrally duplicated genes.

### Complex angiosperm genome dynamics leads to convergence on gene functions rather than gene identity

Most genes that evolve unusually rapidly or slowly in CPs are lineage-specific, reflecting strong phylogenomic constraints within the broader context of dynamic angiosperm genome evolution (De Bodt et al., 2005; Pavlovič, 2025). Repeated whole-genome duplications (and even triplications), followed by extensive gene loss and contraction, produced distinct genomic inventories in the ancestors of modern carnivorous lineages—even before the emergence of carnivory (Qiao et al., 2022). Importantly, additional rounds of genome expansion and contraction occurred after carnivory evolved, as seen in Carnivorous Caryophyllales (i.e. crown group containing Droseraceae and Nepenthaceae), Lentibulariaceae (Fleck et al., 2025; Palfalvi et al., 2020; Robledillo et al., 2025; Wicke et al., 2014), and Sarraceniaceae (Alabady et al., 2026; Albert et al., 2025), further shaping lineage diversification and enabling adaptive radiations within several CP families. Furthermore, recent phylogenomic studies show that hybridization and introgression are pervasive in multiple carnivorous lineages (e.g. carnivorous Caryophyllales and *Sarracenia*) and likely facilitated the adaptive radiation of new carnivorous trap types (e.g. *Nepenthes*), adding an additional layer of reticulate genome evolution to these systems (Baldwin, 2025; Robledillo et al., 2025; Scharmann et al., 2021). As a result of these complex genome evolutionary histories, natural selection for carnivory acted on different sets of available genes in each lineage (Pavlovič, 2025), leading to convergence at the level of biological function rather than gene identity.

Across independent carnivorous lineages, accelerated and expanded gene families are consistently enriched for defense- and stress-response pathways compared with their non-carnivorous relatives, although the magnitude of acceleration varies. This pattern reinforces the view that genome duplications provided the raw material for innovation: duplicated stress-related genes were repeatedly co-opted and repurposed for carnivorous functions such as prey attraction, digestion, and nutrient assimilation (Alabady et al., 2026; de Vries & de Vries, 2020; Fukushima et al., 2017). In other words, the molecular convergence underlying plant carnivory primarily reflects the repeated redeployment of similar functional pathways—not the reuse of the exact same orthologs.

A clear example of this principle is seen in the WRKY transcription factor family, which exhibits convergent acceleration alongside lineage-specific co-option for carnivorous functions. For instance, WRKY6 has been shown to be both duplicated and co-opted for carnivorous function in Droseraceae (Palfalvi et al., 2020). Our analyses revealed extensive variation in WRKY copy number among carnivorous taxa, ranging from fewer than five copies in Lentibulariaceae to more than twenty in *C. follicularis* and Sarraceniaceae species (Supplementary Table G). These differences indicate that distinct WRKY paralogs were independently recruited from ancestral copies, producing parallel regulatory roles in trap development and stress-associated metabolism despite their divergent origins. This family thus encapsulates the broader evolutionary trend observed across CPs—functional convergence built upon lineage-specific genomic substrates.

Similarly, in the case of genes associated with the response to light stimulus, only a handful exhibited convergent acceleration across multiple lineages, while the majority showed lineage-specific acceleration. These accelerated genes belong to shared functional modules involved in photoreception, signaling, transcriptional regulation, and metabolic adjustment. This pattern again points to parallel fine-tuning of similar functional pathways, rather than convergence on identical gene homologs, as the dominant mechanism shaping the evolution of carnivorous physiology.

Taken together, our results indicate that while convergence at the level of individual genes is relatively rare (e.g. adaptive molecular convergence of several digestive enzymes of independent CP lineages (Fukushima et al., 2017)), convergence in gene function is widespread. Thus, the evolution of carnivory shows how deep genomic histories can limit strict molecular reuse, yet still allow repeated innovation via the recruitment of alternative, functionally equivalent genes.

### Convergent acceleration of light-stimulus–responsive genes suggests links to trap development

Among the genes accelerated in CPs, many associated with the response to light stimulus category may have contributed directly to the evolution of trap morphology and physiology. Genes involved in polarity and morphogenesis (e.g., WAV2, HAT3/5, HLS1, IAA29) likely underlie modifications to leaf curvature and lamina polarity that gave rise to the distinctive concave, cup-like, or tubular architectures of traps (Bou-Torrent et al., 2012; Mochizuki et al., 2005; Ohto et al., 2006; Tang et al., 2022). The concurrent acceleration of cell wall–remodeling and surface-structure genes (e.g., XTH15/16, SCP2, KCS1) further suggests that epidermal flexibility and the development of specialized surface properties (Todd et al., 1999; Yokoyama & Nishitani, 2001; Zheng et al., 2008)—such as slippery pitcher walls or absorptive glandular tissues—evolved in tandem with these morphological changes.

Similarly, proteostasis and signaling regulators such as COP1, POB1, and CUL4—key modulators of photomorphogenesis and developmental transitions—may mediate light-dependent differentiation between glandular, absorptive, and photosynthetic tissues within traps (Han et al., 2020; Orosa et al., 2017; Zhang et al., 2008). The acceleration of hormone and stress integrators (e.g., MPK3, ARR7, ERF5) (Jagodzik et al., 2018; Moffat et al., 2012; O’Brien & Benková, 2013) and Ca² and ion transporters (e.g., CML23, CAS, KCO3) further points to refined environmental responsiveness in trap tissues (Dabravolski & Isayenkov, 2021; Li et al., 2022; Vadassery et al., 2012), enabling coordinated growth, prey detection, and movement in active traps such as *Dionaea* and *Aldrovanda* (Böhm et al., 2016; Iosip et al., 2023).

Because carnivorous function is energetically costly and closely tied to photosynthetic activity, many CPs exhibit a tradeoff between carnivory and photosynthesis, often reducing trap activity under low-light conditions (Givnish et al., 2018; Karagatzides & Ellison, 2009). This ecological dynamic implies that selection has favored individuals capable of modulating this balance efficiently (Gilbert et al., 2025). The repeated acceleration of light-stimulus– responsive genes therefore may also reflect adaptive fine-tuning of the molecular mechanisms governing this resource-allocation tradeoff (Givnish et al., 1984), optimizing trap performance and photosynthetic efficiency under variable light environments (Gilbert et al., 2025).

Taken together, these gene categories highlight a modular reorganization of epidermal, developmental, and physiological networks underlying trap evolution. The repeated acceleration of genes controlling light-mediated growth, surface differentiation, and stress signaling supports the view that carnivorous traps arose through the progressive redeployment and fine-tuning of preexisting photoregulatory and epidermal programs, rather than through the emergence of entirely novel genetic pathways (Agrawal et al., 2022; Fukushima et al., 2015; Palfalvi et al., 2020).

### Accelerated gene network communities reflect core stress and defense pathways convergently co-opted in CPs

This study identifies a co-evolving gene network community that exhibits pronounced lineage-specific acceleration in Sarraceniaceae and moderate acceleration in several other CP lineages, including the independently evolved pitcher plant N. gracilis and the Lentibulariaceae species *P. moranensis* and *G. aurea*.

Consistent with our lineage-specific enrichment results, the accelerated ERC community is significantly enriched for stimulus- and stress-response functions— BPs that are repeatedly accelerated across CP lineages. Furthermore, many of the ERC-linked gene pairs are co-expressed in non-carnivorous model species, suggesting that this network represents a conserved stress- and stimulus-response module that predates the diversification of major eudicot lineages.

It is likely that components of this ancestral pathway were repeatedly co-opted for carnivorous functions, with lineages such as Sarraceniaceae potentially recruiting a greater proportion of its central hub genes for trap-specific adaptations. The shared acceleration of this network across independently evolved carnivorous lineages highlights the convergent repurposing of a conserved stress-response module for novel carnivorous functions (e.g. prey attraction, capture & digestion). Collectively, these findings support the view that the molecular evolution of plant carnivory represents a functional transition from defense to offense— shaped by lineage-specific genome histories and driven by convergent shifts in evolutionary rate dynamics.

## Materials and Methods

### Computing environment

All computationally intensive bioinformatic analyses were executed on Ubuntu 22.04.5 LTS (kernel 6.8.0-60- generic, x86_64) and managed by SLURM 21.08.5, with 64 physical cores (128 threads with hyperthreading) at 2.6 GHz, 1 TB RAM, and 256 TB local storage. Software was installed and managed via conda/mamba. A workflow diagram summarizing the analytical pipeline described below, from input sequences to ERC network community inference, is provided in Supplementary Figure 25.

### Sampling and processing of CP and non-CP datasets

Protein-coding DNA (CDS) sequences for 30 sampled carnivorous (n = 11) and non-carnivorous taxa (n = 19) were obtained from publicly available datasets (Supplementary Data). For *H. tatei* and *S. purpurea*, respectively, raw RNA-seq reads were obtained from Liu & Smith (2025) and downloaded from the NCBI Sequence Read Archive (accessions PRJDB3436). These transcriptomes were assembled de novo with Trinity v2.15.2 (Grabherr et al., 2011), then annotated following Liu and Smith (Liu & Smith, 2025). All CDS files were preprocessed with the Python program Squeakuences (Lane et al., 2024) to detect and correct formatting irregularities before translation into protein sequences. Dataset completeness was assessed with BUSCO v6.0.0 (Manni et al., 2021), using the “Eudicotyledons odb10” dataset.

Because source genomic resources ranged from chromosome-level, well-annotated genomes to low-coverage, single-tissue transcriptomes, we applied additional filtering prior to downstream analyses. We used CD-HIT v4.8.1 (Fu et al., 2012) to remove identical sequences and putative alleles or alternative splice isoforms at 99% identity. We then inferred orthologous gene families with OrthoFinder v3.0.1b1 (Emms & Kelly, 2019), supplying a user-defined constraint species tree of the 30 sampled taxa (-s) so that hierarchical orthogroup (HOG) inference and gene-tree/species-tree reconciliation were guided by the known phylogenetic relationships among taxa rather than a species tree inferred de novo. OrthoFinder was run in multiple-sequence-alignment gene-tree mode (-M msa) with DIAMOND ultra-sensitive sequence searches (-S diamond_ultra_sens), MAFFT alignments (-A mafft), and FastTree gene-tree inference (-T fasttree); all other parameters were left at their defaults (full command provided in Supplementary Data). The resulting HOGs (OrthoFinder N1 level) served as input for all downstream ERCnet/phylogenomic analyses.

### Phylogenomic analyses with gene–tree/species–tree and branch-length reconciliations

Phylogenomic analyses (Phylogenomics.py) were performed on OrthoFinder outputs as the first step in the ERCnet workflow (Forsythe et al., 2025). We required ≥15 species represented per HOG and allowed ≤20 gene copies per species per HOG. Gene families passing these filters were processed through a pipeline that used MAFFT v 7.525 to generate amino-acid alignments (Rozewicki et al., 2019), TAPER to identify and remove outlier sequences and composition/heterogeneity-driven noise (Zhang et al., 2021), Gblocks v0.91b to trim poorly aligned (we retained any sequence in the alignments with consensus coverage > 95%) or ambiguous regions (Castresana, 2000), and IQ-TREE v 2.3.6 to infer a gene tree for each HOG (Minh et al., 2020).

After gene trees were inferred for all retained families, gene–tree/species–tree (GT/ST) reconciliation (GTST_reconciliation.py) was performed (Forsythe et al., 2025). Each HOG gene tree was mapped onto the species tree to infer gene duplications and losses and to reconcile topological conflicts between gene and species histories (Forsythe et al., 2025). The resulting reconciled trees were used for downstream analyses.

Following GT/ST reconciliation, we ran the BL reconciliation stage of the ERC analyses (ERC_analysis.py) to obtain BL estimates for internal and terminal branches. BL reconciliation extracts BL measurements in a species-tree–aware framework, enabling direct comparisons across gene trees (Forsythe et al., 2025).

### Branch-length consensus, log transformation, and normalization

To account for estimation uncertainty introduced during upstream steps of the ERCnet pipeline (e.g., gene-tree inference and gene-tree/species-tree reconciliation), we performed five replicate ERCnet runs using identical input parameters. These runs produced five raw BL files (as output BL_results/ bxb_BLs.tsv), which we used to quantify upstream noise and to remove BL estimates that were inconsistent across replicates. A custom Python script (BL_Consensus.py) combined the five replicates into a consensus BL matrix as follows. For each HOG × branch cell with an estimate in at least three of the five replicates (“classifiable” cells), we computed the replicate median and median absolute deviation (MAD), and flagged a replicate value as discordant (an outlier) when it deviated from the median by more than 2 × MAD. Conventional MAD-based rejection thresholds span 2–3 × MAD (Leys et al., 2013); we adopted the more stringent end of this range (2 × MAD), a choice supported empirically by the sensitivity analysis below. This robust criterion is appropriate because replicate raw BL distributions are often heavy-tailed; unlike the standard deviation, the MAD is not inflated by the outliers being detected, keeping the threshold anchored to the central cluster of replicates (Leys et al., 2013; Rousseeuw & Croux, 1993). A cell was retained when more than 65% of its present replicates were concordant, and its consensus BL was set to the mean of the concordant values. Across the 56 estimable branches of the species tree, 70.2% of the 1,793,313 individual replicate estimates were averaged into the consensus and 29.8% were excluded as outliers. This procedure operates within cells rather than removing whole gene families: 12,490 of 12,518 HOGs with an estimate in ≥ 1 replicate (99.8%) are retained (median 30 of 56 possible branches per HOG; the 28 exceptions lacked any cell with ≥3-replicate coverage), and no HOG was removed in the consensus step (Supplementary Figures 26–28). To confirm robustness to the chosen threshold, we varied the MAD threshold from 0.5 to 5.0 (Supplementary Figure 29): the fraction of replicate estimates excluded as outliers declines steeply to MAD = 2.0 (29.8%) and then plateaus, so that 2.0 sits at the elbow of the curve — excluding clearly discordant replicate values while retaining the bulk of consistent estimates. The cross-replicate variability targeted by this step arises predominantly from uncertainty in gene-tree inference for weakly informative HOGs rather than from alignment. On average, internal branches exhibited shorter genome-wide mean BLs than terminal branches (0.056 vs. 0.123; P < 0.001; Supplementary Figures 2 & 3).

After generating consensus BLs, we manually inspected alignments for HOGs with extreme values. Cases deemed likely to reflect alignment error or contamination were removed prior to normalization. To account for genome-wide variation in substitution rates among lineages—arising from differences in mutation rate and the evolutionary time specific to each terminal and internal branch—we computed the genome-wide mean raw BLs for each species-tree branch across all HOGs and used these per-branch means to normalize HOG-specific BLs, as described below, to account for lineage-specific rate differences and variation in branch durations (Forsythe et al., 2025; Rasmussen & Kellis, 2007).

Raw branch lengths (amino-acid substitutions per site) should not be compared directly across lineages because such values conflate evolutionary rate with divergence time. Instead, all of our downstream cross-lineage inferences operate on per-branch normalized values, obtained by dividing each HOG’s branch length by the genome-wide mean for that same branch. For example, HOG0000629 has a raw branch length of 0.099 on the terminal branch leading to *C. follicularis* (N15_to_CepFol), whereas the genome-wide mean for that branch is 0.151 (n = 8,421 HOGs), giving a normalized value of 0.099 / 0.151 = 0.655. Because the focal gene and the genome-wide reference share identical elapsed time and lineage context, their ratio is a pure measure of rate deviation of a focal relative to the genome-wide background. This is conceptually equivalent to a local molecular-clock correction, simultaneously controlling for both branch-specific divergence time and lineage-wide differences in overall substitution rate, and yielding a quantity that is comparable across lineages. Future incorporation of time-calibrated, absolute-rate estimates could provide complementary, fully time-scaled comparisons of evolutionary rates across carnivorous lineages.

For log-normalized BLs, consensus raw BLs were first log-transformed and then normalized using the corresponding genome-wide mean of the log-transformed BL. This reduces heteroskedasticity and limits the influence of long-branch outliers (Box & Cox, 1964; Little et al., 2025). However, log transformation can also magnify relative differences among short branches in downstream ERC analyses, so we retained both datasets and restricted our interpretations to ERC hits overlapping between them as a conservative approach.

To assess phylogenetic signal in both normalized and log-normalized BLs, we estimated Pagel’s λ (Pagel, 1999) separately for each HOG with at least 10 branch estimates by restricted maximum likelihood, using a species-tree correlation matrix in which each branch is placed at its midpoint, and tested λ > 0 with a likelihood-ratio test followed by Benjamini–Hochberg correction across HOGs.

### ERC and Network Analyses

ERC analyses (ERC_analysis.py) were run on both datasets (normalized and log-normalized BL matrices) in all-versus-all, branch-by-branch (BxB) mode, which estimates covariation among all HOG pairs while accounting for shared evolutionary history. Significant pairs were defined as those with slope (β) > 0.75, Pearson FDR-adjusted P < 1 × 10, and >10 overlapping branches in both datasets. The two significant-edge sets were then intersected, retaining only pairs significant in both, to yield the final network edge list. Networks were visualized in Cytoscape (v3.10.3), and communities were identified using GLay (Su et al., 2010).

### Arabidopsis co-expression framework for functional interpretation of ERC networks

To connect ERC-derived relationships in CPs with functional co-expression patterns in a model system, we visualized selected network communities—particularly Community 10 (Figure 5)—using *A. thaliana* co-expression data from ATTED-II (release v23-11; dataset Ath-u.v23-11.G19783-S35094.combat_pca.subagging.z.d) (Obayashi et al., 2022). This ATTED-II release integrates thousands of *A. thaliana* transcriptome profiles from both microarray and RNA-seq experiments, batch-corrected with ComBat-PCA and aggregated using subagging Z-score normalization to yield robust, genome-wide co-expression estimates. The LS represents the logit-transformed probability that two *A. thaliana* genes are co-expressed across the integrated dataset (LS ≈ 0 indicates random expectation; larger LS values reflect stronger positive co-expression, while negative LS values indicate anti-correlated expression).

Each CP HOG was mapped to its *A. thaliana* ortholog(s) via TAIR AGI locus IDs and cross-referenced to Entrez Gene IDs through Ensembl Plants BioMart (Kinsella et al., 2011). For every HOG–HOG pair in the ERC network, all available *A. thaliana* ortholog-pair edges were retrieved from ATTED-II; when multiple ortholog pairs existed, their LS values were averaged to yield a single edge-level LS.

### Identification of accelerated & conserved HOGs in each CP Lineage

To identify HOGs that are more accelerated or conserved in each CP lineage compared to non-CPs, we computed, for each HOG, Z-scores representing the deviation of each CP BL from the mean of all non-CP branches, scaled by the corresponding standard deviation, separately for normalized and log-normalized BLs. Because these Z-scores are computed from per-branch normalized and log-normalized BLs rather than raw branch lengths, they are directly comparable across lineages, independent of branch-specific divergence time and lineage-specific genome-wide rate variation. The non-CP reference included both terminal and internal branches (median 25 non-CP branches per HOG, range 3–39; Supplementary Table D; per-HOG counts in Supplementary Data). Because normalized BLs are more variable on internal than on terminal non-CP branches (median within-HOG SD: 0.52 vs. 0.32 for normalized and 0.084 vs. 0.039 for log-normalized BLs), including internal branches widens the reference distribution and makes the resulting Z-scores more conservative. For the carnivorous focal branches, however, we limited our analyses to terminal branches leading to CP taxa, given that BL estimates for internal branches could be more prone to uncertainty in the context of complex genome histories (Mendes & Hahn, 2016). For each CP lineage, HOGs with Z ≥ 1 or Z ≤ −1 in both the normalized and log-normalized datasets were classified as accelerated or conserved, respectively. To assess whether data quality influenced these classifications, we regressed the per-lineage fraction of HOGs with |Z| ≥ 1 on BUSCO completeness and mean alignment length (Gblocks-trimmed alignments used for gene-tree inference, averaged over the HOGs contributing to each fraction), and in a second model additionally on the log genome-wide mean raw BL of the terminal branch, using ordinary least squares. Transcriptome-and genome-derived CP lineages were compared with a Mann–Whitney U test.

To test whether accelerated HOGs tend to contain more paralogs than conserved HOGs within each CP species, we quantified the number of homologous gene copies per species in each reconciled HOG tree. We then compared the distributions of gene counts between the accelerated and conserved sets using both parametric and non-parametric tests. Specifically, Welch’s t-tests were used to evaluate differences in the mean number of paralogs, whereas Mann– Whitney U and Mood’s median tests were applied to assess differences in medians and overall distributional shifts. These complementary analyses provided a robust statistical framework for determining whether accelerated genes are significantly more expanded in CP genomes than conserved genes.

### Identification of ERC communities convergently accelerated or conserved in CPs

We restricted analyses to GLay communities with >20 members (HOGs) before testing for convergent acceleration or conservation in CPs versus non-CPs. Communities with a significantly positive mean Z-score across CP branches were classified as convergently accelerated; significantly negative values indicated convergent conservation. Community-level Z-scores were computed and evaluated independently on the normalized and log-normalized BL datasets. For each HOG, we calculated the mean Z-score across all available CP branches, then defined the community Z-score as the mean of these per-HOG means. Communities were called convergently accelerated (positive) or conserved (negative) if the community Z-score was significant after FDR adjustment (FDR-adjusted P < 0.05; marginal at < 0.10) in both the normalized and log-normalized datasets, with consistent direction.

To further verify and localize these patterns, we performed lineage-specific Z-score analyses to identify communities exhibiting rate shifts restricted to a CP lineage or species. For each CP lineage, we extracted branch-specific Z-score columns and tested whether community means differed significantly from zero using one-sample t-tests and Wilcoxon signed-rank tests (two-sided). These tests were conducted for each community independently in both the normalized and log-normalized datasets, with FDR correction applied across all communities per lineage. Communities showing significant deviations (FDR-adjusted P < 0.05) in a single lineage were considered lineage-specifically accelerated or conserved, whereas those significant in multiple independent lineages were interpreted as showing convergent patterns of rate change.

### Gene ontology (GO) enrichment analysis

#### Annotation

We performed GO enrichment at the level of HOGs. Because a single HOG may map to multiple Arabidopsis orthologs (and therefore multiple GO terms), for each HOG we assembled the union of unique GO terms associated with its mapped *A. thaliana* orthologs from our annotation table (HOG_AT_GO_Table.tsv), using a custom Python script (run_hog_go_enrichment.py). Annotations were not propagated to GO ancestors; only directly annotated terms were used.

#### Study and background sets

Each enrichment test compares a study set (the foreground HOGs of interest) against a background set (the reference population of HOGs from which the study set is drawn). In every analysis the study set was intersected with its background so that the study set was a strict subset of the background. The two sets were defined as follows:

#### Lineage-specific tests

For each CP lineage, the study set comprised the HOGs classified as accelerated (Z-score ≥ 1) or conserved (Z-score ≤ −1) in that lineage (the two directions tested as separate study sets). The background was the set of HOGs assayable in that lineage — i.e., HOGs with a Z-score for that lineage and at least one GO annotation — not the full set of OrthoFinder HOGs. Restricting the background to HOGs present in the lineage avoids spurious enrichment driven by HOGs absent from that species.

#### ERCnet Community-level tests

Each significantly accelerated GLay community was tested as a study set, both individually and as a pooled union across accelerated communities; significantly conserved communities were tested the same way. For all community-level tests the background was fixed as all ERC-input HOGs with at least one GO annotation (n=8848 HOGs).

#### Enrichment test

For each candidate GO term, GOATOOLS (Klopfenstein et al., 2018) constructed a 2×2 contingency table of study-vs-background membership and tested over-representation with a one-sided Fisher’s exact test (hypergeometric model, “greater” tail), using the go-basic.obo ontology graph (Aleksander et al., 2023). Before testing, we (i) collapsed GO terms that annotated identical sets of background HOGs into a single representative term, and (ii) removed sparsely represented terms by requiring ≥2 background HOGs per term.

Multiple testing was controlled with the Benjamini–Hochberg FDR across the tested representative terms. For each representative term we report its namespace (BP, MF, or CC), the study and background counts and totals, the uncorrected and FDR-adjusted P-values, and the enrichment ratio [(study_count/study_total)/(pop_count/pop_total)]. The collapsed-term groups (each representative term and its members) are recorded in the run logs and provided in Supplementary Data.

### Subcellular compartments assortativity of ERC networks

Beyond community- or lineage-level enrichment, we asked whether proteins annotated to the same subcellular compartment tend to cluster across the full filtered ERC network. Using the GO annotation table (HOG_AT_GO_Table.tsv), each HOG was assigned a CC category under the following rules: Plastid if all CC terms mapped to plastid compartments (GO:0009536 plastid; GO:0009507 chloroplast; GO:0009579 thylakoid), Mitochondrion if all CC terms mapped to the mitochondrion (GO:0005739), Nucleus if all CC terms mapped to the nucleus (GO:0005634) or nucleolus (GO:0005730), and Other for all remaining cases (no CC terms, additional compartments such as cytosol/ER/peroxisome, or mixed assignments spanning multiple categories). Only HOGs whose CC terms mapped exclusively to a single category were placed in that category; mixed CC annotations were grouped as Other.

We quantified subcellular clustering using Newman’s categorical assortativity coefficient, r, computed on the undirected ERC network with nodes restricted to the four CC categories above (Newman, 2003). Statistical significance was assessed by permutation: we randomly shuffled node labels 10,000 times while preserving network topology and category counts to obtain an empirical P-value (right-tailed test for homophily). As a sensitivity analysis, we repeated the test on the subgraph excluding nodes labeled Other (i.e., retaining only Plastid, Mitochondrion, and Nucleus).

### Selection analysis: HyPhy aBSREL

To infer differences in selection strength across the phylogeny for each HOG, we applied HyPhy’s adaptive Branch-Site Random Effects Likelihood (aBSREL) model (v2.5) (Smith et al., 2015). For each HOG, we provided (i) a codon alignment derived from the corresponding CDS sequences and (ii) a reconciled gene tree inferred with GeneRax (Morel et al., 2020). We used GeneRax instead of our GTST_reconciliation.py script because its Newick output and branch labeling were fully compatible with HyPhy’s input requirements. Codon alignments were generated using PAL2NAL (v14) based on MAFFT protein alignments (Rozewicki et al., 2019; Suyama et al., 2006). Alignments containing frame errors, internal stop codons, or insufficient post-trimmed codon length were excluded.

For every branch in each HOG, aBSREL fits a mixture of nonsynonymous/synonymous rate ratios (ω = dN/dS) across sites and performs a likelihood-ratio test (LRT) comparing a null model (no sites with ω > 1 on that branch) to an alternative model that allows a fraction of sites with ω > 1. P-values were corrected across branches within each HOG using HyPhy’s default multiple-testing procedure (Holm–Bonferroni). Branches were called as experiencing episodic diversifying selection when the adjusted P was < 0.05.

Because some species have multiple paralogous copies within a HOG, we summarized selection per terminal lineage by averaging ω across paralogous terminal branches that passed quality control. To limit inflation from artifactual or poorly estimated rates (e.g., alignment issues or extremely low synonymous rates), we applied two filters to terminal branches before averaging: (i) drop any branch with overall ω > 100; and (ii) if overall ω > 10, retain the branch only if its multiple-testing-corrected P was < 0.05. The resulting per-species mean ω were then used in downstream comparisons of selection intensity among accelerated and conserved HOGs in each carnivorous lineage.

For each species, we compared per-HOG mean ω (dN/dS) among three categories—Accelerated, Conserved, and Other—defined from our ERC BL Z-score analysis. Because ω values were strongly right-skewed and heteroskedastic, we used pairwise Mann–Whitney U tests (two-sided) as our primary, distribution-free comparison. To provide a robust empirical significance measure, we also performed a label-permutation test (10,000 replicates) on the difference in group medians. Effect sizes were quantified using Cliff’s δ, which estimates the probability that a randomly selected value from one group exceeds a randomly selected value from another. P-values from both tests were adjusted for multiple pairwise comparisons within each species using the Benjamini–Hochberg FDR procedure, with FDR < 0.05 considered statistically significant.

### Selection analysis: HyPhy RELAX

To assess whether carnivorous lineages experienced systematic shifts in selection strength, we applied HyPhy’s RELAX model (v.4.5) to each HOG using the same codon alignments and GeneRax-reconciled trees as above (Wertheim et al., 2015). RELAX partitions branches into test (focal carnivorous lineage/clade branches) and reference (all remaining non-CP branches) sets, fits a three–site-class distribution of ω values on the reference set, and estimates a selection-intensity parameter K that scales this distribution on the test set. Values K>1 indicate intensification (purifying and positive-selection classes move farther from neutrality), K<1 indicate relaxation (classes collapse toward neutrality), and K=1 indicates no change. Model support is evaluated by a likelihood-ratio test of K=1 versus K≠1; we controlled the FDR across HOGs using Benjamini–Hochberg (significant at FDR < 0.05).

To assess whether accelerated and conserved gene sets differ in selection intensity, we compared RELAX K values among Z-scores defined categories using both log-transformed and original (non-transformed) K data. Because K values were not normally distributed, we used nonparametric tests throughout. Overall variation among categories was assessed using the Kruskal–Wallis test. Pairwise differences were then evaluated using two-sided Mann– Whitney U tests, with Holm correction to control the family-wise error rate; Cliff’s δ was reported as a nonparametric effect size for each comparison. To test for categorical differences in the direction of selection (intensified vs. relaxed), we summarized counts of genes with K > 1 and K < 1 for each gene set and evaluated independence between category and direction using both chi-square and Fisher’s exact tests. For these contingency analyses, we also reported the test statistic, degrees of freedom, p-value, and Cramer’s V as a measure of association strength.

### Use of generative artificial intelligence

Generative AI tools were used in limited ways in this study: ChatGPT (OpenAI; accessed via web interface, 2024– 2025) during the initial analyses and manuscript preparation, and Claude (Anthropic; accessed via the Claude desktop application, 2025–2026) during revision. Their use fell into three categories.

i. Illustrations. Color drawings of representative CP species were generated with a generative image model based on photographs of living plants. The prompts specified key morphological features, and all images were manually checked and edited for biological accuracy before inclusion in the figures.
ii. Language editing. ChatGPT was used for grammar, spelling, and minor stylistic editing of text drafted by the authors. Multiple AI-edited versions were compared against the original text, and any suggested changes that altered scientific meaning or interpretation were rejected. Conceptual development, study design, data interpretation, and all substantive writing were performed by the authors.
iii. Script drafting. ChatGPT was used to generate initial versions of custom Python and R scripts from natural-language descriptions of the desired file-processing logic and example input/output formats. All AI-generated code was manually reviewed, iteratively corrected (with and without AI assistance), and validated using test datasets before being used in analyses (Supplementary Figure 24). Final scripts and command lines are provided in the associated GitHub repository.

## Data Availability Statement

All custom scripts and example bioinformatic command lines used in this study are available in the GitHub repository cp-erc-analysis (https://github.com/sukuanliu/cp-erc-analysis). Processed data and analysis outputs, including BL and Z-score matrices, ERC edge lists and assortativity results, GO enrichment tables and GO term frequency heatmaps, HyPhy aBSREL and RELAX summary files, and the Cytoscape ERC network session file, are deposited on Zenodo (DOI: 10.5281/zenodo.17806961). No new raw sequencing data were generated for this study; all genome resources are from previously published datasets listed in Supplementary Table A.

## Acknowledgements

This work was supported by grants from the National Science Foundation (IOS-2114641 and MCB-2322154). We thank the two anonymous reviewers for their valuable comments and suggestions, which improved the readability of the manuscript, the rigor of our analyses, and the clarity and transparency of our methods and data reporting.

---

**Supplementary Figure 1.**
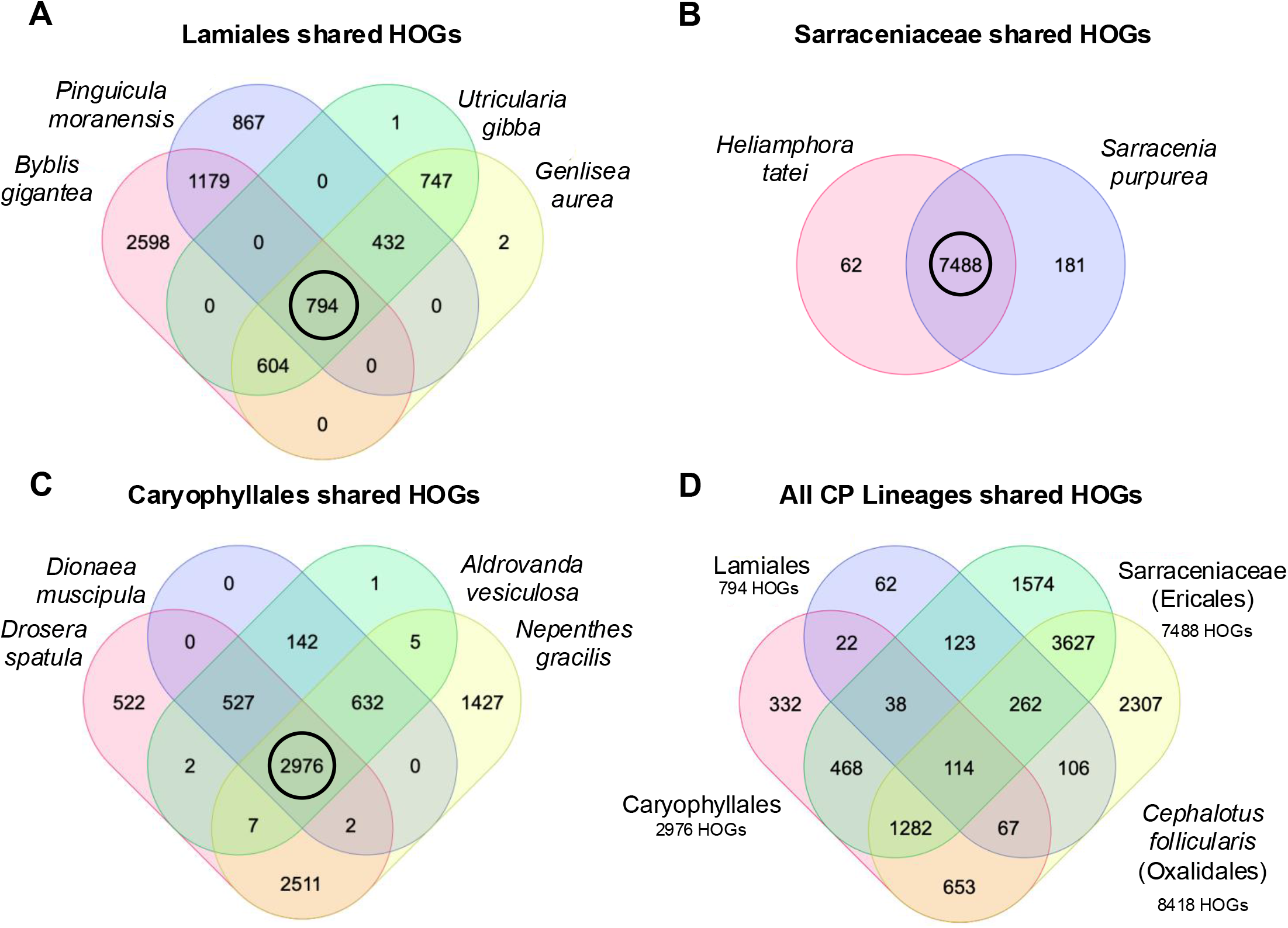
Venn diagrams showing the overlap of HOGs (with available Z-score estimates) among sampled carnivorous plant taxa. **(A)** Shared HOGs among Lamiales carnivores: *Byblis gigantea* (Byblidaceae), *Pinguicula caudata*, *Utricularia gibba*, and *Genlisea aurea* (Lentibulariaceae). **(B)** Shared HOGs between the two Sarraceniaceae (Ericales) pitcher plants: *Heliamphora tatei* and *Sarracenia purpurea*. **(C)** Shared HOGs among Caryophyllales carnivores: *Nepenthes gracilis* (Nepenthaceae), *Drosera spatulata*, *Dionaea muscipula*, and *Aldrovanda vesiculosa* (Droseraceae). **(D)** Shared HOGs across all carnivorous plant lineages,

**Supplementary Figure 2.**
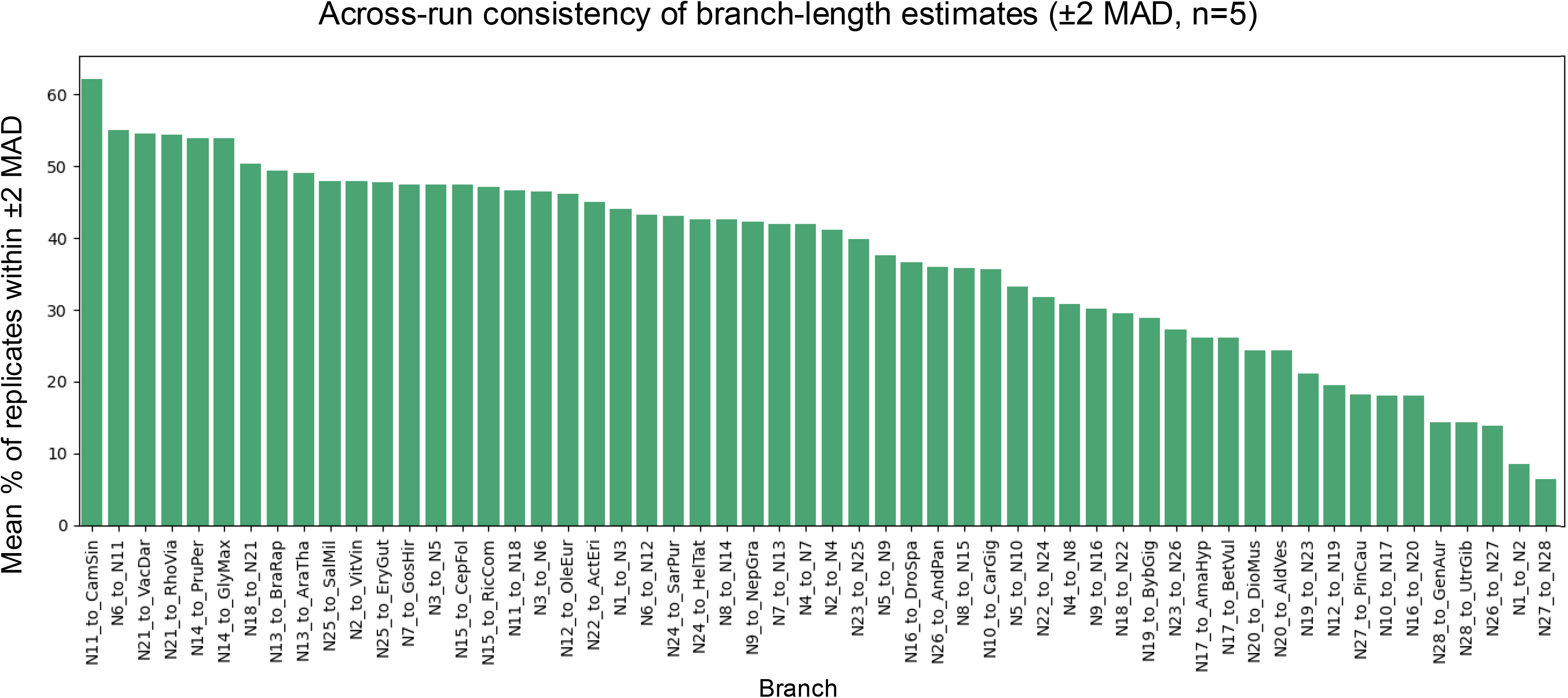
Across-run consistency of BL estimates by species-tree branch. For each HOG × branch, we computed the fraction of the five ERCnet replicate BL estimates that lay within ±2 MAD of the HOG-specific median; the plotted value is the mean of these fractions across HOGs for the given branch. Larger bars denote more consistent BL estimates across runs. Branch labels correspond to internal and terminal node names in the phylogeny shown in Figure 1.

**Supplementary Figure 3.**
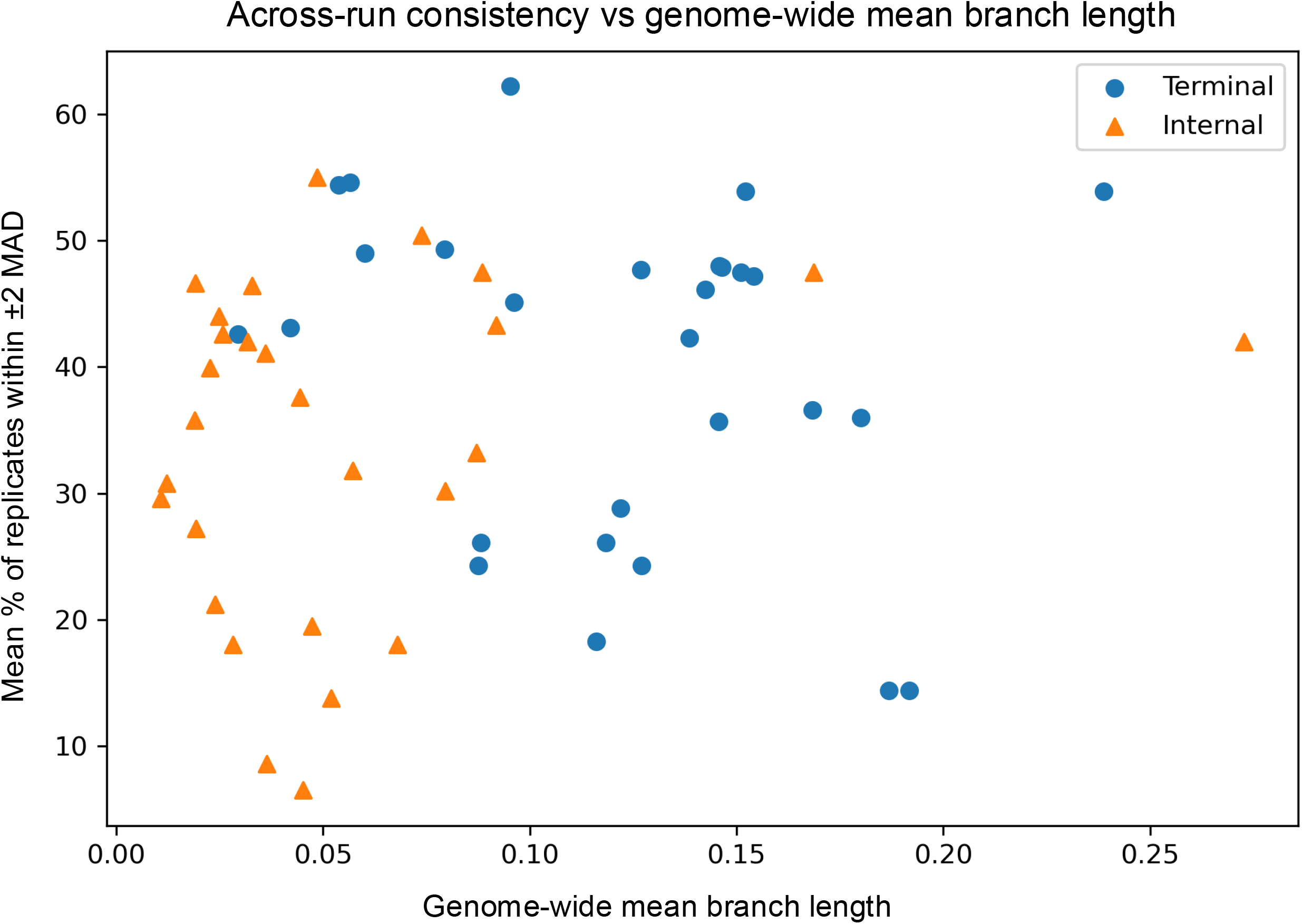
Across-run consistency of BL estimates versus genome-wide BL; internal nodes (triangles) and terminal nodes (circles). Internal branches had significant shorter genome-wide average BL than terminals (means 0.056 vs 0.123; Welch’s t-test p < 0.001). Across-run consistency (mean % of replicates within ±2 MAD) was marginally lower for internal branches (33.9% vs 40.0%; p = 0.09).

**Supplementary Figure 4.**
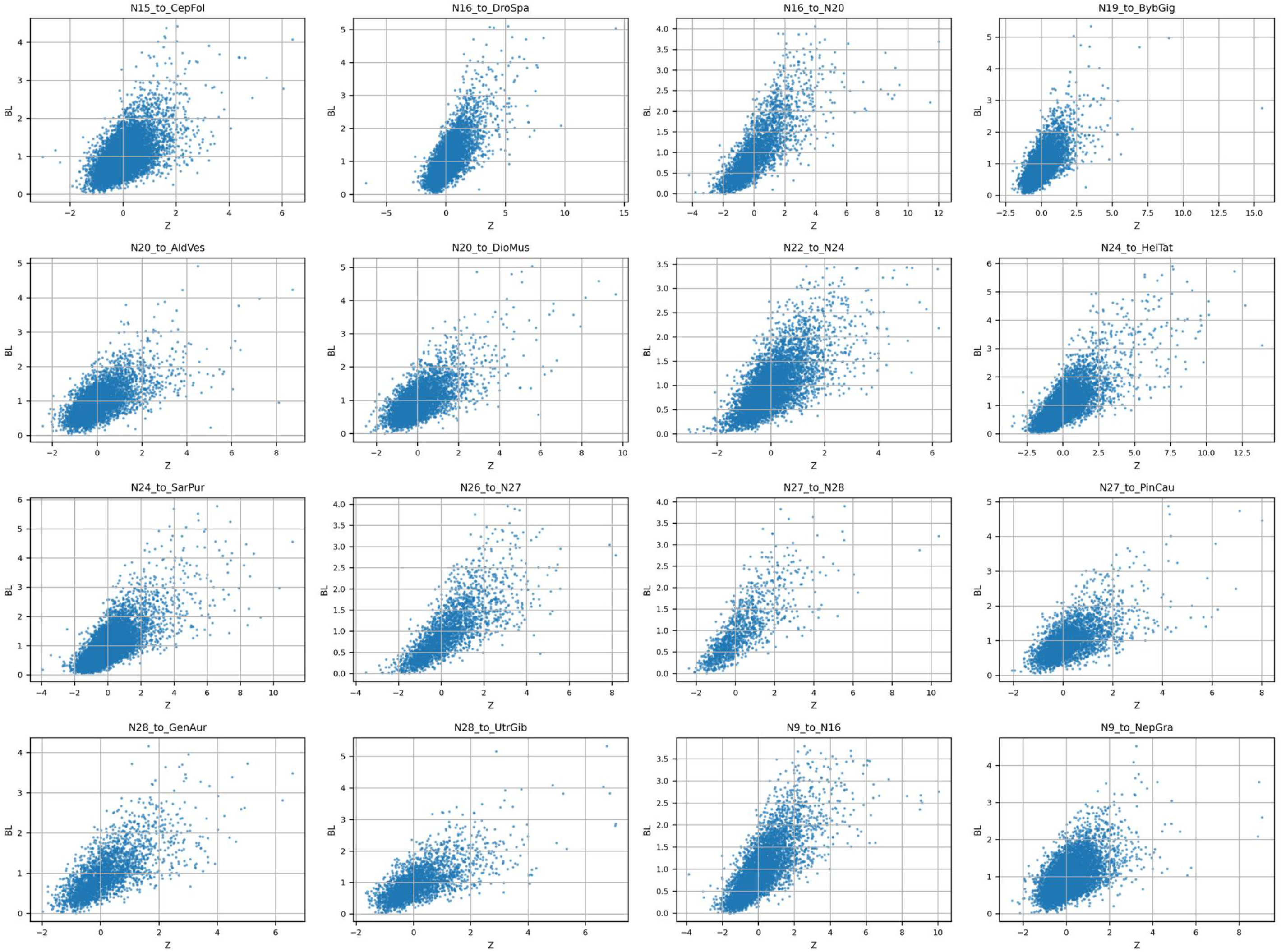
Normalized Branch Length vs. Z score among CP internal and terminal branches.

**Supplementary Figure 5.**
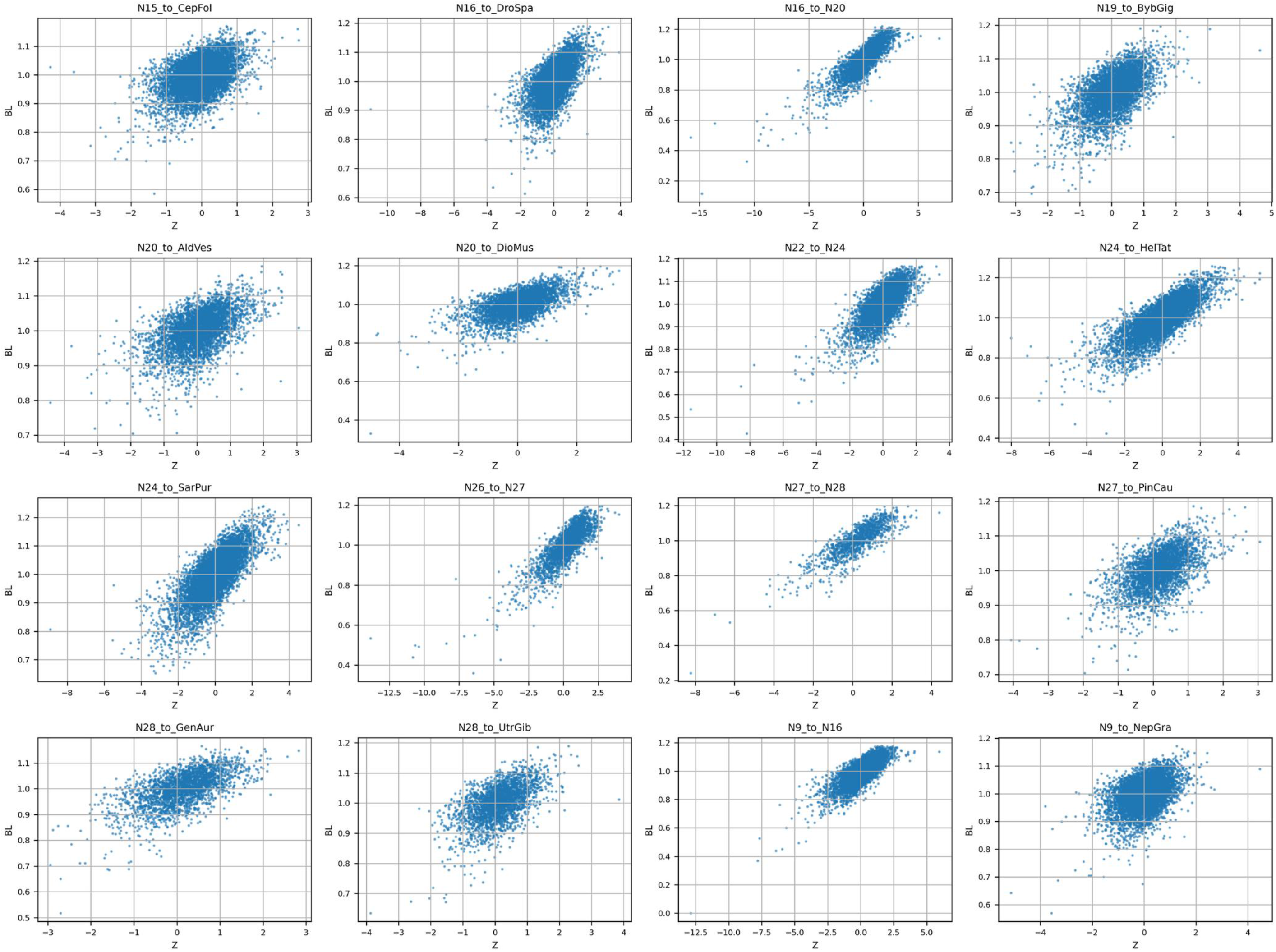
Log Normalized Branch Length vs. Z score among CP internal and terminal branches.

**Supplementary Figure 6.**
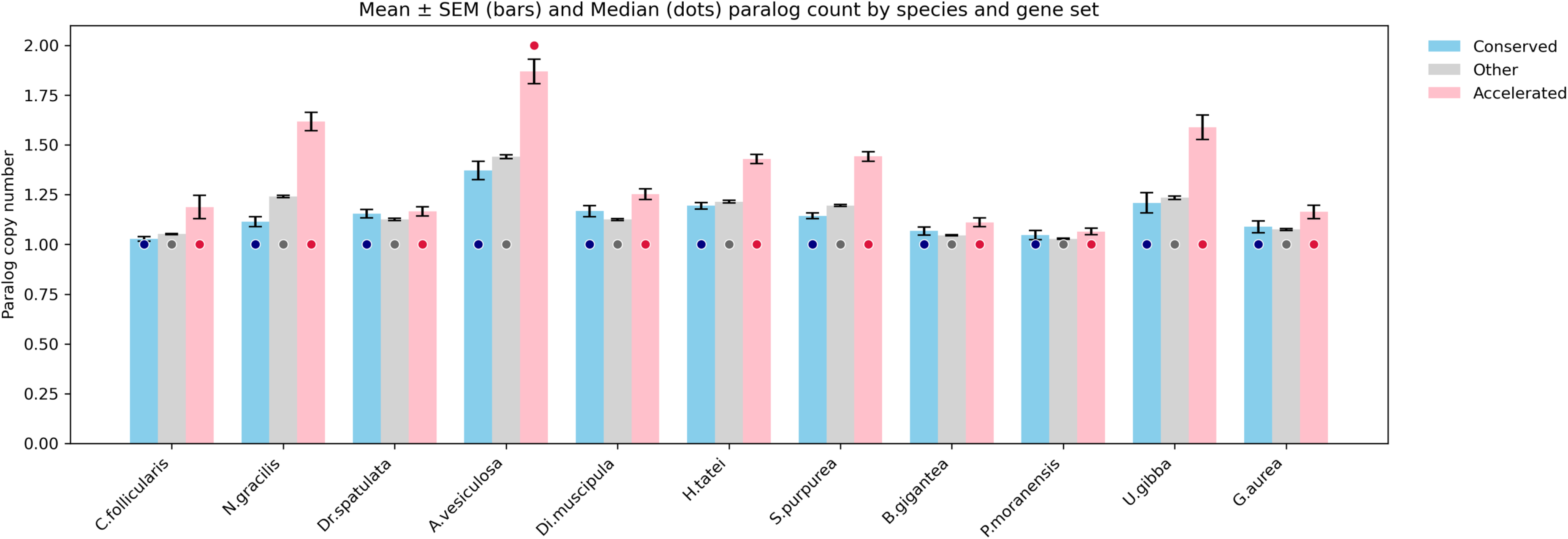
**Mean ± SEM (bars) and median (dots) paralog copy number** for accelerated (pink), conserved (blue), and background (grey, labeled as “other”) genes across carnivorous plant genomes. Accelerated genes generally exhibit higher copy numbers than conserved and background genes in most lineages, consistent with widespread gene family expansion associated with evolutionary rate acceler ation.

**Supplementary Figure 7.**
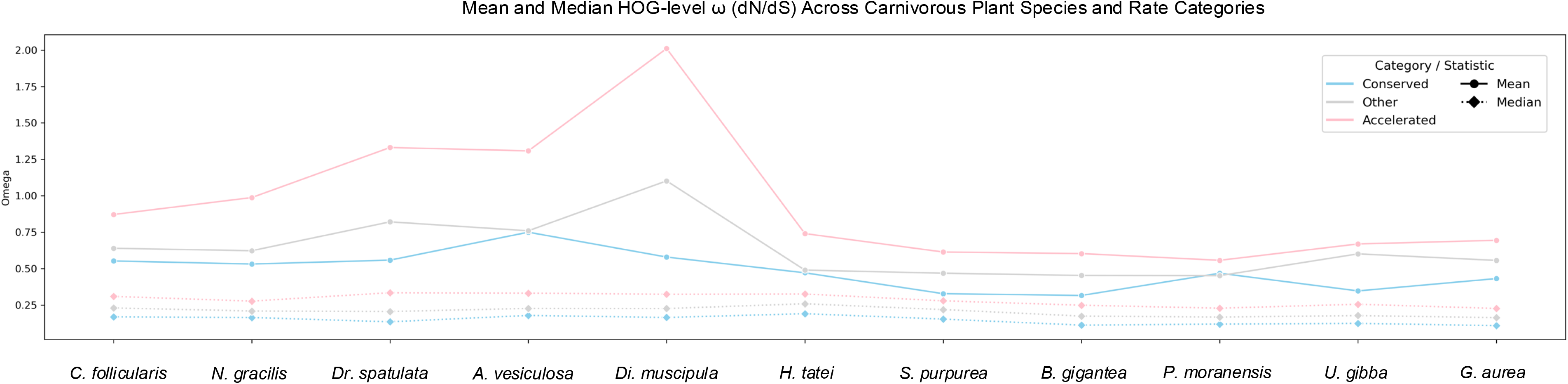
Mean and median HOG-level ω (dN/dS) for three rate categories (Accelerated, Conserved, and Other) across 11 carnivorous plant species. Solid lines (circles) show the mean ω, while dotted lines (diamonds) show the median ω for each category within each species.

**Supplementary Figure 8.**
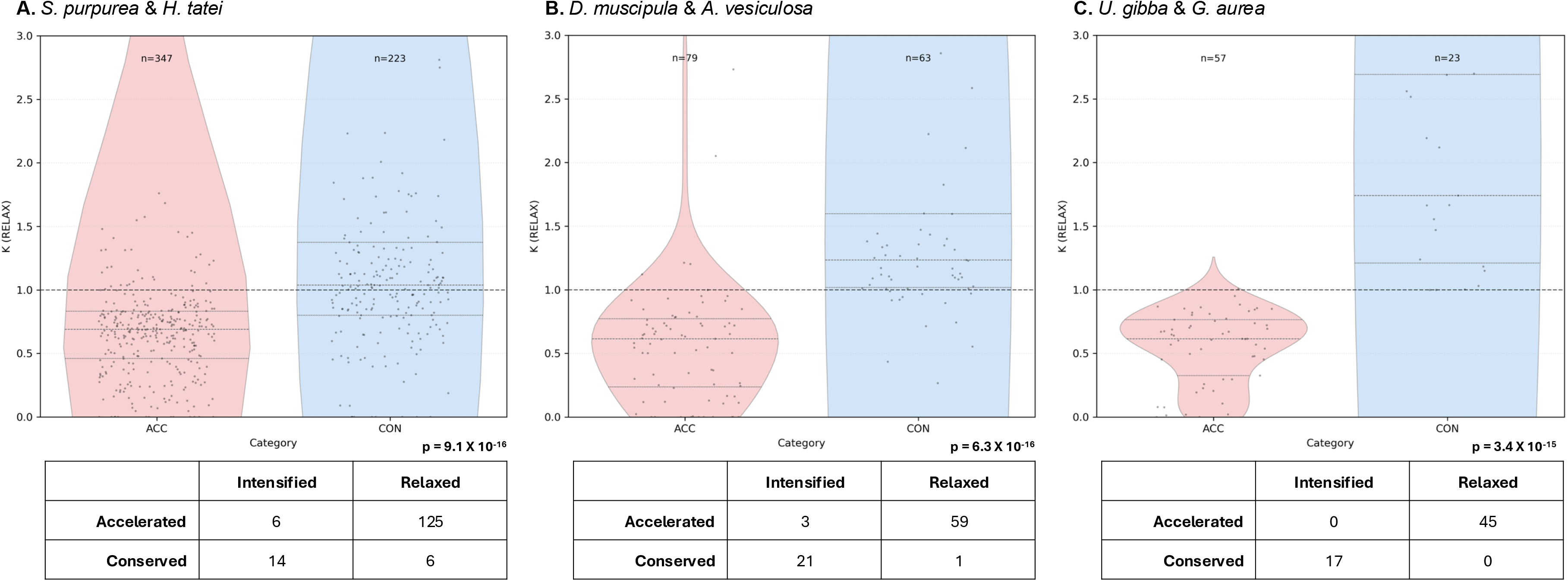
**Selection-intensity parameter (K, untransformed) by rate category** for homologous genes (HOGs) shared between representative carnivorous plant lineages. **(Top) Violin plots show the distribution of per-HOG selection-intensity values (K) for Accelerated (ACC) and Conserved (CON) gene sets**. **(Bottom) Corresponding contingency tables** summarize the numbers of significantly intensified and relaxed HOGs within each rate category, as used in chi-square tests. Results are shown for (A) *S. purpurea* and *H. tatei* (Sarraceniaceae), (B) *D. muscipula* and *A. vesiculosa* (Droseraceae), and (C) *U. gibba* and *G. aurea* (Lentibulariaceae). Points represent individual HOGs. Horizontal dotted lines indicate medians and interquartile ranges, while the dashed line marks K = 1, corresponding to no net change in selection intensity (values < 1 indicate relaxation; values > 1 indicate intensification). n denotes the number of HOGs per category. Reported p-values correspond to chi-square tests comparing the proportions of intensified versus relaxed HOGs between rate categories.

**Supplementary Figure 9.**
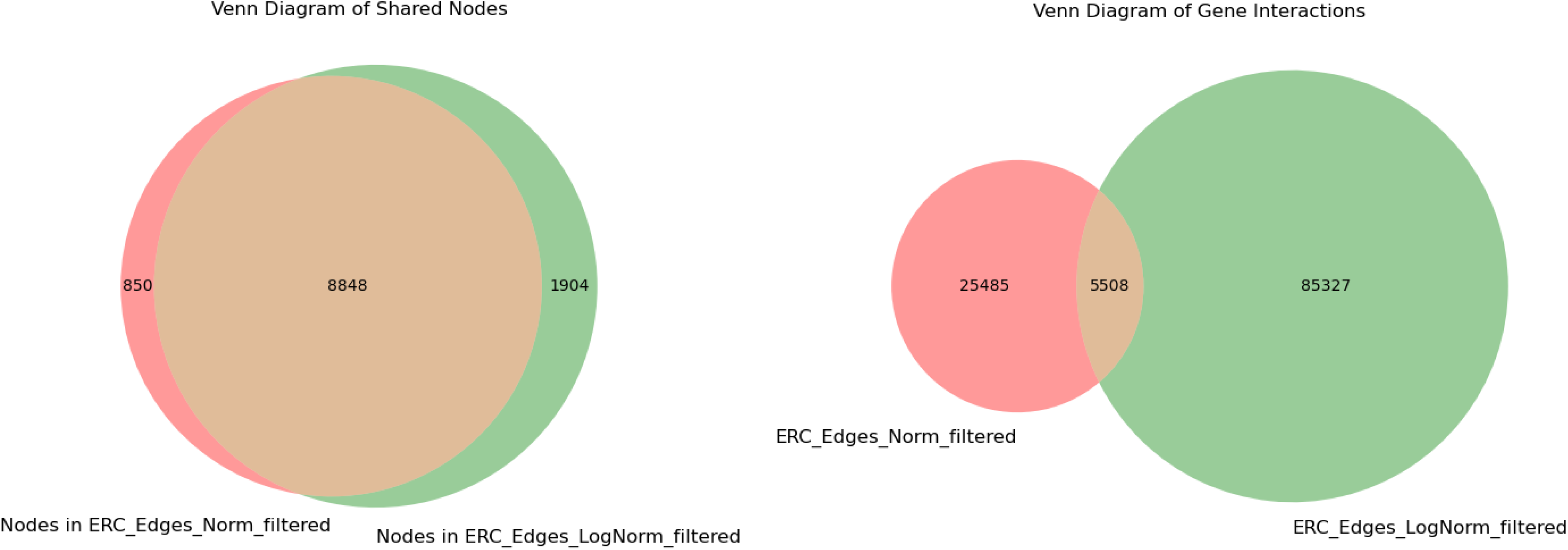
Venn diagrams showing shared nodes and edges from filtered ERC networks (slope (β) > 0.75, P < 1 × 10⁻⁴, and >10 overlapping branches, r^2>0.4) inferred from normalized BL and log normalized BL matrices.

**Supplementary Figure 10.**
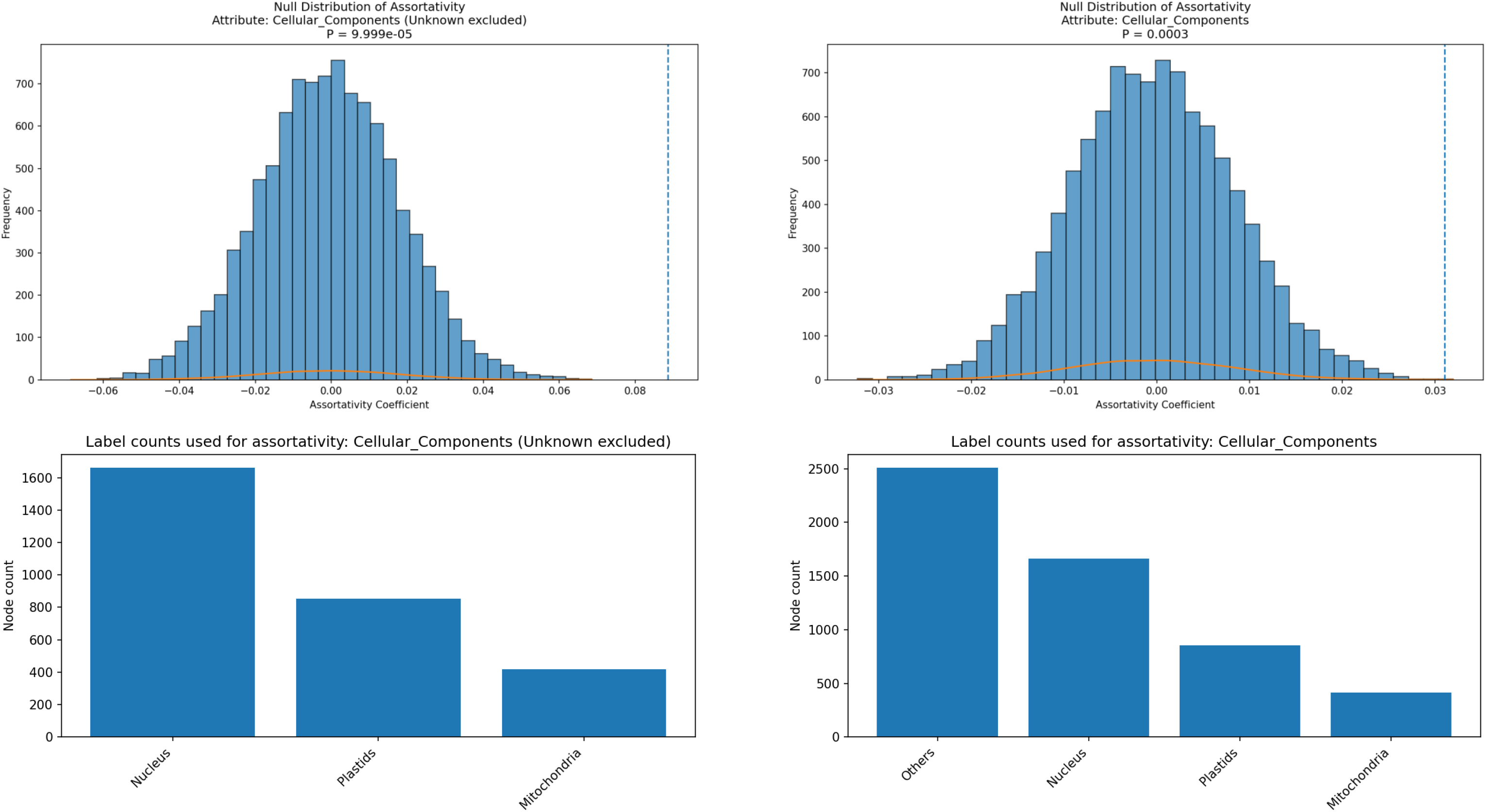
Subcellular-compartment assortativity in the shared ERC network. **Top:** Null distributions of Newman’s attribute assortativity coefficient (*r*) generated by 100,000 random permutations of node labels (network topology preserved). The dashed line marks the observed *r*. Left, excluding Other (three compartments: plastid, nucleus, mitochondrion): *r* = 0.089, *P* = 1 × 10⁻⁵. Right, including Other: *r* = 0.031, *P* = 3 × 10⁻⁴. **Bottom:** Node counts per compartment used in each test. Only nodes with GO Cellular Component annotations were included; edges are the intersection of significant ERC pairs from the normalized and log-normalized networks.

**Supplementary Figure 11.**
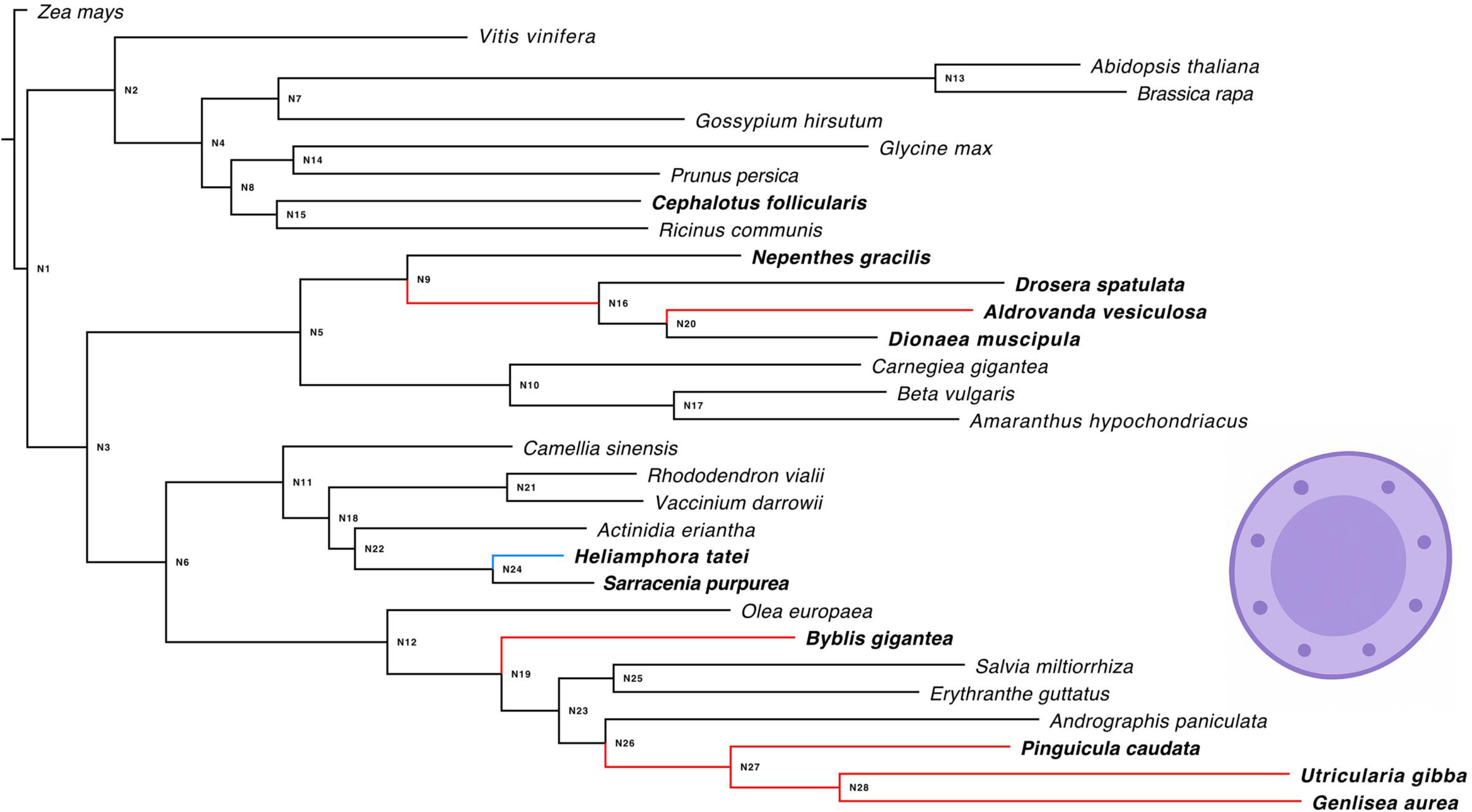
Phylogeny of sampled taxa showing evolutionary rate shifts in nuclear-localized genes associated with carnivorous plant lineages. Branches where genes localized to the nucleus exhibit significant acceleration relative to non-carnivorous plants are shown in red, whereas those showing significant conservation are shown in blue. Terminal branches represent extant species, while internal branches correspond to inferred ancestral nodes.

**Supplementary Figure 12.**
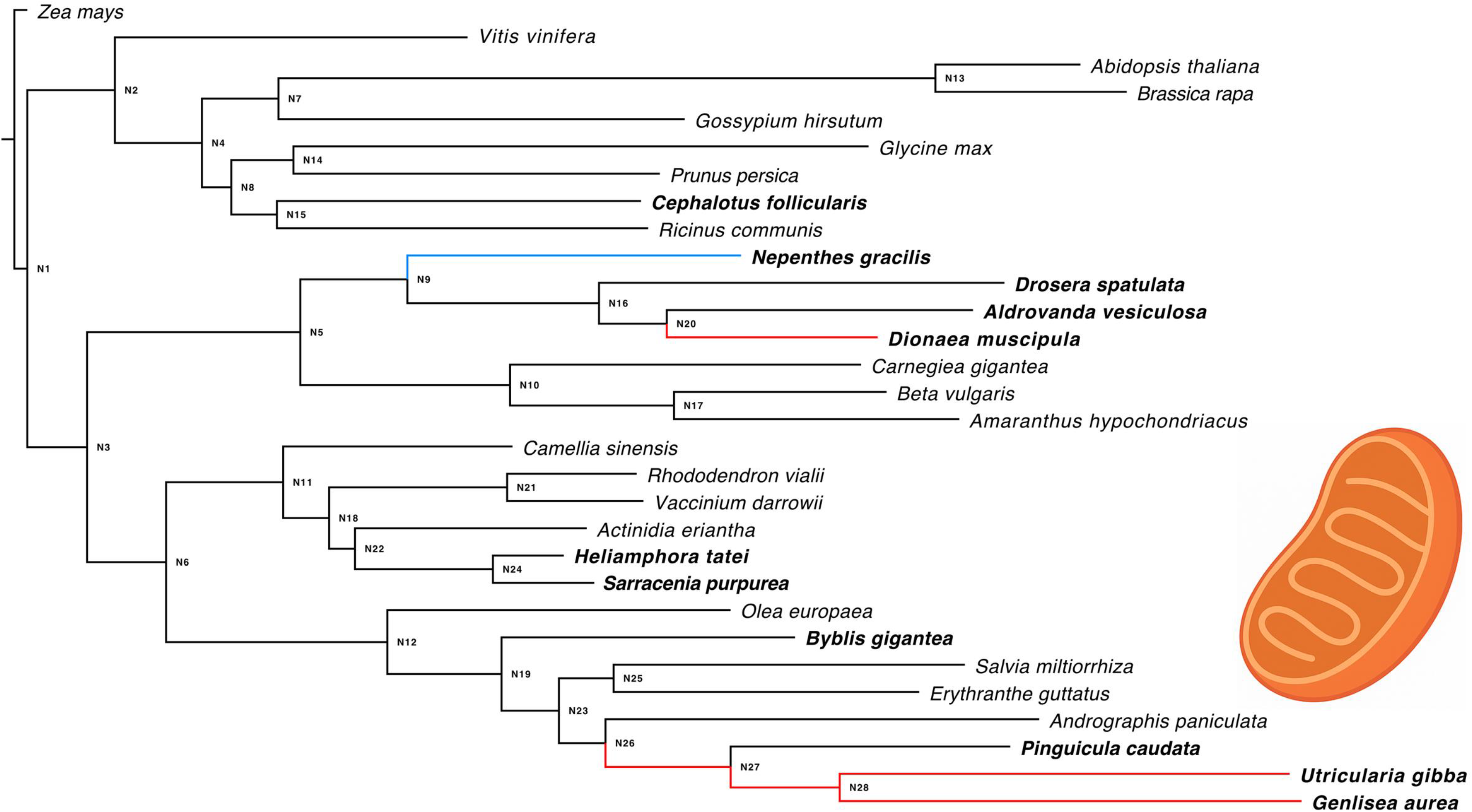
Phylogeny of sampled taxa showing evolutionary rate shifts in mitochondria-localized genes associated with carnivorous plant lineages. Branches where genes localized to the nucleus exhibit significant acceleration relative to non-carnivorous plants are shown in red, whereas those showing significant conservation are shown in blue. Terminal branches represent extant species, while internal branches correspond to inferred ancestral nodes.

**Supplementary Figure 13.**
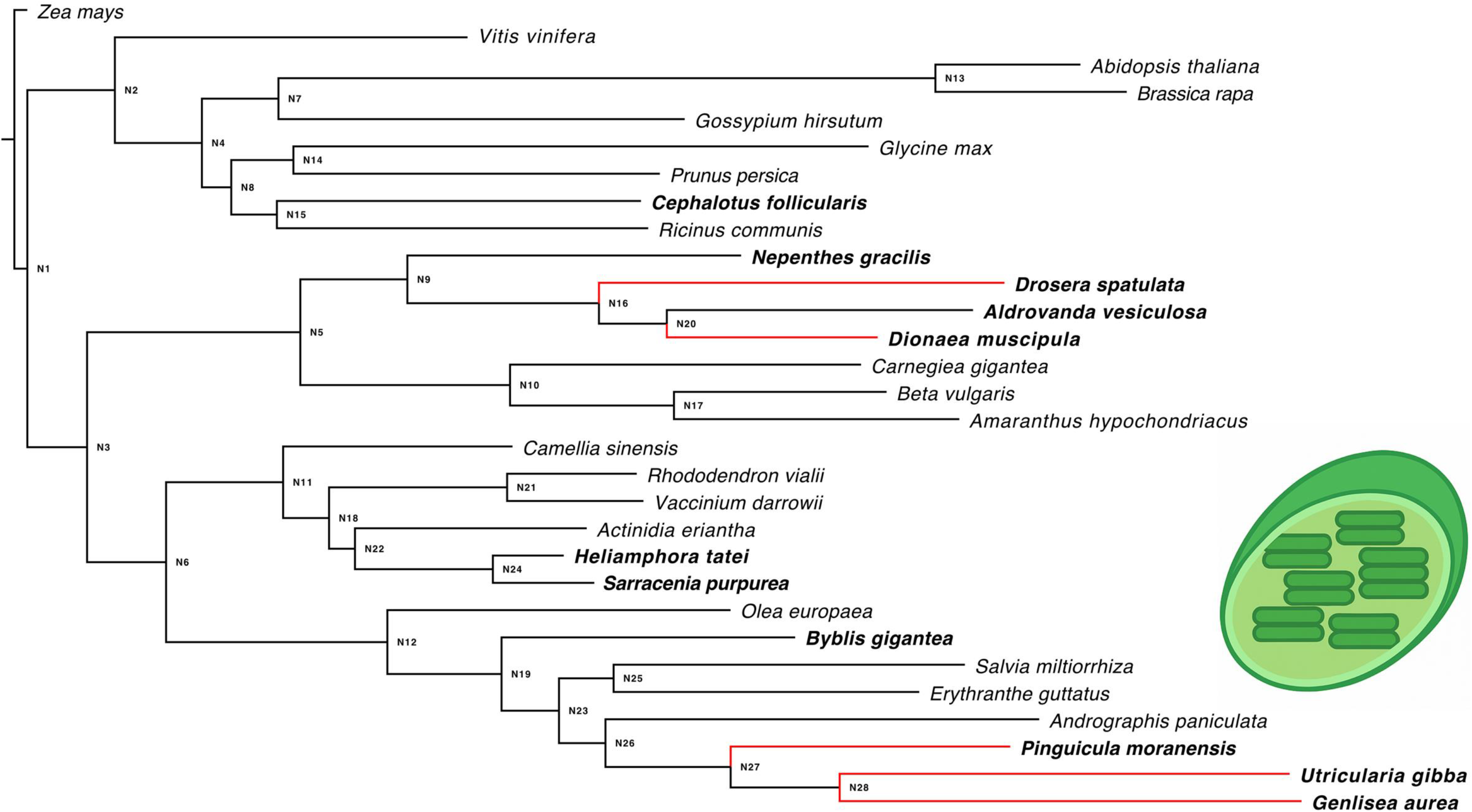
Phylogeny of sampled taxa showing evolutionary rate shifts in plastid-localized genes associated with carnivorous plant lineages. Branches where genes localized to the nucleus exhibit significant acceleration relative to non-carnivorous plants are shown in red, whereas those showing significant conservation are shown in blue. Terminal branches represent extant species, while internal branches correspond to inferred ancestral nodes.

**Supplementary Figure 14.**
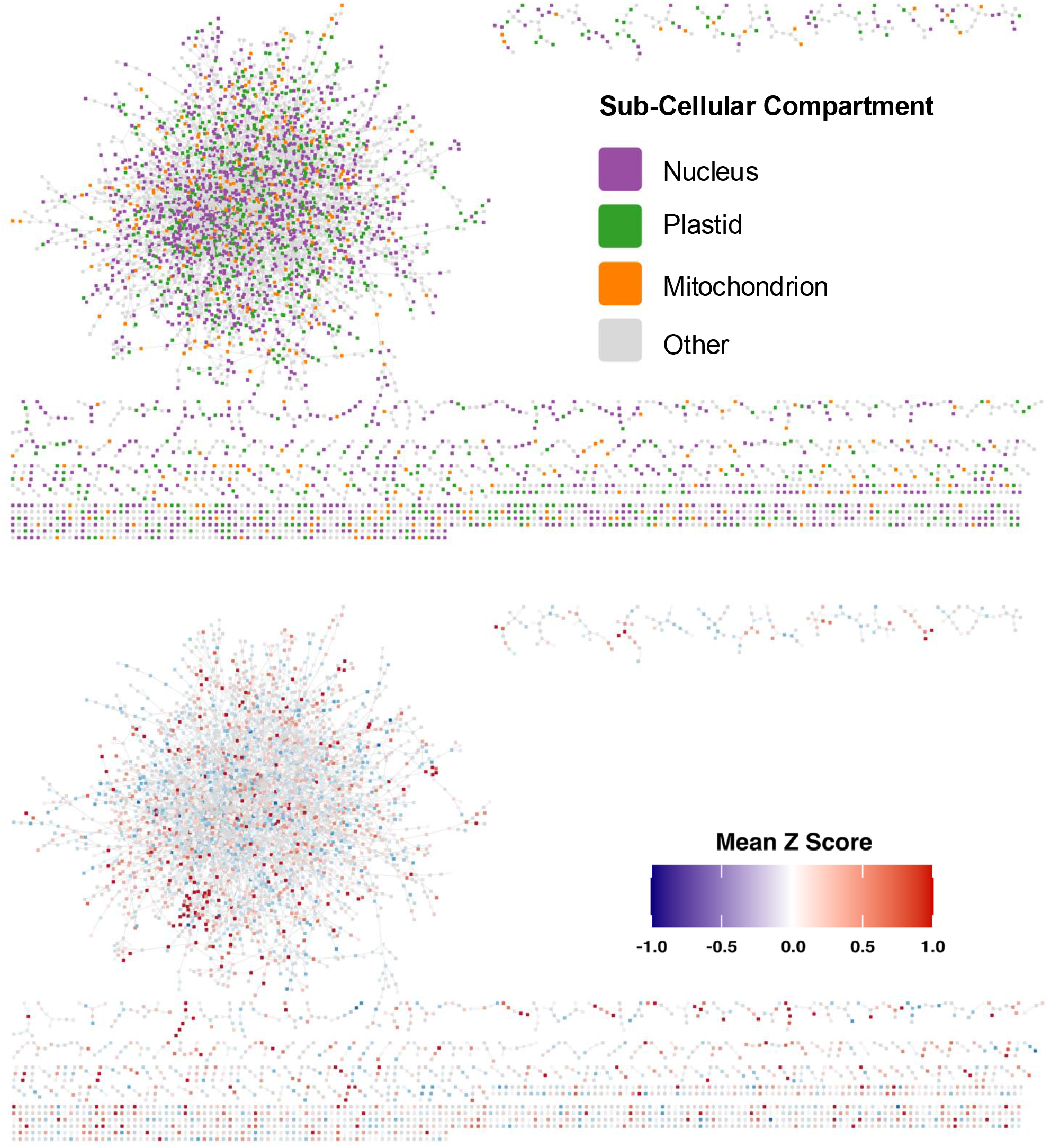
Shared ERC network colored by subcellular compartment (Top) and branch-length signal (Bottom). Nodes are HOGs; edges are ERC pairs significant in both the normalized and log-normalized analyses (β > 0.75, P < 10⁻⁴, >10 overlapping branches). Layout shown for the giant component with remaining connected components arranged below (Cytoscape v3.10.3). Top: nodes colored by GO Cellular Component category—nucleus (purple), plastid (green; includes chloroplast/thylakoid), mitochondrion (orange), and other (gray; mixed/absent CC terms). Bottom: same network with nodes colored by the mean Z-score of normalized branch-length difference between carnivorous and non-carnivorous lineages (blue = lower in carnivores, white ≈ 0, red = higher; scale −1 to +1).

**Supplementary Figure 15.**
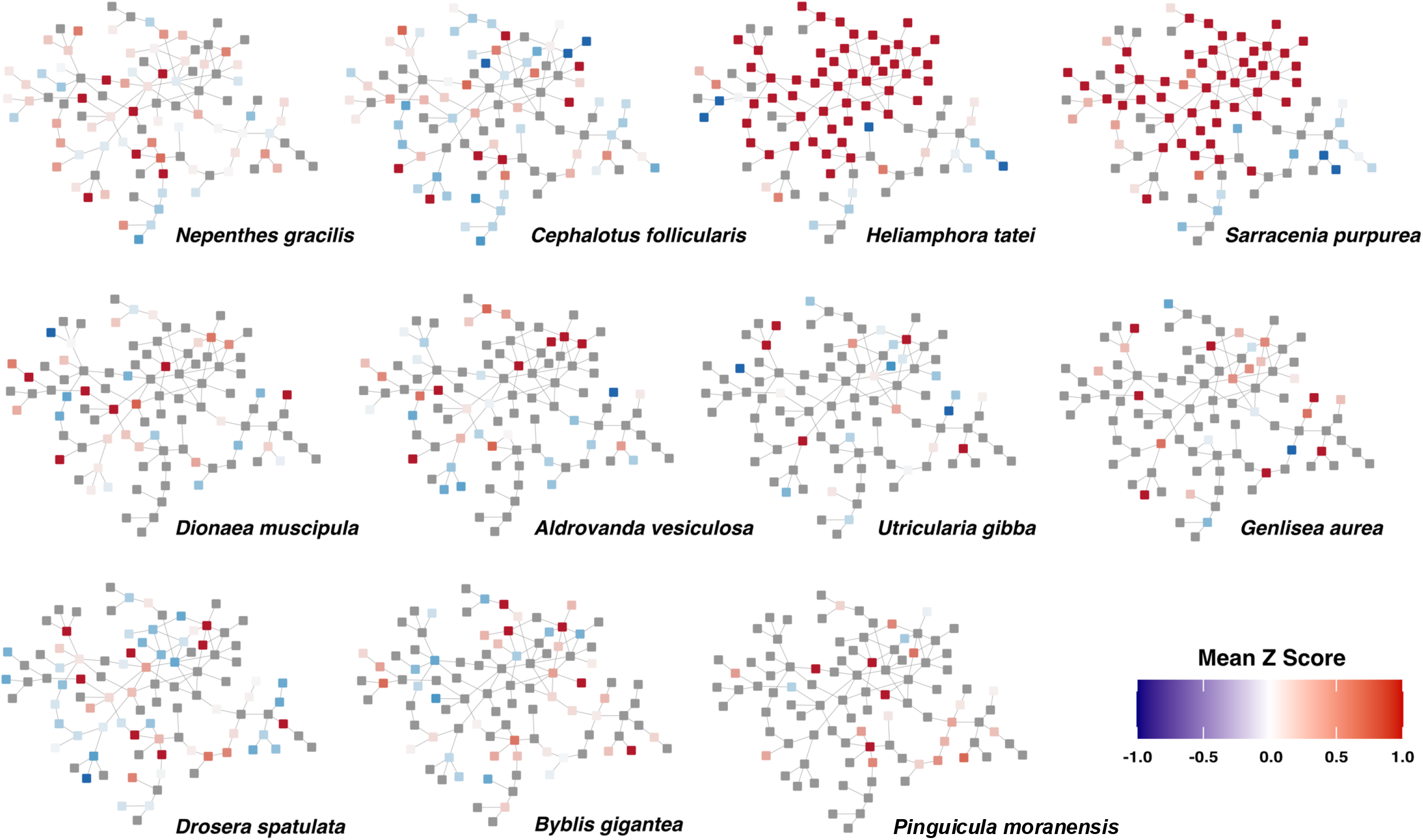
Z-scores (calculated from normalized branch lengths) for Community 10 from the shared ERC network, plotted by HOG and species ( terminal branches). Gray nodes indicate HOGs missing Z-scores for the corresponding species. Notably, the strongest acceleration signals are observed in the two sampled Sarraceniaceae species, highlighting lineage-specific rate shifts within this community. Of the 111 inferred ERC hits (edges) in this community, 75, 73, 51, 47, 46, 25, 16, 16, 8, 8, and 6 edges connected HOGs (nodes) present in the genomes of H. tatei, S. purpurea, N. gracilis, D. spatulata, C. follicularis, B. gigantea, D. muscipula, A. vesiculosa, G. aurea, U. gibba, and P. moranensis, respectively. Taken together, this pattern suggests that Community 10 reflects both lineage-specific acceleration in Sarraceniaceae and convergent acceleration across multiple, independently evolved CP lineages.

**Supplementary Figure 16.**
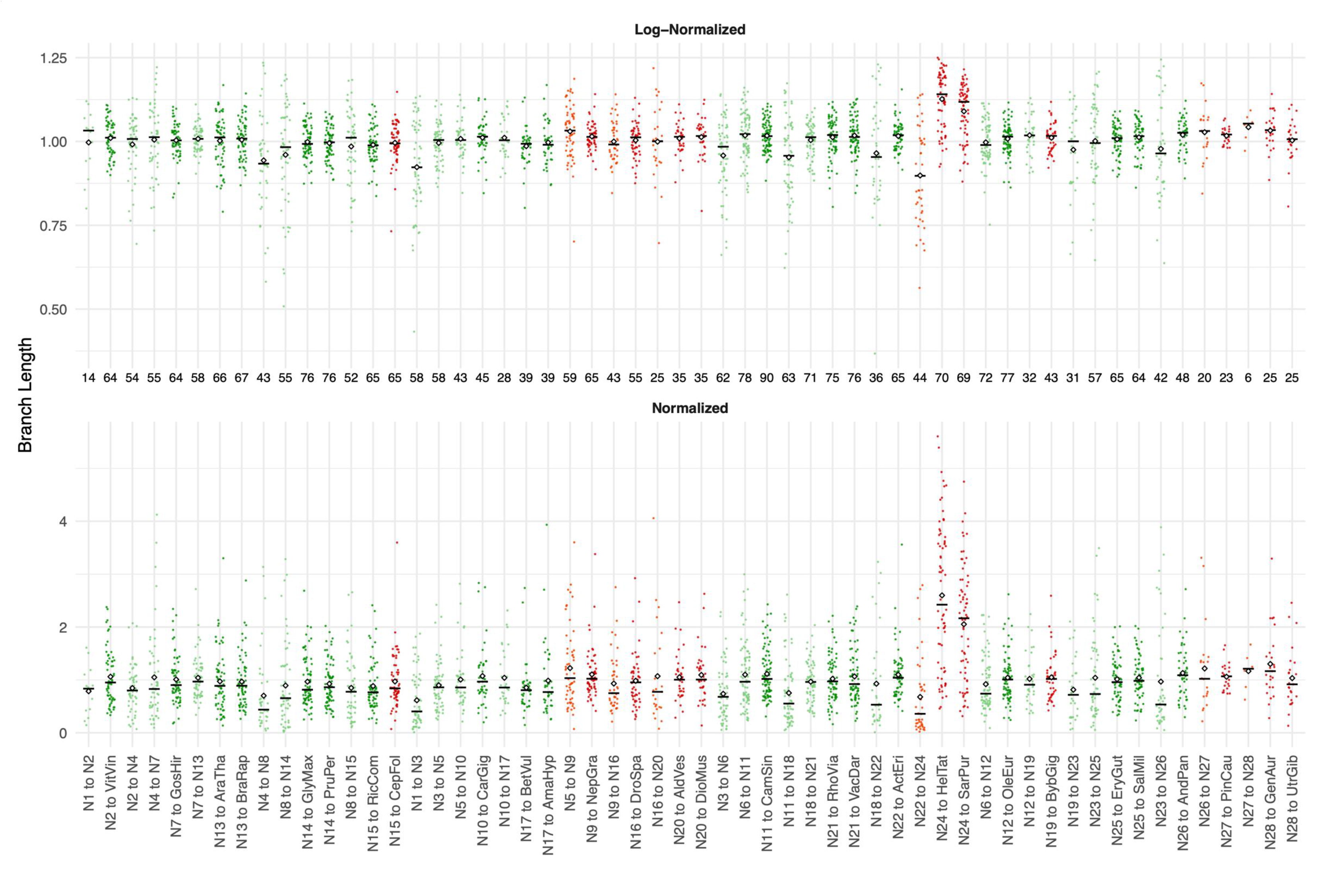
Distributions of HOG branch lengths for Community 10 in the log-normalized (top) and normalized (bottom) datasets. Non-carnivorous internal and terminal branches are shown in light and dark green, respectively, while carnivorous internal and terminal branches are shown in light and dark red. The number of branch-length estimates for each branch is indicated along the x-axis below the log-normalized plot, and branch labels (internal vs. terminal) are shown below the normalized plot.

**Supplementary Figure 17.**
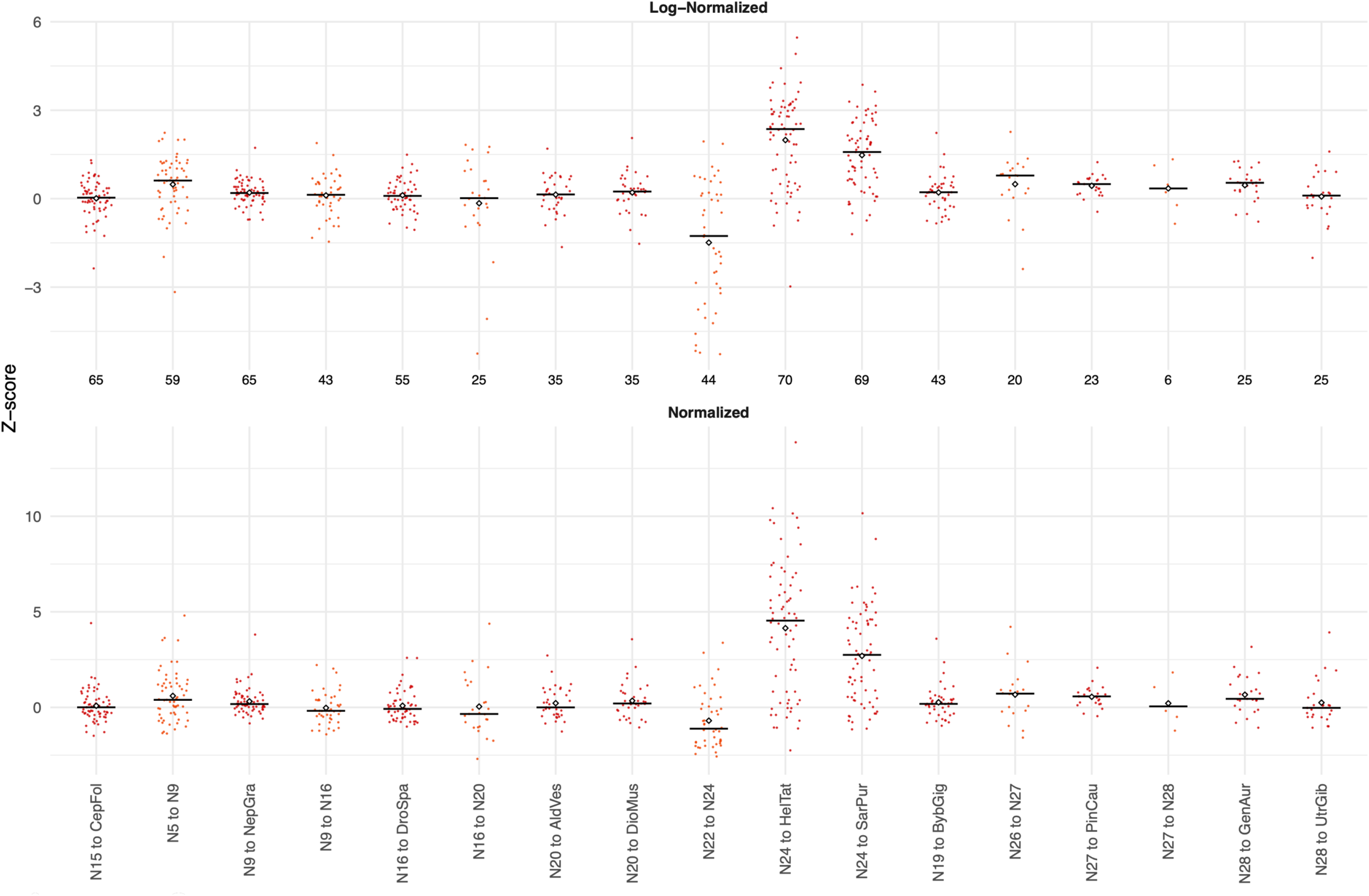
Distributions of HOG Z-scores for Community 10 in the log-normalized (top) and normalized (bottom) datasets. The number of Z-score estimates for each carnivorous branch is shown along the x-axis below the log-normalized plot, and branch labels are provided below the normalized plot. Positive Z-scores indicate relative rate acceleration, whereas negative Z-scores indicate relative conservation for each carnivorous branch. Notably, the two Sarraceniaceae branches (*H. tatei* and *S. purpurea*) show particularly strong positive Z-scores, consistent with pronounced lineage-specific acceleration in this community.

**Supplementary Figure 18.**
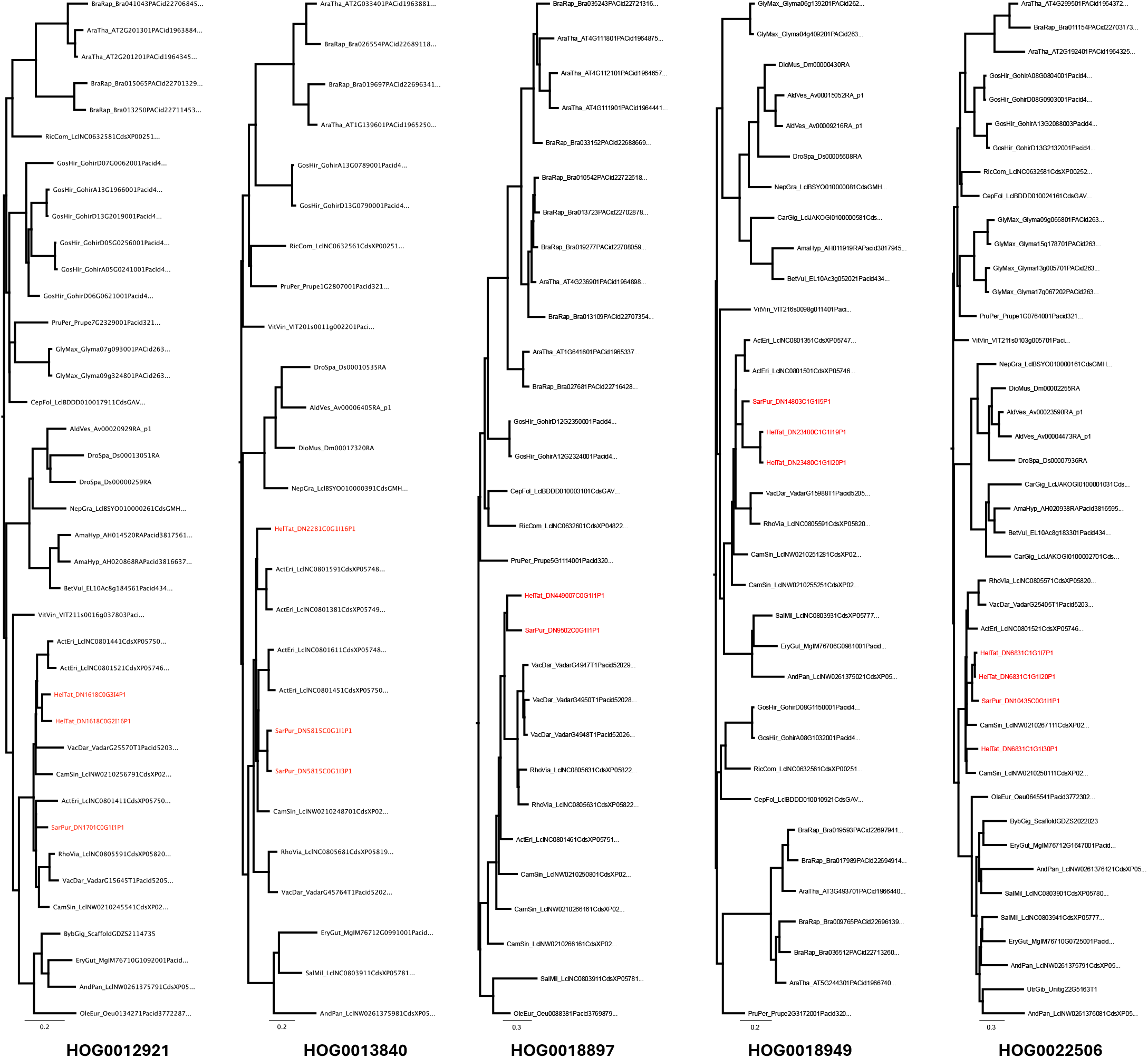
Examples of reconciled gene trees for selected HOGs within Community 10. Sarraceniaceae sequences are highlighted in red. These examples illustrate representative topologies showing lineage-specific clustering of *Heliamphora* and *Sarracenia* sequences, reflecting accelerated evolution and potential paralog retention in the Sarraceniaceae lineage.

**Supplementary Figure 19.**
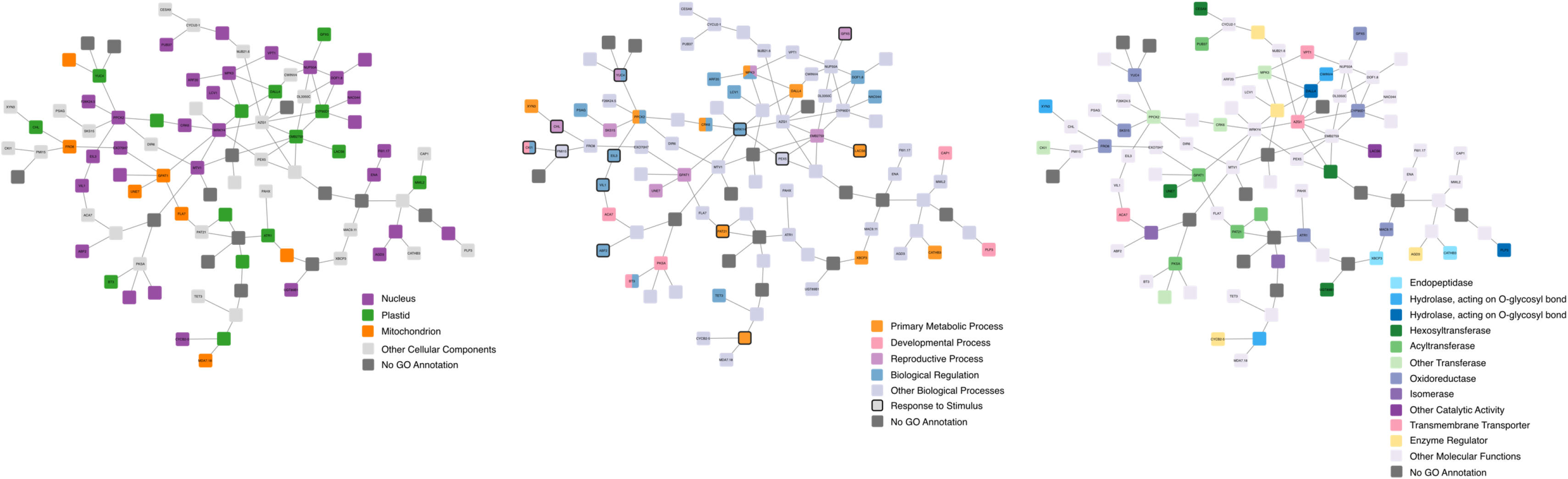
Network visualization of Community 10, highlighting (A) subcellular localization of gene products (major organelles), (B) enriched Biological Process GO terms, and (C) enriched Molecular Function GO terms.

**Supplementary Figure 20.**
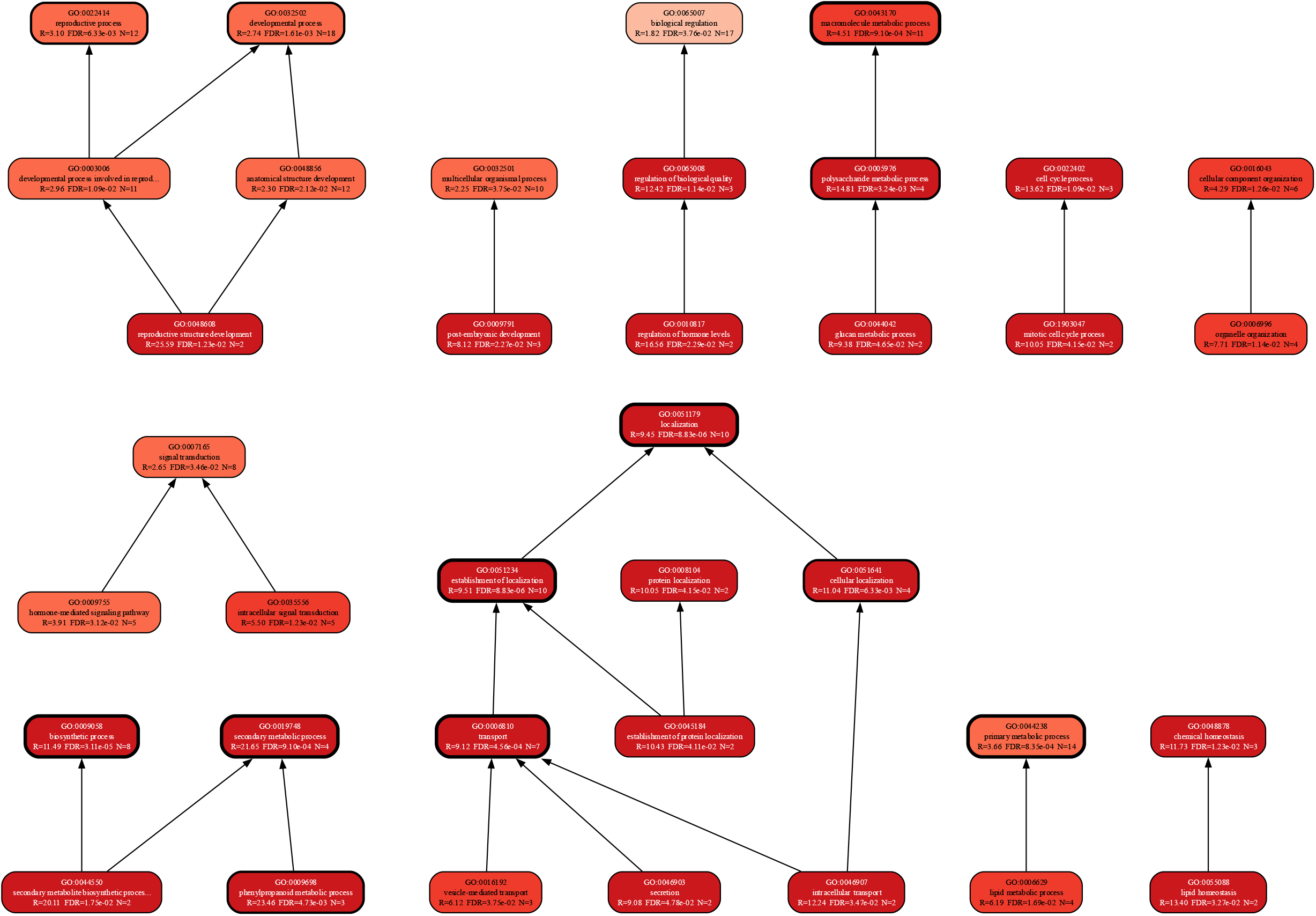
Enriched Biological Process GO terms for Community 10 from the shared ERC network. Terms are organized hierarchically according to the GO ontology, with arrows indicating parent–child relationships. Node shading reflects the enrichment ratio, and bold outlines denote statistically significant terms (FDR-adjusted P-values).

**Supplementary Figure 21.**
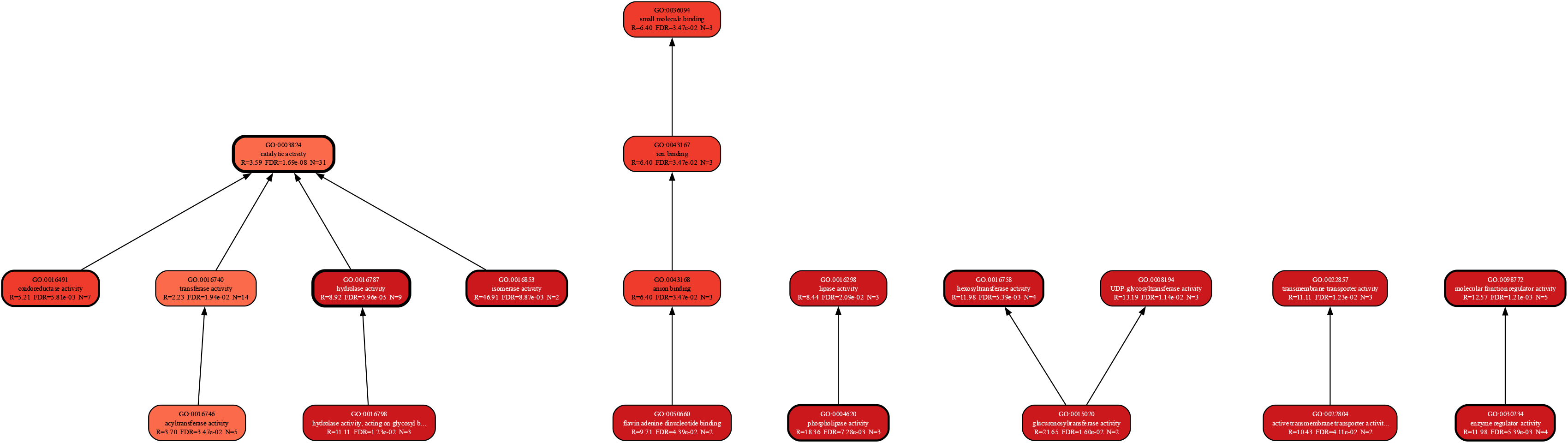
Enriched Molecular Function GO terms for Community 10 from the shared ERC network. Terms are organized hierarchically according to the GO ontology, with arrows indicating parent–child relationships. Node shading reflects the enrichment ratio, and bold outlines indicate statistically very significant terms (FDR-adjusted P-values).

**Supplementary Figure 22.**
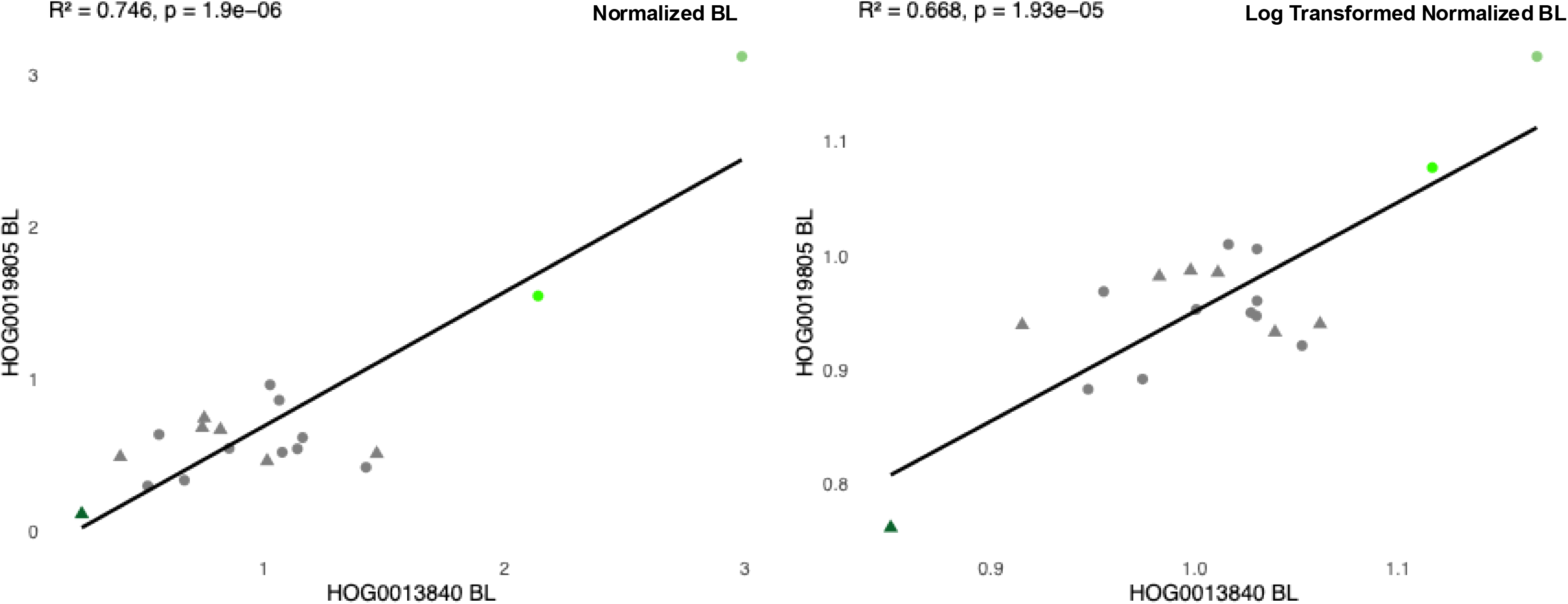
Example of a significant evolutionary rate covariation (ERC) hit between HOG0013840 (WRKY3/4) and HOG0019805 (PEX5). (A) ERC inferred from normalized branch lengths. (B) ERC inferred from log-transformed normalized branch lengths. Each point represents a species; circles correspond to terminal branches and triangles to internal branches. Carnivorous taxa (*H. tatei* and *S. purpurea*) are highlighted in darker and lighter green, respectively, whereas non-carnivorous taxa are shown in gray. Both analyses reveal a strong positive correlation (R² = 0.746 and 0.668, respectively; p < 2 × 10⁻⁵), indicating concordant evolutionary rate shifts across lineages. Notably, this gene pair also shows co-expression in Arabidopsis thaliana, suggesting that the observed ERC reflects a conserved functional association that predates the evolution of carnivory.

**Supplementary Figure 23.**
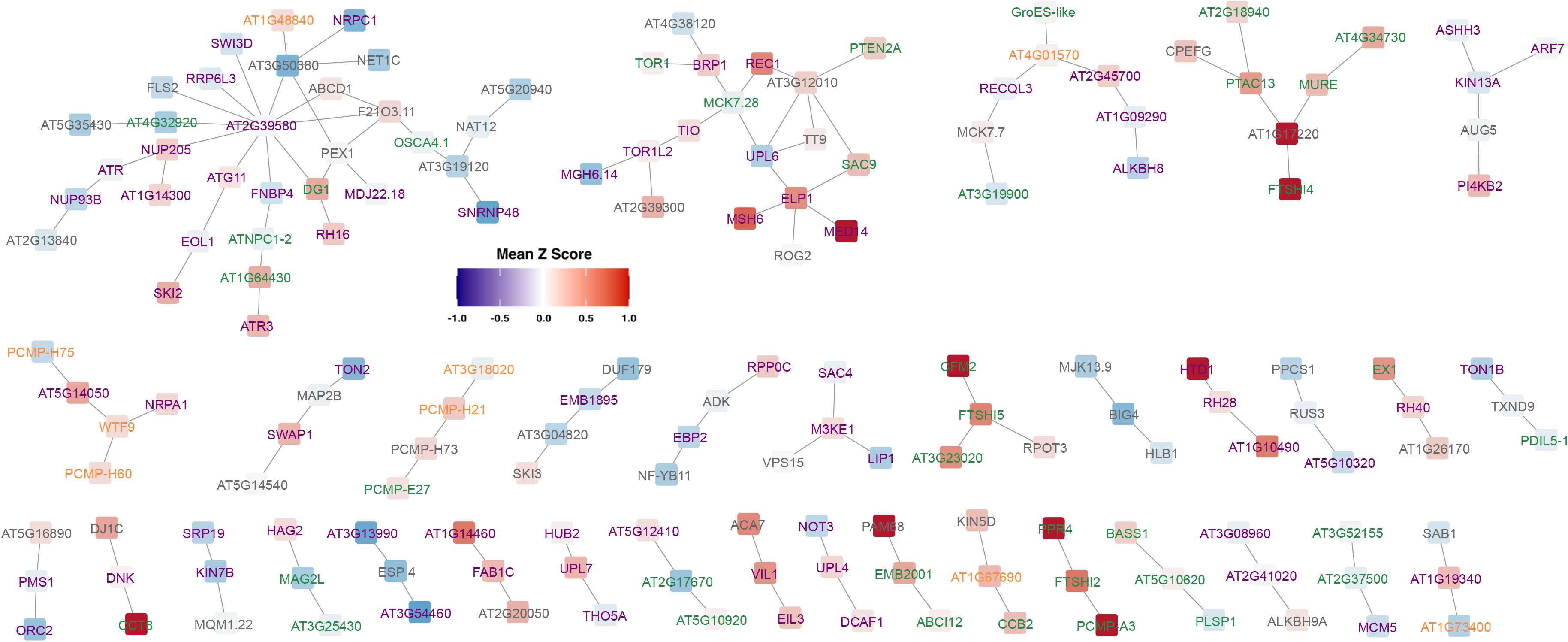
**Shared ERC network of gene clusters (n HOGs > 2) exhibiting significant positive co-expression (LS > 1.5)** in *Arabidopsis thaliana*. Nodes are colored by mean normalized CP BL Z-scores, with red indicating higher and blue indicating lower relative rates of evolution. Nodes lacking carnivorous plant Z-scores were filtered out. Node labels are colored by Gene Ontology (GO) Cellular Component category —nucleus (purple), plastid (green; including chloroplast and thylakoid), mitochondrion (orange), and other (gray; mixed or unassigned CC terms).

**Supplementary Figure 24.**
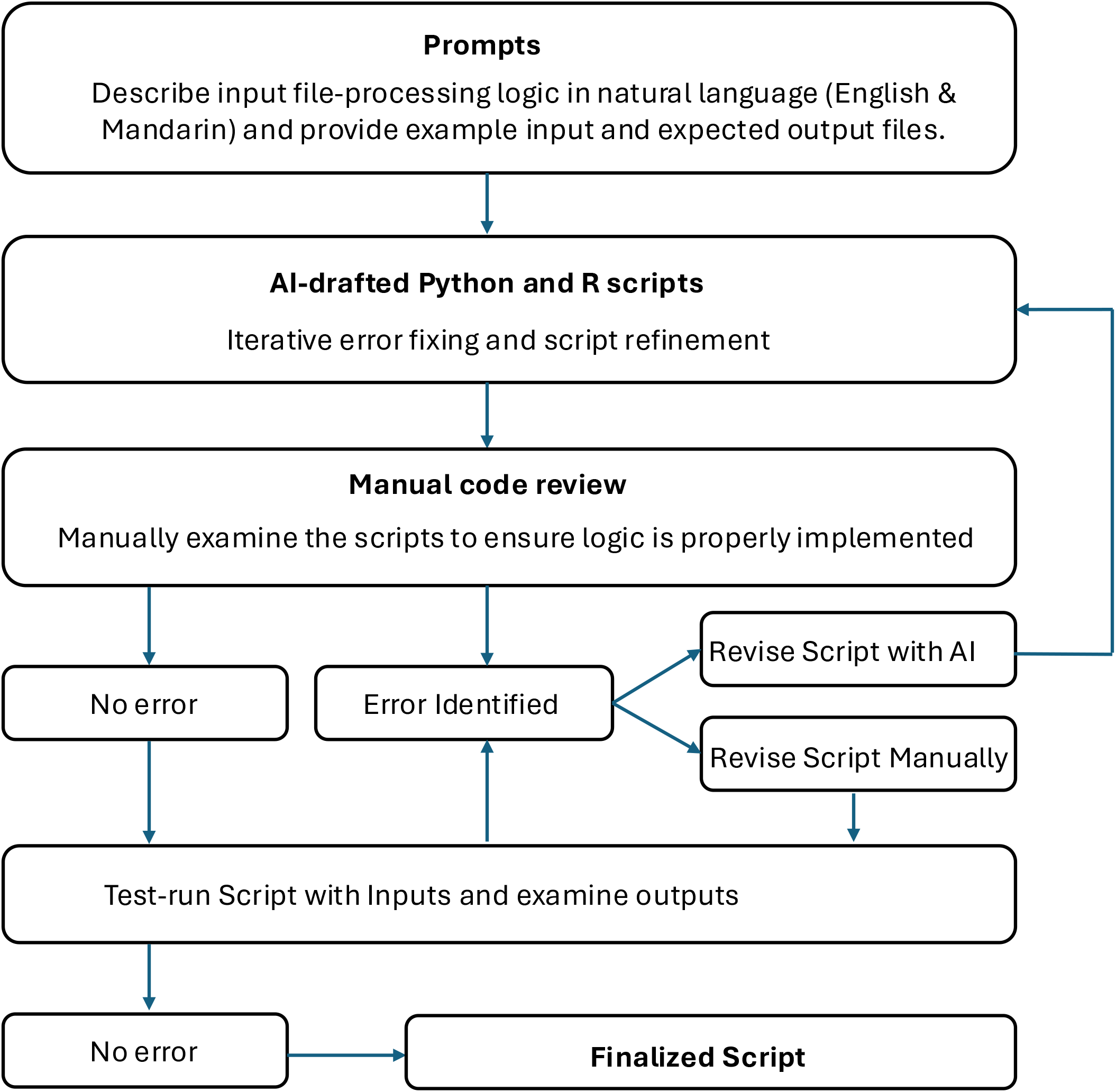
Workflow for using generative AI in script development. Natural-language prompts describing the desired file-processing logic (in English and Mandarin) and example input/expected output files were provided to a generative AI tool, which produced draft Python and R scripts. The authors then manually reviewed each script to verify that the intended logic was correctly implemented. If errors were identified, scripts were revised either with additional AI-assisted prompts or by manual editing, followed by repeated review. Scripts that passed manual inspection were test-run using example inputs and expected outputs. Only scripts that produced correct results in these tests were designated as finalized and used in the analyses.

**Supplementary Figure 25.**
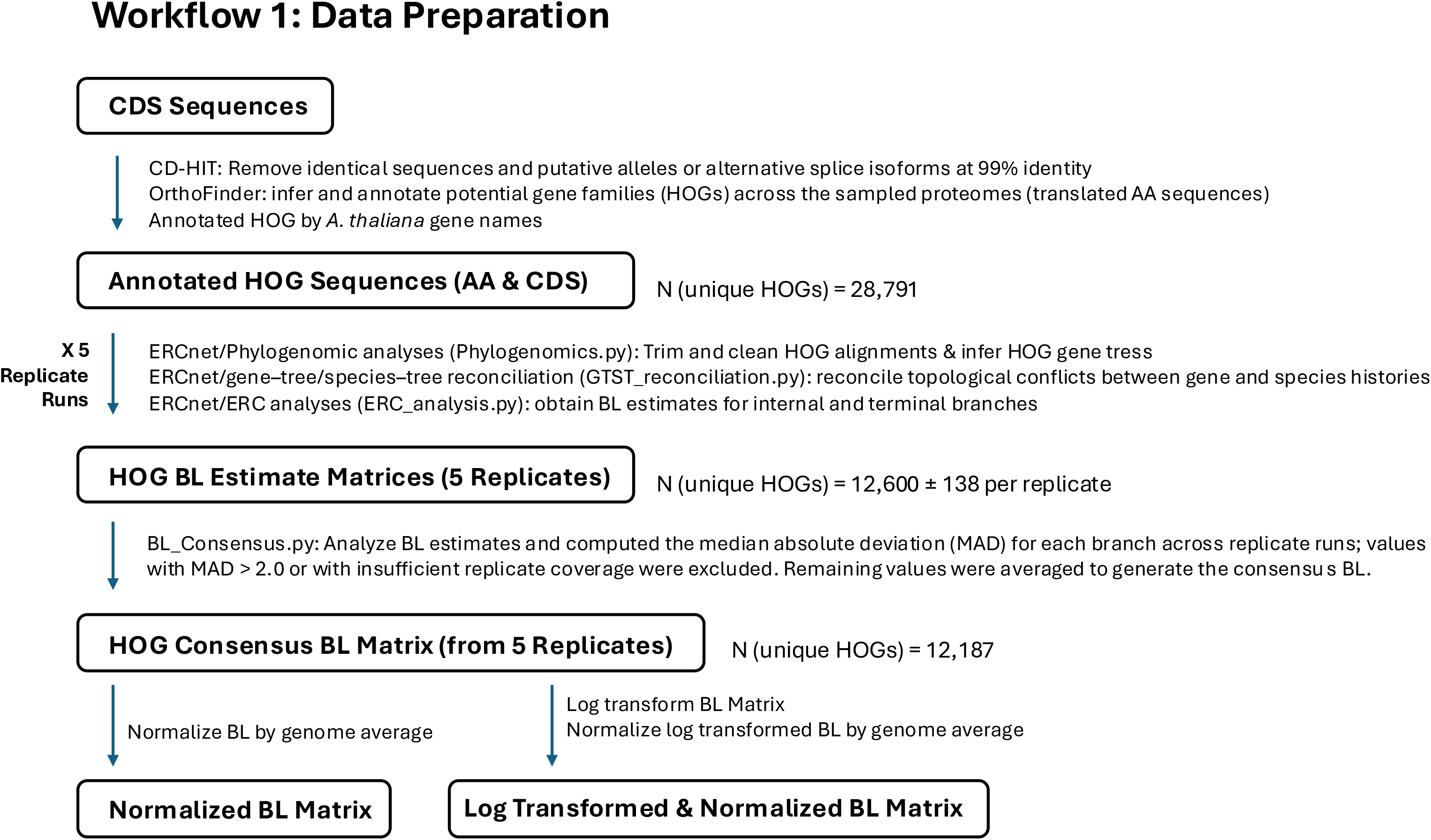
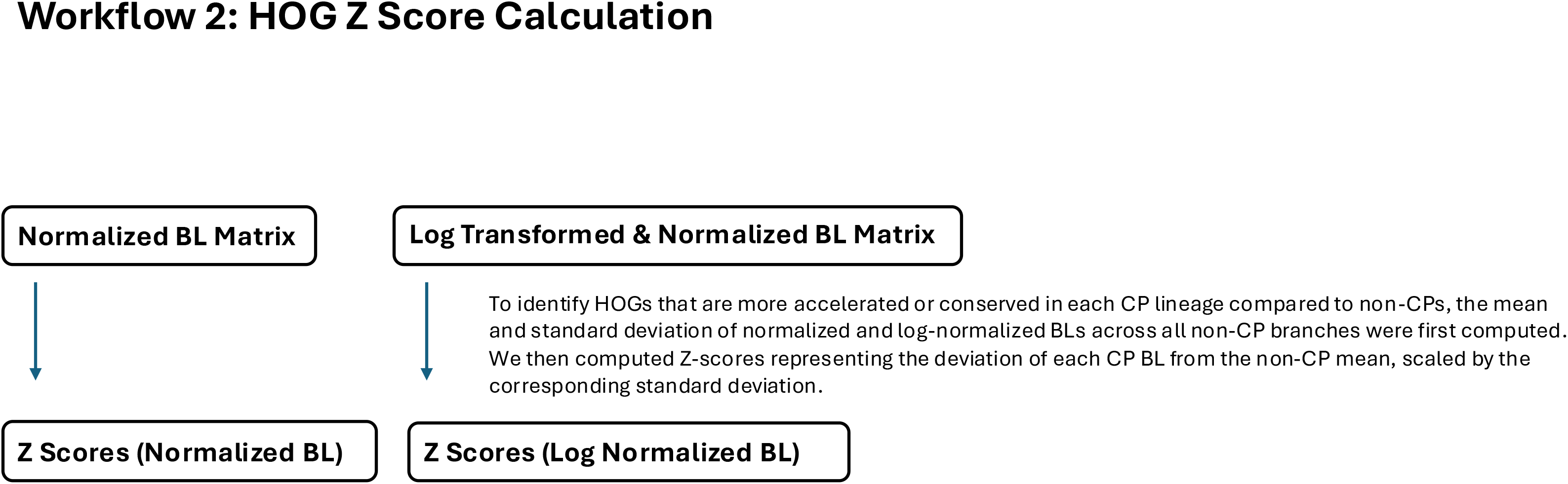
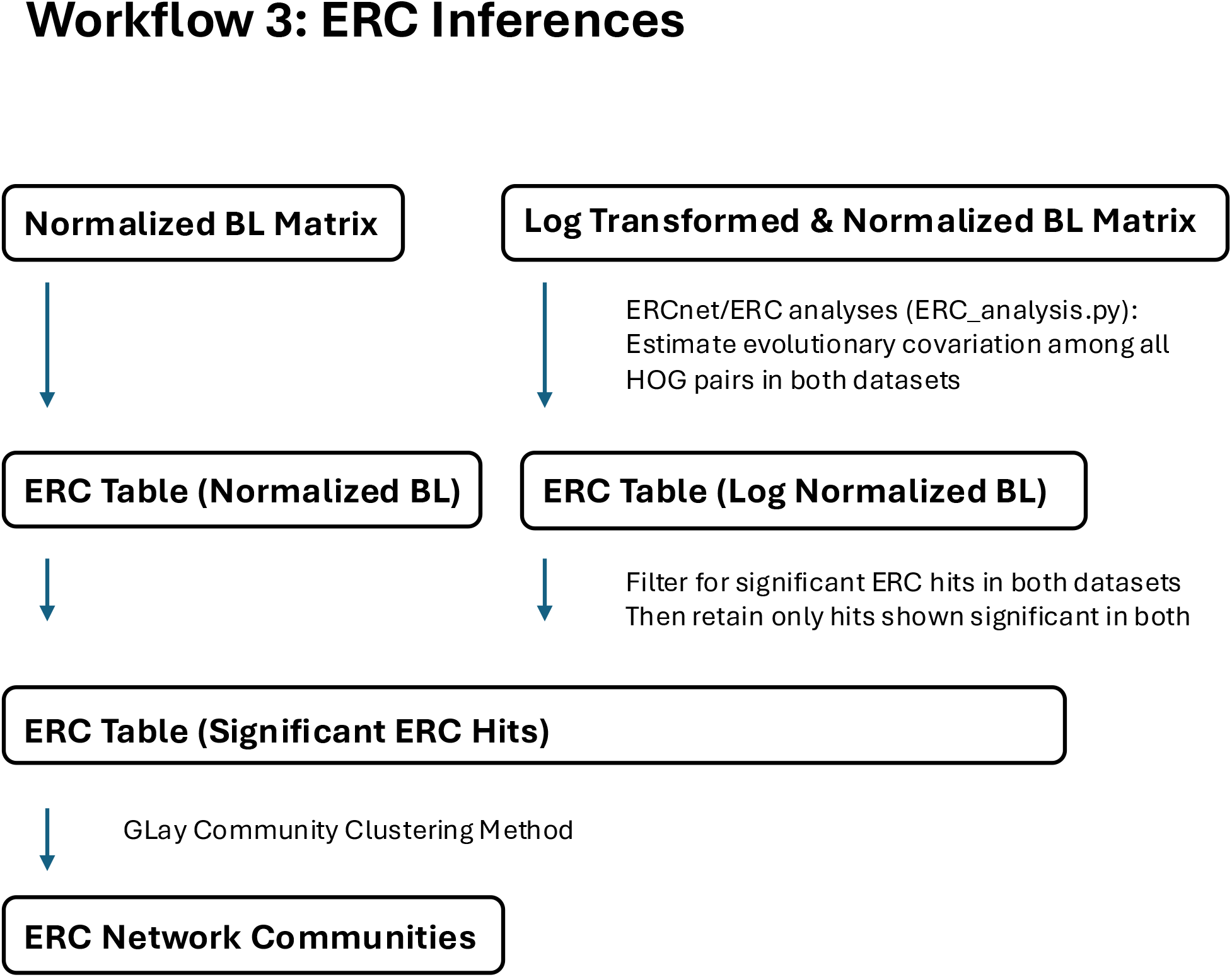
Overview of the analytical pipeline, from input sequences to ERC network communities. Schematic of the complete analytical workflow. Rounded boxes denote datasets (inputs, intermediate, and final outputs); arrows denote processing steps, with the associated software or custom script (italicized) and key parameters listed alongside. Abbreviations: CDS, coding sequence; AA, amino acid; HOG, hierarchical orthologous group; BL, branch length; MAD, median absolute deviation; CP, carnivorous plant; ERC, evolutionary rate covariation. **(A) Data preparation.** Input CDS sequences are de-duplicated with CD-HIT (99% identity, removing identical sequences and putative alleles/splice isoforms), and gene families (HOGs) are inferred from translated proteomes with OrthoFinder and annotated by *A. thaliana* gene names. Annotated HOGs are then processed through five replicate ERCnet runs—alignment trimming and gene-tree inference (*Phylogenomics.py*), gene-tree/species-tree reconciliation (*GTST_reconciliation.py*), and branch-length estimation for internal and terminal branches (*ERC_analysis.py*)—yielding five replicate BL matrices. *BL_Consensus.py* computes the MAD for each branch across replicates, excludes estimates with MAD > 2.0 or insufficient replicate coverage, and averages the remainder into a consensus BL matrix. The consensus matrix is normalized by the genome-wide per-branch mean to produce the Normalized BL Matrix, and log-transformed before normalization to produce the Log-Transformed & Normalized BL Matrix. **(B) HOG Z-score calculation.** For both normalized matrices, the mean and standard deviation of BLs across all non-CP branches are computed, and Z-scores representing the deviation of each CP branch length from the non-CP mean are derived, identifying HOGs with accelerated or conserved rates in each CP lineage. **(C) ERC inference.** ERC is estimated across all HOG pairs in both normalized datasets (*ERC_analysis.py*); pairs significant in both datasets are retained to form the final edge list, and GLay community clustering is applied to define ERC network communities.

**Supplementary Figure 26.**
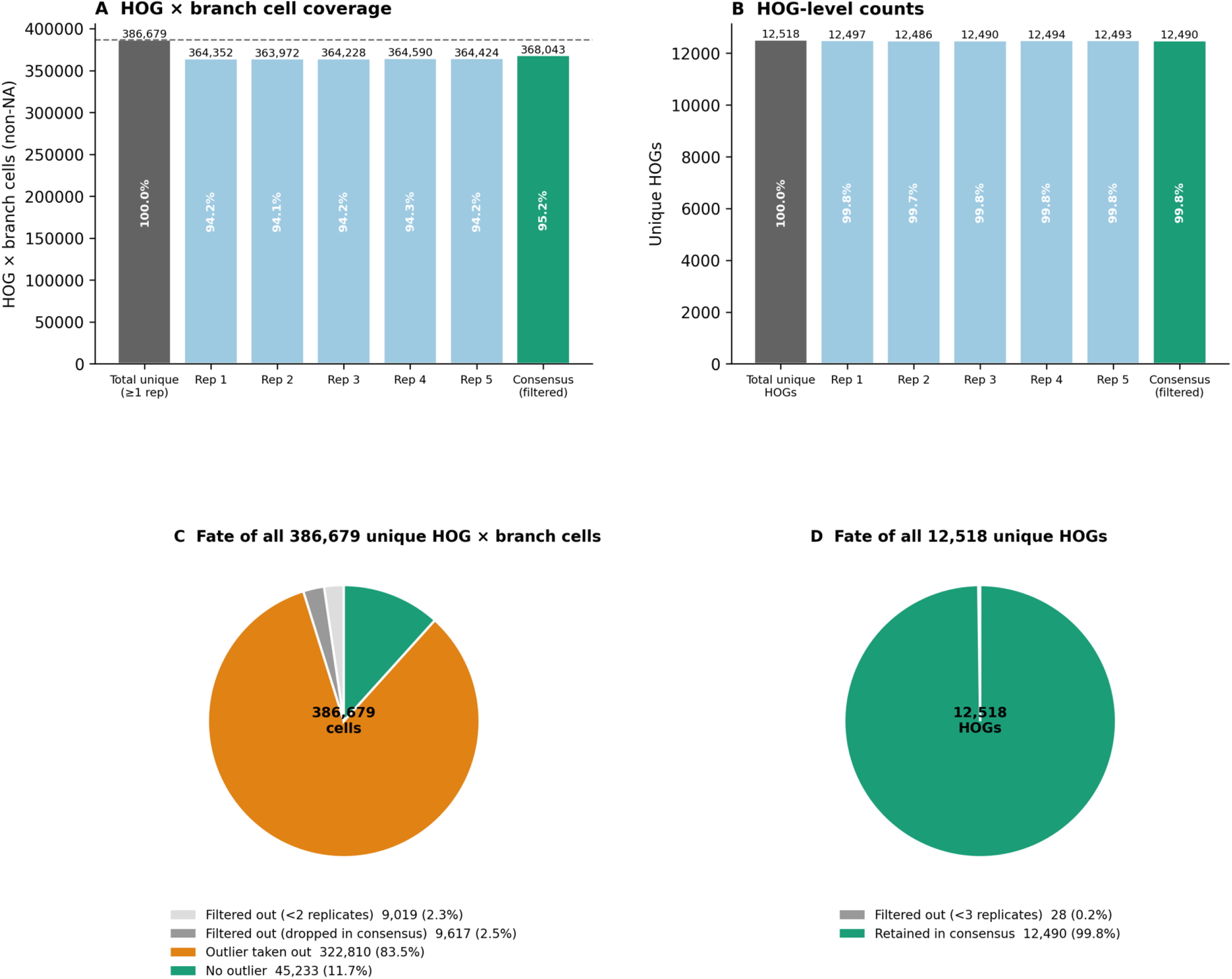
Branch-length (BL) estimates from five replicate ERCnet runs were combined into a consensus matrix. Each HOG × branch cell estimated in ≥3 of the 5 replicates was retained, and its consensus value computed as the mean of the replicate estimates falling within 2 × MAD of the cell median; individual outlier replicate values (>2 × MAD) were excluded from this average. **(A)** Coverage of the HOG × branch cell space: of 386,679 unique cells with an estimate in ≥1 replicate, each replicate recovered ∼94.2%, and 368,043 cells (95.2%) entered the consensus matrix. **(B)** The same stages at the level of unique HOGs: of 12,518 HOGs with any estimate, 12,490 (99.8%) are represented in the consensus. The count is nearly unchanged across stages because filtering operates *within* cells — removing individual outlier replicate values — rather than removing whole gene families. **(C)** Fate of all 386,679 unique HOG × branch cell estimates after consensus: 9,019 (2.3%) were present in only one replicate and 9,617 (2.5%) in only two (both too few to form a consensus and excluded), while of the classifiable cells (≥3 replicates), 322,810 (83.5%) were retained after dropping ≥1 outlier replicate value and 45,233 (11.7%) were retained with no outliers removed. **(D)** Fate of all 12,518 HOGs: only 28 (0.2%) were filtered out for insufficient replicate coverage (<3 replicates), and 12,490 (99.8%) were retained in the consensus. Together, panels C and D show that the consensus step discards individual outlier replicate estimates rather than whole cells or gene families, so nearly all HOGs are preserved — though most retained HOGs are represented by only a subset of their branches (Supplementary Figure 27).

**Supplementary Figure 27.**
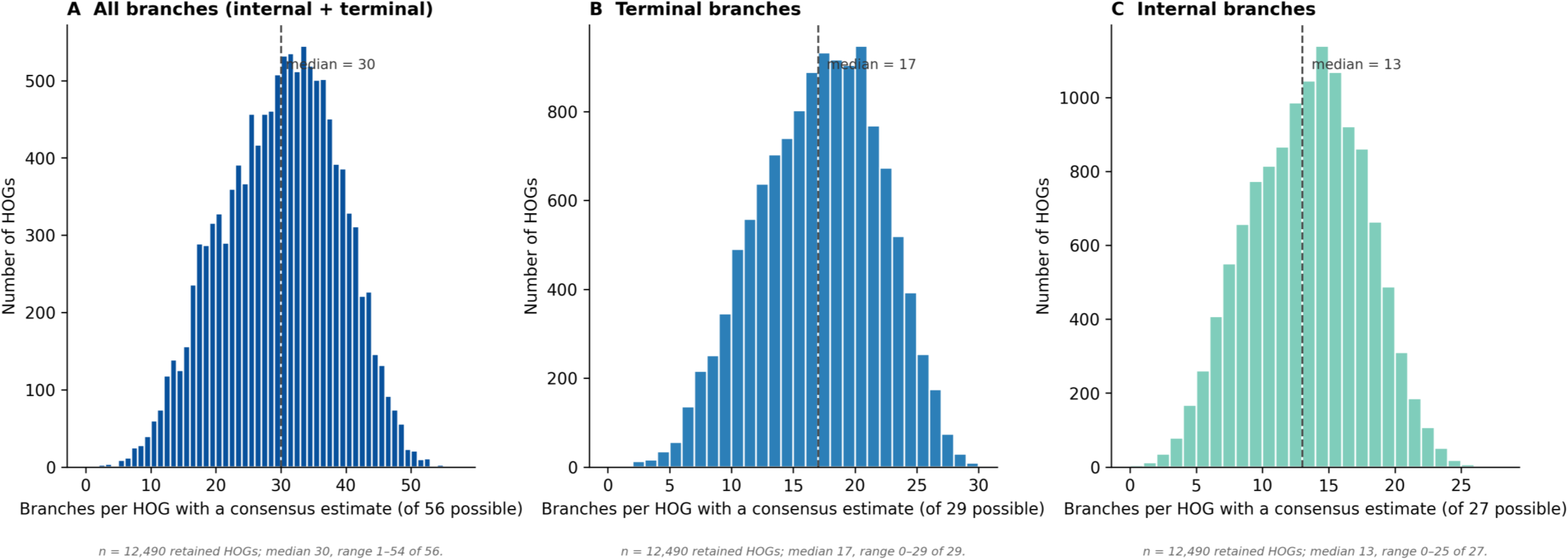
Number of branches with a consensus estimate per retained HOG, by branch type. For each of the 12,490 HOGs retained in the consensus branch-length matrix (Supplementary Figure 26D), the distribution of the number of branches with a consensus estimate (i.e., a branch estimated in ≥3 of the 5 replicates). **(A)** All branches (internal + terminal): median 30 of 56 possible branches (range 1–54). **(B)** Terminal branches only: median 17 of 29. **(C)** Internal branches only: median 13 of 27. Dashed lines indicate medians; a value of 0 in (B) or (C) denotes a retained HOG with no estimated branch of that type. Most retained HOGs are represented by a subset of the full branch set, reflecting that gene families are present in only a fraction of the sampled lineages (so many branches have no estimate to begin with) rather than removal of branches by the consensus step. Internal and terminal branches are retained in proportions broadly similar to their availability (27 and 29 possible, respectively), indicating that the consensus does not preferentially exclude either branch type.

**Supplementary Figure 28.**
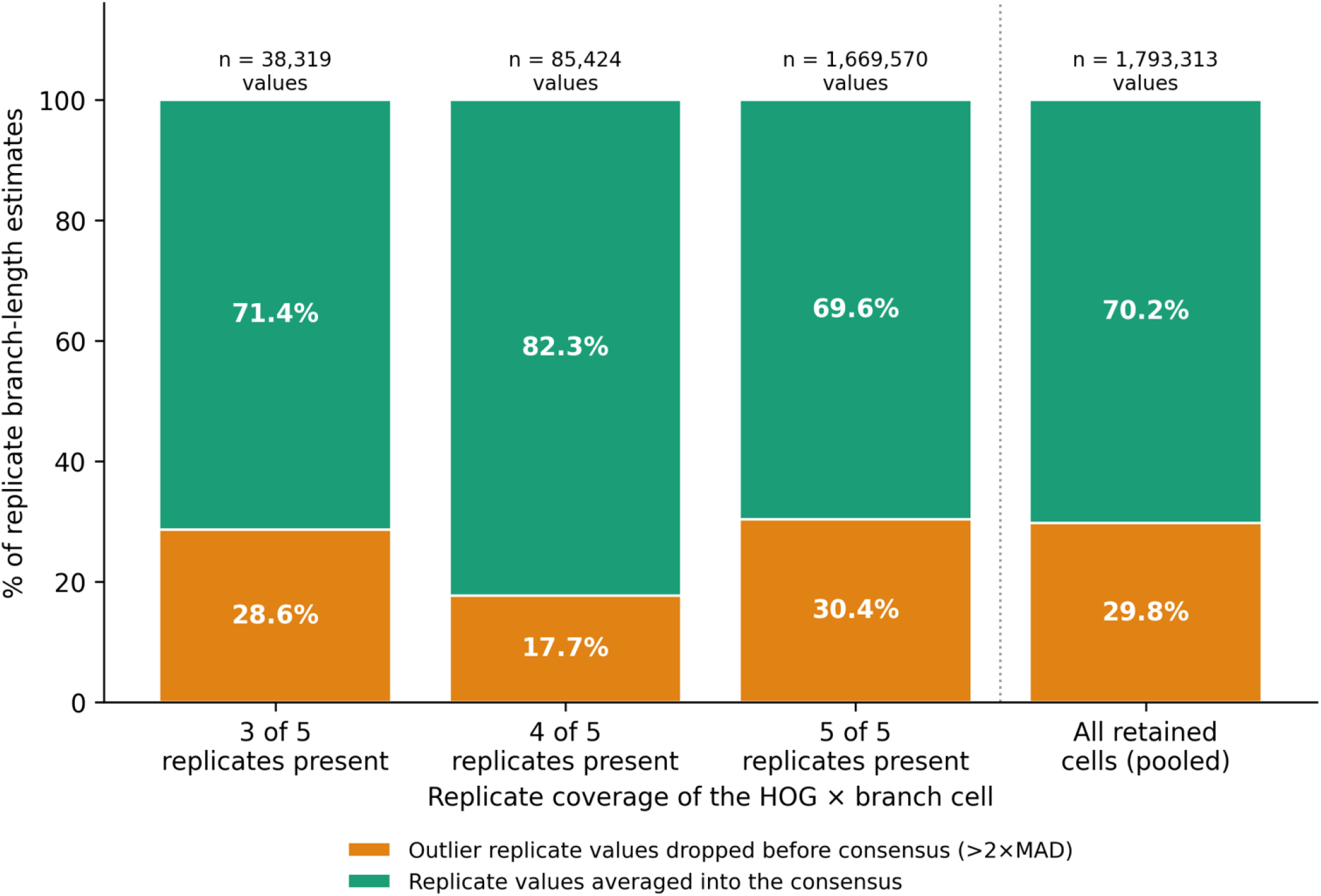
Outlier replicate values filtered out prior to consensus, by replicate coverage. For each classifiable HOG × branch cell (present in ≥3 of 5 replicates — all of which are retained), the proportion of individual replicate branch-length estimates that were averaged into the consensus versus excluded as outliers (deviating >2 × MAD from the cell median), stratified by the number of replicates in which the cell was estimated. Bars are normalized to 100%; *n* above each bar gives the number of replicate estimates, and the pooled bar (right) combines all coverage levels.

**Supplementary Figure 29.**
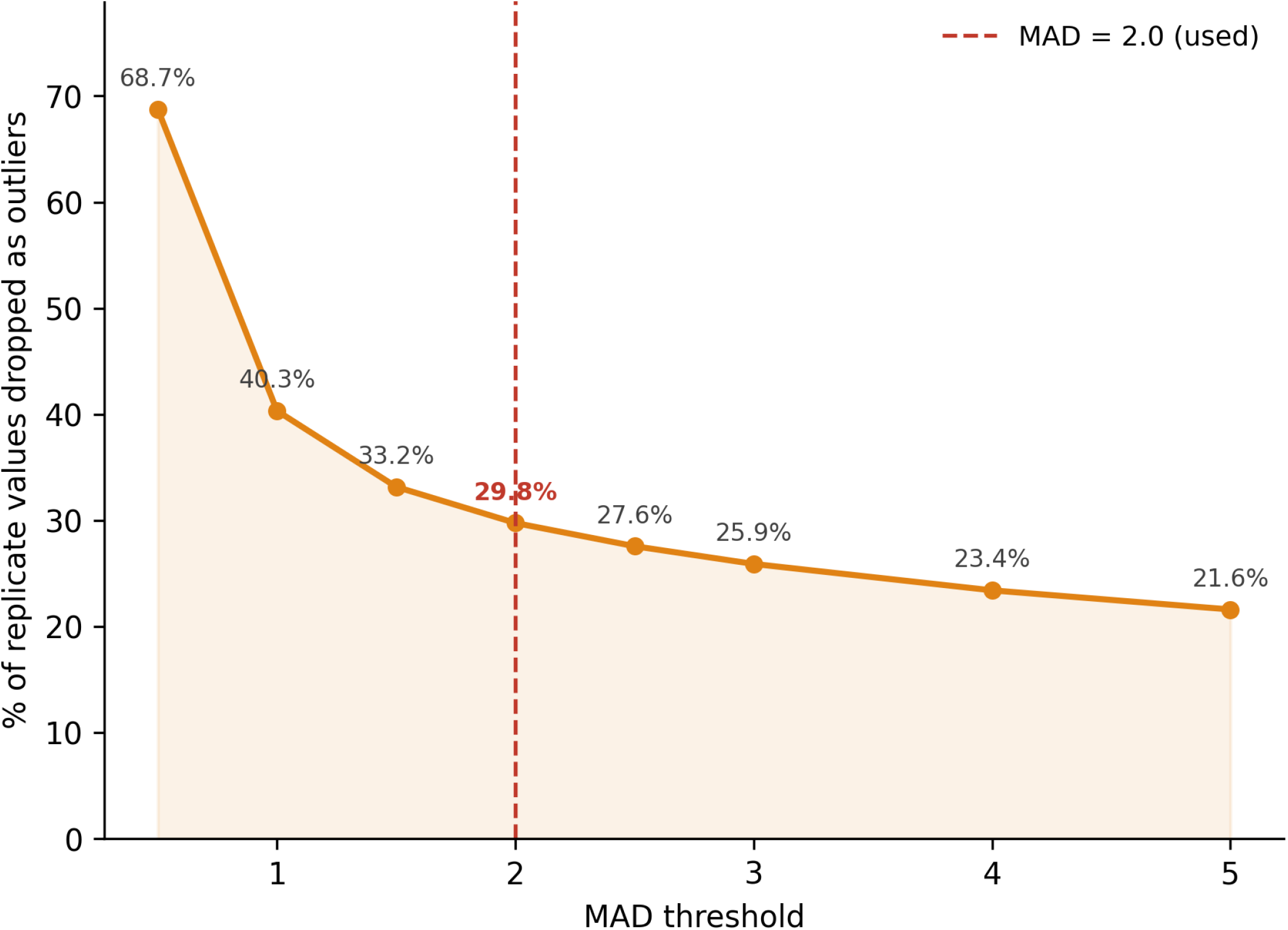
Sensitivity of outlier removal to the MAD threshold. Percentage of individual replicate branch-length estimates excluded as outliers as a function of the MAD threshold, across all classifiable HOG × branch cells (present in ≥3 of 5 replicates).

**Supplementary Figure 30.**
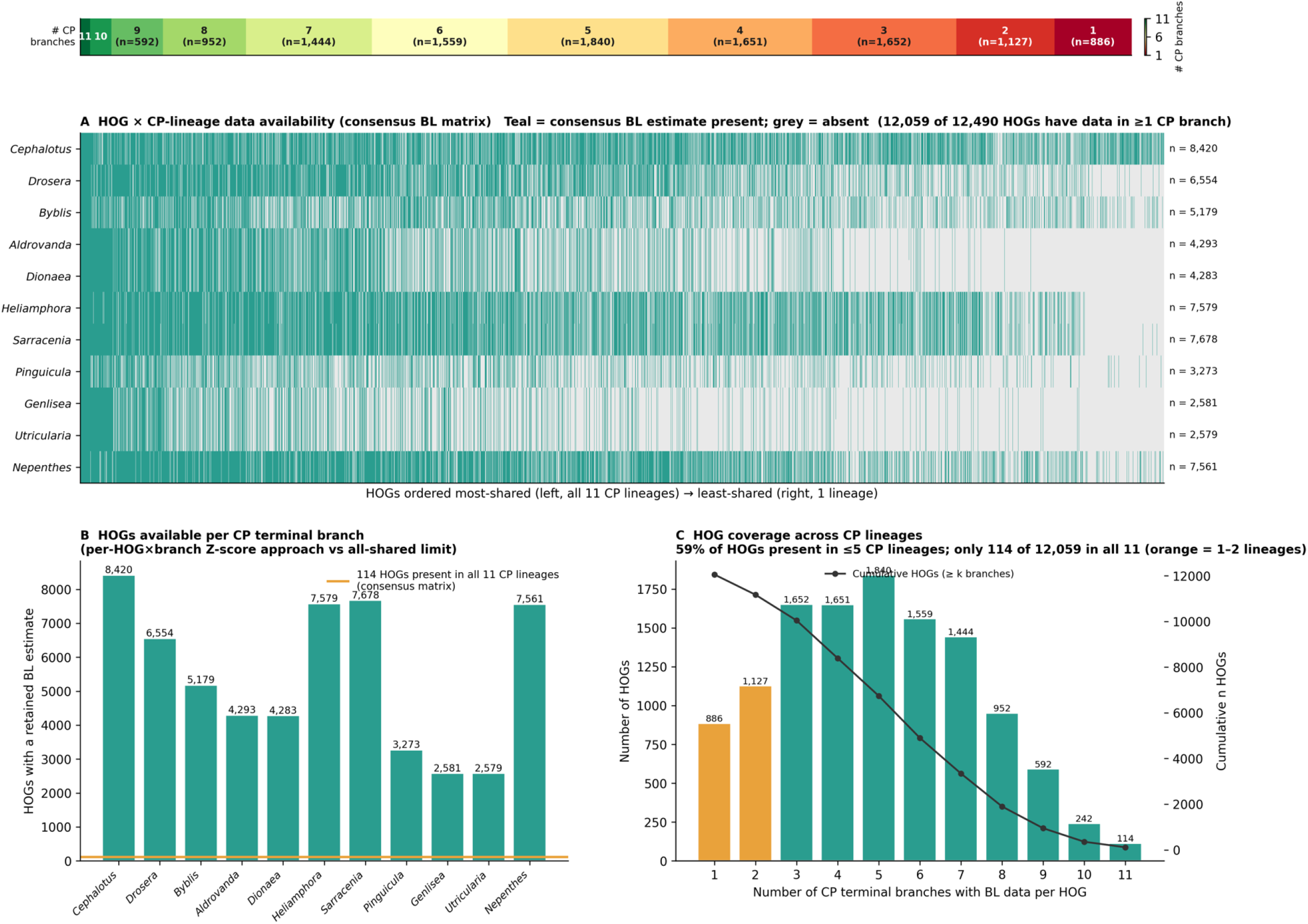
Per-lineage availability of branch-length estimates in the consensus matrix across carnivorous-plant lineages. Availability of consensus branch-length (BL) estimates across the 11 carnivorous-plant (CP) terminal lineages, for the classifiable consensus matrix used in all downstream rate analyses. Of 12,490 HOGs in the consensus matrix, 12,059 have a BL estimate on at least one CP terminal branch. **(A)** Availability matrix: each column is a HOG (ordered most-shared at left to least-shared at right) and each row a CP lineage; teal indicates a consensus BL estimate is present, grey its absence. The strip above colors each HOG by the number of CP lineages in which it is present (green = 11, red = 1; class sizes given as n). Per-lineage totals (n) are listed at right. **(B)** Number of HOGs with a consensus BL estimate on each CP terminal branch (2,579– 8,420); the orange line marks the 114 HOGs present on all 11 CP terminal branches simultaneously (the fully shared set, consistent with the four-way intersection in Supplementary Figure 1). **(C)** Distribution of the number of CP lineages each HOG is present in. Only 114 of 12,059 HOGs occur in all eleven lineages, and 59% are present in five or fewer (cumulative counts overlaid).

**Supplementary Figure 31.**
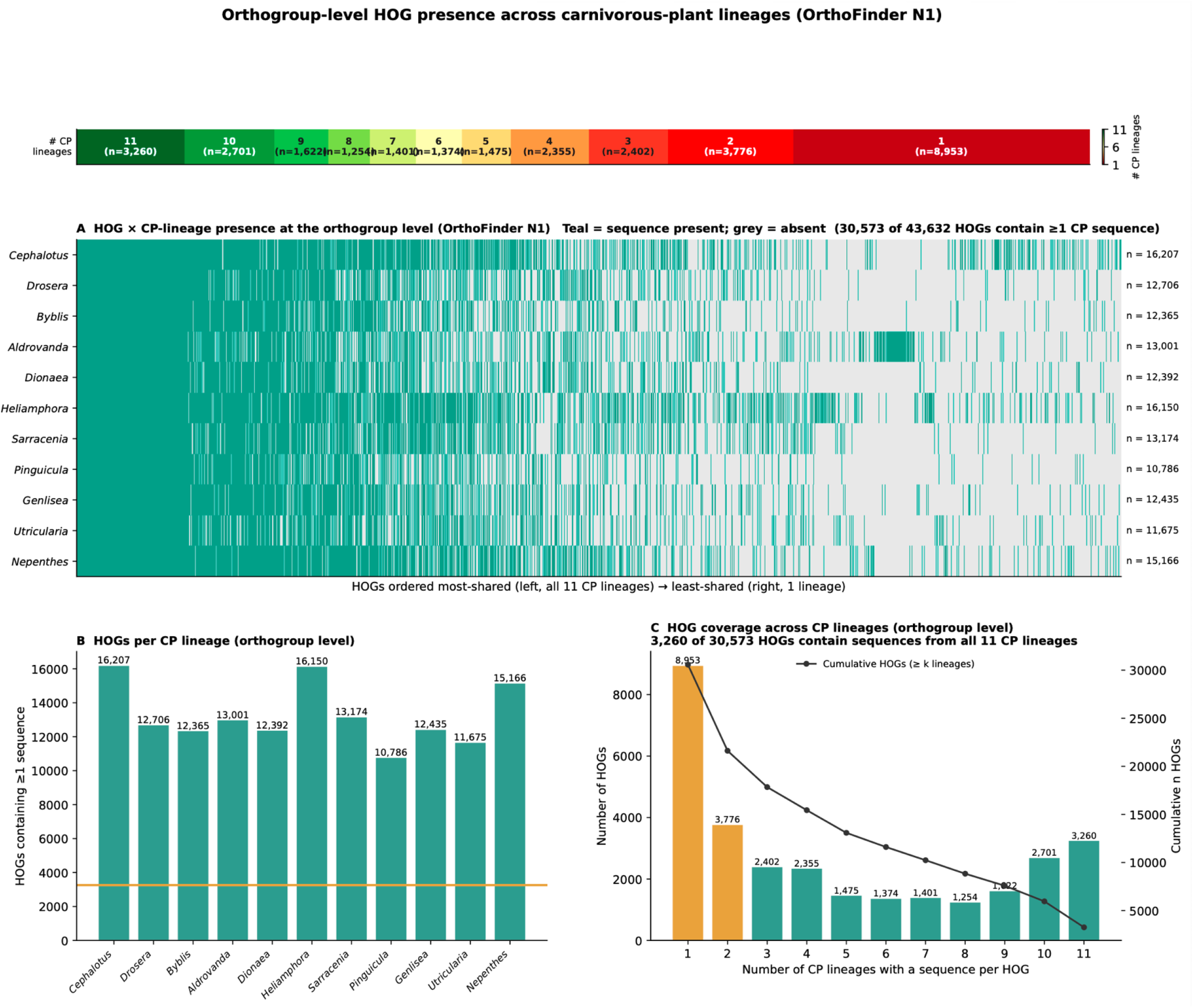
Orthogroup-level presence of HOGs across carnivorous-plant lineages (OrthoFinder N1). Presence of sequences from each of the 11 carnivorous-plant (CP) terminal lineages in the OrthoFinder hierarchical orthogroups (N1 level, the partition used by the ERCnet pipeline), before the ERCnet filters. Of 43,632 orthogroups, 30,573 contain at least one sequence from a CP lineage. **(A)** Presence matrix: each column is a HOG (ordered most-shared at left to least-shared at right) and each row a CP lineage; teal indicates ≥1 sequence from that lineage is present, grey its absence. The strip above colours each HOG by the number of CP lineages in which it is present (green = 11, red = 1; class sizes given as n). Per-lineage totals (n) are listed at right. **(B)** Number of HOGs containing sequences from each CP lineage (10,786–16,207); the orange line marks the 3,260 HOGs present in all 11 CP lineages. **(C)** Distribution of the number of CP lineages each HOG contains sequences from: 3,260 of 30,573 HOGs contain sequences from all eleven lineages (cumulative counts overlaid). In contrast to the far smaller number of HOGs that retain comparable branch-length estimates on all 11 lineages after the full pipeline (114; Supplementary Figures 1 and 30), thousands of orthogroups are shared at the sequence level — indicating that the sparsity of the consensus matrix’s cross-lineage intersection reflects downstream filtering and incomplete assemblies, not a lack of shared orthologs.

**Supplementary Figure 32.**
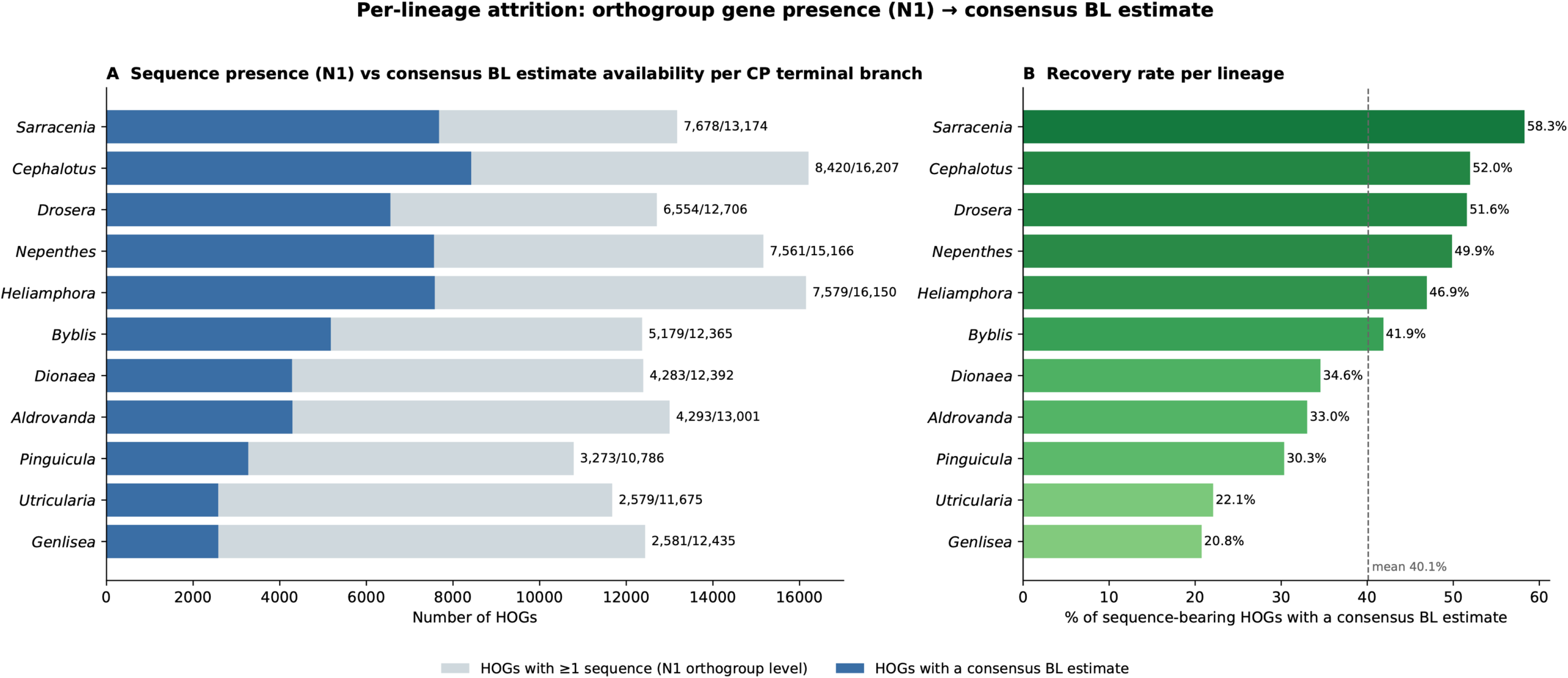
Per-lineage comparison of orthogroup gene presence and the availability of comparable branch-length (BL) estimates across CP lineages. For each of the 11 CP terminal branches, HOGs containing at least one sequence for that species in the OrthoFinder N1 table (grey) are compared with the subset of HOGs that yielded a consensus branch-length (BL) estimate on the same branch (blue). **(A)** Number of HOGs with ≥ 1 sequence (N1; grey) versus the number with a BL estimate (blue) per lineage; labels give the BL-estimate counts (BL/N1). **(B)** The corresponding recovery rate—the percentage of sequence-bearing HOGs that yielded a BL estimate at the consensus stage (dashed line, cross-lineage mean = 40.1 %).

